# When can we trust population trends? A method for quantifying the effects of sampling interval and duration

**DOI:** 10.1101/498170

**Authors:** Hannah Wauchope, Tatsuya Amano, William Sutherland, Alison Johnston

## Abstract

1. Species’ population trends are fundamental to conservation: they are used convey the global, national and local state of nature, justify calls to action and underpin many prioritisation exercises, including the IUCN red-list. It is crucial to be able quantify the degree to which population trend data can be trusted, yet there is not currently a straightforward means to do so.
2. We present a method that compares trends derived from various samples of ‘complete’ population time-series, to see how often these samples correctly estimate the direction and magnitude of the complete trend. We apply our method to a dataset of 29,226 waterbird population time-series from across North America.
3. Our analysis shows that if a significant trend is detected, even from only a few years, it is likely to reliably describe the direction (positive or negative) of the complete trend, though often does not approximate the magnitude of change well. However, if no significant trend is detected, a many-years long sample is required to be confident that the population is truly stable. Further, an insignificant trend is more likely to be missing a decline rather than an increase in the population. Sampling infrequently, but regularly, was surprising reliable in determining trend direction, but poor at determining the magnitude of change.
4. By providing percentage estimates of reliability for combinations of sampling regimes and lengths, we have a means to determine the reliability of species population trends. This will increase the rigor of large-scale population analyses by allowing users to remove time-series that do not meet a reliability cut-off, or weighting time series by reliability, and could also facilitate planning of future monitoring schemes. Our methods are applicable to other taxa and we provide the tools to do so.

## Introduction

Many crucial conservation decisions rely on knowing the overall trend of a species or population. This information underpins IUCN red-list classification (Rodrigues *et al.*, 2006), many national threatened species ranking systems (e.g. NESP Threatened Species Recovery Hub, 2018; U.S. Fish & Wildlife Service, 2018) and can convey to policy makers the state of nature globally, regionally and locally (Gärdenfors, 2001; Collen *et al.*, 2009). It is important that conservationists appreciate the extent to which they can trust the apparent trend of a population, both to ensure that at-risk species are not ignored and to avoid misallocating conservation resources towards species that are not actually at risk. The reliability of population trends are poorly understood, especially when data on variability and measurement error are not available. In addition, many large-scale analyses and policy recommendations (e.g. Collen *et al.*, 2009; WWF, 2016) rely on aggregating trends across numerous populations with little guidance on how to weight trends by their likely veracity.

Estimating the trend of a population requires a series of counts over time (typically years, as considered here). Linear, or non-linear, models are then fit to estimate yearly change (e.g. 1% decline per year), or modelled counts are compared between years at the start and end of a time period (e.g. a 10% increase over 10 years). The number of years of data, sampling frequency, degree of measurement error and population variability all affect the reliability of the derived trend. When data are available on measurement error and population variability, power analyses are recommended to estimate degree of reliability in trend estimates (Hayes & Steidl, 2002; Magurran *et al.*, 2010; Johnson *et al.*, 2014). Although power analyses are useful where sufficient data are available, there is often insufficient information, especially when assessing many populations, or using existing count data.

Previous studies have attempted to quantify reliability of trends using both simulated and real data. Simulated studies conclude that longer time scales are needed for better trend estimates, and that there are high margins of error when detecting small population declines (Wilson *et al.*, 2011; Prozt *et al.*, 2012; Tománková *et al.*, 2013; Connors *et al.*, 2014; Fox *et al.*, 2018). Studies working with real data on diverse taxa have found that populations exhibiting a particular trend across one time-interval often show an opposing trend in later years (Dunn, 2002; Keith *et al.*, 2015). The number of years of sampling required to reliably detect a trend has been estimated at both 10 (both by Rueda-Cediel *et al.*, 2015 for a snail species and White, 2018 for various vertebrates) and 21 years (Reynolds *et al.*, 2011 for brown bears). These investigations are useful for gaining an approximate idea of reliability, but do not provide a straightforward way for a study to assign a value of reliability to population time-series of varying lengths (i.e. numbers of years).

Therefore, in the absence of better guidance, studies based on population trends often lack the data to make any quantification of uncertainty (e.g. Craigie *et al.*, 2010, Loh *et al.*, 2005). Further, most studies assume that there is a ‘true’ trend exhibited by each population, but given most populations fluctuate over time in response to the positive and negative pressures affecting them, it is incorrect to maintain that any has one true detectable trend over long time-periods.

We propose a method to quantify uncertainty in trend estimates. Our analyses hinge on the concept of comparing the trend derived from a ‘sample’ (a subset of the full set of counts for a population) to the ‘complete’ trend of that population, derived from the full set of counts. We have chosen to use the word ‘complete’ in this study rather than ‘true’ as even with yearly counts we cannot claim to know the true trend of a population. Normally, one would possess only the sample, and we therefore hope to provide an estimate of how likely that sample is to represent the complete trend, regardless of sample length or complete trend length. In our analysis we quantify reliability both in terms of accuracy of trend direction and magnitude of change.

We ask two questions: 1) How reliable are trends derived from a certain number of years of data, based on the time over which a trend estimate is desired? For example, how well do 5 consecutive years of survey data represent the trend of a population over 10 years?; and 2) How reliable are trends derived from data sampled at different intervals, such as samples taken every year over a 30-year period compared to every 5 years over the same period? We also investigate two factors that we expect to impact reliability: species generation time and shape of the complete trend. We expect that species with longer generation times will require longer survey periods, as there will be a lag before populations show responses to changes in birth rate while older individuals are still living (Kuussaari *et al.*, 2009). We also expect that trends estimated from samples of populations with complex non-linear complete trends will be less accurate than samples from populations with linear or near-linear complete trends.

As a case study, we use an empirical dataset of yearly counts of 129 waterbird species at 1,110 sites in North America (a total of 29,226 site by species combinations). Providing these estimates for waterbird data is particularly beneficial as data on waterbirds are available at large spatio-temporal scales and waterbird studies can provide insights into broader conservation goals (Piersma & Lindström, 2004; Amat & Green, 2010; Amano *et al.*, 2018). However, our methods are general, and we provide code and instructions to generalise to other taxa. Our work provides an explicit measure of the reliability of a trend and gives an evidence-based justification for excluding samples below a certain length, according to the degree of confidence desired for a study. Finally, these results can be used to plan multi-species monitoring programs, to give the highest likelihood of capturing representative trends for the most species.

## Methods

### Data

We obtained an initial dataset of yearly count data for 174 waterbird species in North America from the Christmas Bird Count (CBC; Dunn *et al.*, 2005) at 1,123 sites spanning the years 1966 to 2013 (Amano *et al.*, 2018), from which 30 years of consecutive counts were taken for each site by species combination. We selected 30 years because it represented a good balance between a long-term survey period, yet one for which adequate data were still available. In cases where a site was sampled for over 30 years, the most recent 30 years were taken. The supporting material provides a list of species and a map of spatial coverage. The CBC has been conducted yearly since 1900, primarily in the US and Canada, and involves a count of all bird individuals within a ‘site’, defined as a circle 24.1km in diameter. Sites are accessed by a variety of means including foot, car, boat, and snow mobile. Effort varies between sites, but the number of survey hours spent per count is documented, allowing effort to be accounted for (see below). We considered each species at each site as an independent population; as we were not attempting to estimate the trends of entire species, correlations between sites were irrelevant. This gave us an initial dataset of 56,022 populations.

### Data preparation

Christmas Bird Count data have variable sampling effort and this must be accounted for in the modelling process. The most common expected relationship between effort and detection is a linear relationship between log-transformed count and effort. Following Butcher & McCulloch (1988) and Xu *et al.* (2015), we chose to retain only those species where a significant linear relationship between detection and log of effort was shown, found by running a negative binomial generalized linear model (see Modelling Specifications, below) for each species, at all years and sites:

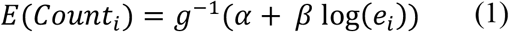

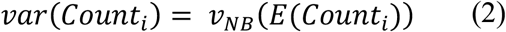

The link function g(·) is ‘log’, so the inverse is an exponential. The expected value of Count for species *i* is predicted by an intercept, α, the log of effort (in hours), *e* (Eq. 1) and it’s coefficient, *β*. The variance of our count data is defined as negative binomial (Eq. 2). Any species found to have a non-significant *β* were removed from analysis, as were those with a significant, but negative *β* (i.e. as effort increased, detection decreased). We then included survey hours as an offset term in our models to account for this sampling effort.

We also removed any populations with a sum of less than 30 observations over the 30-year sampling period, to remove populations with mostly zero counts. This resulted in our final dataset of 29,226 populations, comprising 129 species at 1,110 sites.

As species varied in the extent to which they occurred at sites, we also ran our analysis on a standardised subset of the data: 99 species with 50 randomly selected sites each, 4,950 populations in total. Even though this dataset was less than 20% of the size of our full dataset, the results were highly congruent.

### Modelling Specifications

To estimate the population growth rate, *r*, with population counts as the response variable and years as the explanatory variable, we used Generalized Linear Models (GLMs) run with the R package stats (R Core Team, 2017). We included effort using the ‘offset’ parameter, which allows a covariate with a known slope to be included in the model. For count data it is usual to use Poisson, quasi-Poisson or negative binomial distributions for the response, with the latter two being more appropriate if there is over-dispersion, where the variance of the response variable exceeds the mean. In our dataset 99.7% of samples were over-dispersed, with 77% of these by at least an order of magnitude. We therefore ran our models using the negative-binomial distribution, though our provided code allows specification of any of these three distributions.

Mathematically, the above is expressed as the following:

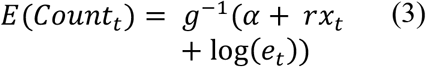

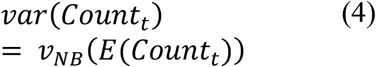

As before, the link function g(·) was ‘log’, so the inverse is an exponential. The expected value of Count in year t is predicted by an intercept, α, the population growth rate, *r*, multiplied by the year value, *x*, and the log of effort (in hours), *e* (Eq. 3). Because the relationship between effort and count is known (i.e. a log linear relationship), it does not need a coefficient. Also, as before, the variance of our count data is defined as negative binomial (Eq. 4).

For each model the population growth rate, *r*, and p-value of *r* were determined. For our main analysis, we followed the convention of setting a significance level of p<0.05. This is an arbitrary threshold, and circumstances may arise where the risk of missing a trend is greater than the risk of erroneously concluding there is one (e.g. a high risk group of species), in which case it is better to set a higher p-value (Taylor & Gerrodette, 1993; Field *et al.*, 2007), and *vice versa*. We provide functionality to adjust p-values in the our code.

All models were run in R version 3.4.1 (R Core Team, 2017) using the Cambridge Service for Data Driven Discovery High Performance Computing service (https://www.hpc.cam.ac.uk, last visited 14th Dec, 2018).

### Sampling Methods (Consecutive and Interval)

We considered two ways in which sampling could be conducted, and took subsamples of the complete data according to these. First, we used Consecutive Sampling (Figure 1a), i.e. sampling from consecutive years, where shorter adjacent subsets were taken from a complete dataset of n years in length. We sampled all possible consecutive subsamples from three years to n-1 years. Second, we used Interval Sampling, where samples were taken at regular intervals from within the complete 30 year dataset (Figure 1b): we varied the interval length (i.e. samples taken every x years) from 1 year (i.e. consecutive years) to 14 years (i.e. samples taken at years 1, 15 and 29, or 2, 16 and 30 years) and took all possible numbers of samples within these iterations to fill the 30-year period. Table 1 gives examples of these.

**Figure 1.**
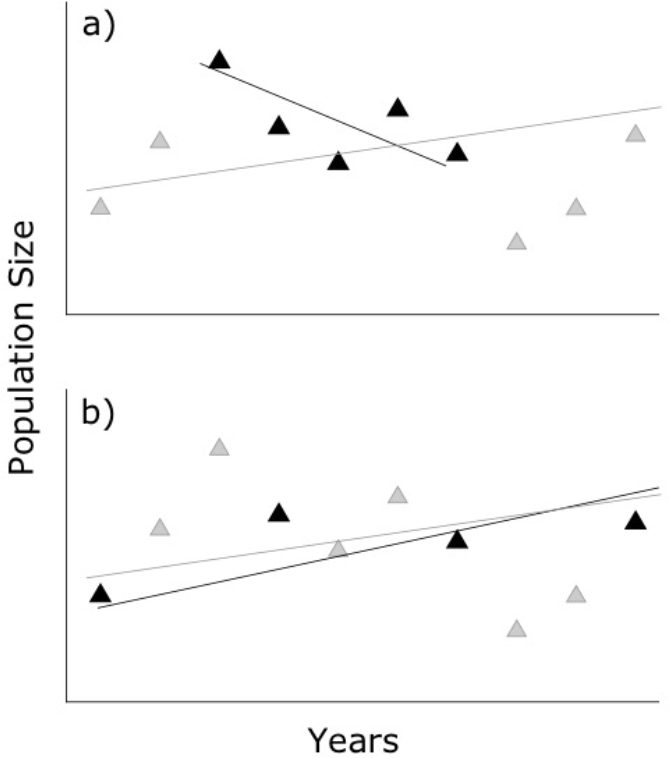
Schematic diagram showing sampling and modelling technique for comparing sample trends and complete trends. Triangles show counts of a population, all counts are used to model the complete population trend (grey line), and only black triangles are used to model the sample trend (black line). Shows examples of: (a) Consecutive Sampling of five-years from a ten-year period; and (b) Interval Sampling, where samples were taken every 3 years from a ten-year period, giving four samples in total. We ran analysis on all possible Consecutive lengths, Interval lengths and complete trend lengths.

The complete trend was initially estimated from annual surveys over the full 30-year time-period. We also conducted our Consecutive Sampling with differing lengths of ‘complete years’. For example, we defined the complete trend as the trend a population exhibited over a 10-year period. To do this, for each of our 29,226 populations, we took all possible 10 year segments (e.g. 1-10, 2-11, 3-12 etc), and calculated the trend on all possible consecutive subsets of those ten years, which ranged from 3 to 9 years in length. We then compared these to trends calculated from the complete 10 years. We repeated this for Complete trend lengths ranging from 4 years to the full 30 years.

Because Interval Sampling already had two dimensions (interval length and number of years sampled), we compared this only to the complete trend length of 30 years.

**Table 1.** All iterations used for interval samples taken every 8 years (i.e. interval length = 8) and number of years sampled as 3 or 4. For shorter interval lengths, larger subset sizes were possible; at the shortest interval of 1, i.e. samples taken every year, up to 29 consecutive years could be sampled.

| 3 years sampled | 4 years sampled |
| --- | --- |
| 1, 9, 17 | 1, 9, 17, 25 |
| 2, 10, 18 | 2, 10, 18, 26 |
| 3, 11, 19 | 3, 11, 19, 27 |
| 4, 12, 20 | 4, 12, 20, 28 |
| 5, 13, 21 | 5, 13, 21, 29 |
| 6, 14, 22 | 6, 14, 22, 30 |
| 7, 15, 23 |  |
| 8, 16, 24 |  |
| 9, 17, 25 |  |
| 10, 18, 26 |  |
| 11, 19, 27 |  |
| 12, 20, 28 |  |
| 13, 21, 29 |  |
| 14, 22, 30 |  |

### Comparison Methods (Direction and Magnitude)

We used two ways to assess whether a sample trend (Consecutive or Interval) was representative of a complete trend. First, we took the ‘direction’ of the trend, defining it as positive, negative or insignificant. Using this, a sample trend would be classified as matching if it was the same direction as the complete trend; opposing if it was the opposite direction; an erroneous positive or negative if it was positive or negative, but the complete trend was insignificant; and a missed positive or negative if it was insignificant, but the complete trend was positive or negative (Table 2). We term this ‘Direction Comparison’. Note that we conducted a final supplementary analysis considering how often insignificant sample trends still approximated the direction of significant complete trends (see Supporting Information Section 4).

**Table 2.** Categories used for Direction Comparison. Columns show direction of trend derived from complete time-series and rows show direction of trend derived from sample time-series. Cells show category, based on sample and complete trend direction.

|  |  | Complete Trend Direction |  |  |
| --- | --- | --- | --- | --- |
|  |  | Positive | Negative | Insignificant |
| Sample Trend Direction | Positive | Matching | Opposing | Erroneous Positive |
|  | Negative | Opposing | Matching | Erroneous Negative |
|  | Insignificant | Missed Positive | Missed Negative | Matching |

Second, for cases where a significant trend was obtained from *both* the sample and complete time-series (i.e. cases of ‘Matching’ or ‘Opposing’ from the Direction Comparison method), we considered the absolute difference between population growth rate *r* (i.e. the slope of a log-linear model) of the two; giving an idea of the degree of ‘correctness’. That is, difference = |*r*_sample_ − *r*_complete_|. We defined tolerance levels ranging from ±0.01 to ±0.5 and if the difference was less than the tolerance level, the sample trend represented the complete trend and was correct, and was incorrect if it did not. We term this ‘Magnitude Comparison’ (Fig. 2).

**Figure 2.**
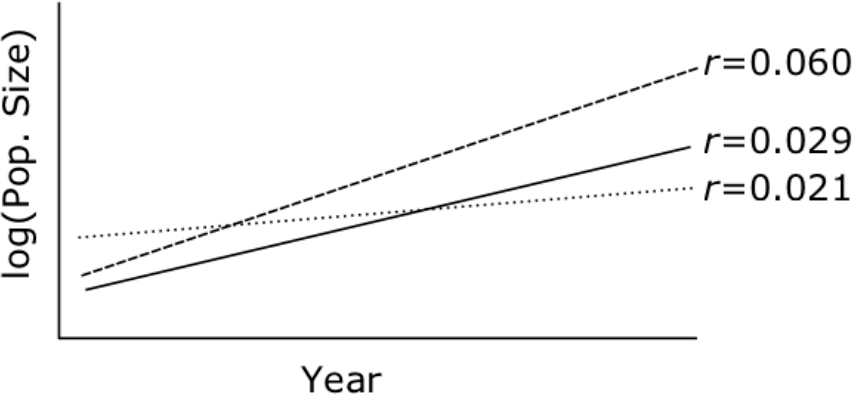
Example of Magnitude Comparison, showing three trends taken from the time-series of a particular population with modelled log population size vs. time. The solid line is the trend derived from the complete time-series, and the dotted and dashed lines are trends derived from samples of the complete time-series. The sample trend shown by the dotted line is correct at a tolerance of ±0.01 (because |0.021-0.029| < 0.01, but the sample trend shown by the dashed line is not (because |0.021-0.060| > 0.01).

In all cases, we obtained a sample *r* and a complete *r* for each population, the significance level of each, and then compared them to give a category for representativeness (using either the Direction or Magnitude Comparison method). We then found, for each combination of sample and complete time-series lengths and sampling types, the percentage of our 29,226 populations in each representativeness category.

### Generation Length

We considered generation length as a major factor that is likely to influence the duration of sampling required. This is because long-lived species often take longer to show responses to environmental pressures, as older individuals can continue to survive even if recruitment is falling (Kuussaari *et al.*, 2009). To assess this, we divided our species into three groups based on generation length: short (1-5 years), medium (6-10 years) and long (11-15 years). Generation length data was obtained from birdlife.org species fact sheets (e.g. http://datazone.birdlife.org/species/factsheet/ruddy-turnstone-arenaria-interpres/details, accessed 26^th^ July 2018). We then organised our standard analysis according to these three categories.

### Trend Shape

To assess how our results are affected by trends of different shapes, we used Generalized Additive Models (GAMs) with the R package MASS (Venables & Ripley, 2002). These non-parametric models allow for non-linear relationships. We ran GAMs on all complete 30 year trends, model specification was the same as the GLMs but with a smoothing term on year, and took the estimated degrees of freedom (EDF) for each. EDFs ranged from 1 to 8.57, so we divided our trends into four shape groups, linear and quadratic up to cubic (EDF = 1 − 2.99), cubic or low degree polynomial (EDF = 3 − 4.99), mid-degree polynomial (EDF = 5 − 6.99) and high degree polynomial (EDF = 7 − 8.99). We then organised our standard analysis (using GLMs) according to these four categories, to see whether sample trends were more of less representative when sample data was taken from complete data of different shapes.

## Results

See supporting information Section 1 for summary statistics of the full dataset. All data used to produce plots are also provided, as well as code to reproduce full results on any set of population counts.

### Direction Comparison of Consecutive Sampling

Reliable estimation of the direction of complete trends required many years of data for Consecutive Sampling. For example, to have an 80–100% chance of a sample trend having the same direction as a complete trend, the sample time-series needed to be almost as long as the complete time-series (Fig. 3a). However, sample trends opposed the complete trend less than 10% of the time (Fig 3b).

**Figure 3.**
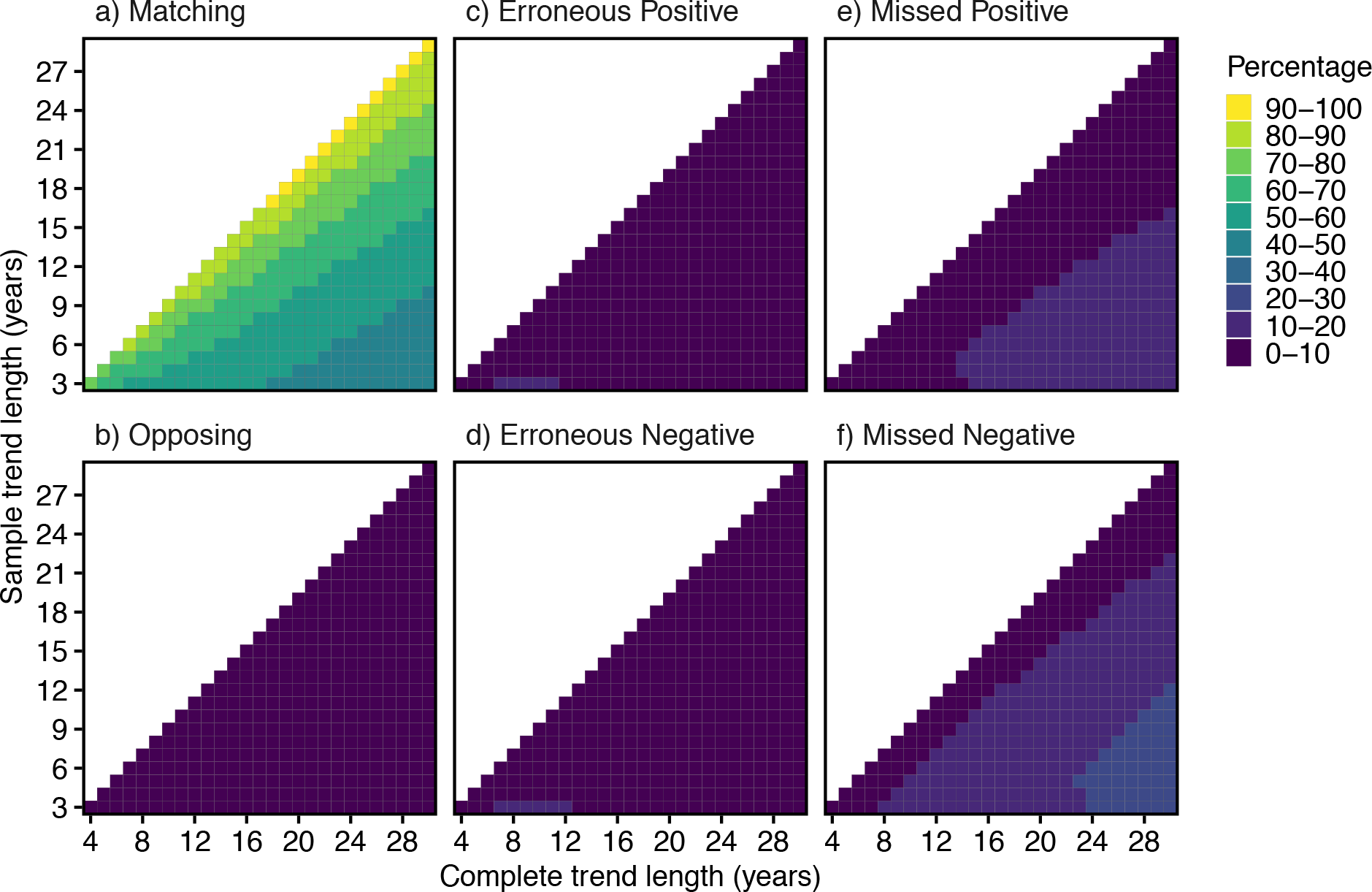
Direction Comparison of sample population trends using Consecutive Sampling. Colour shows percentage of sample trends that, relative to the complete trend, were matching, opposing, an erroneous positive/negative or a missed positive/negative (see Table 1). Shown for all combinations of sample lengths (y-axis), and complete lengths (x-axis).

The chance of an erroneous positive or negative (i.e. the sample indicated a significant trend but the complete trend did not reflect this, Fig 3c, d) was low regardless of the length of the sample or complete trend. However, the chances of a missed positive or negative trend (i.e. the complete trend had a significant direction but the sample did not detect this) were higher (see also Supporting Information Section 4); missed negatives were more likely than missed positives, and both were more likely when the sample time-series was considerably shorter than the complete time-series (Fig 3e, f). This implies that, particularly when trying to detect declines, shorter samples have low power to detect complete trends, but if they do detect a significant trend it is likely to be representative.

### Direction Comparison of Interval Sampling

Our results show that sampling in intervals can be more representative than sampling in consecutive years when considering trend direction. For example, sampling for 26 consecutive years (out of a 30-year complete time-series) gave the matching result 80% of the time (Fig. 4a, bottom row, note this is equal to Fig. 3a rightmost column), but the same level of reliability could be obtained by sampling 13 times every second year (Fig. 4a, second row). More strikingly, 4 years taken every 9 years gave the same percentage matching (60-70%) as up to 20 years of Consecutive Sampling (Fig. 4a).

**Figure 4.**
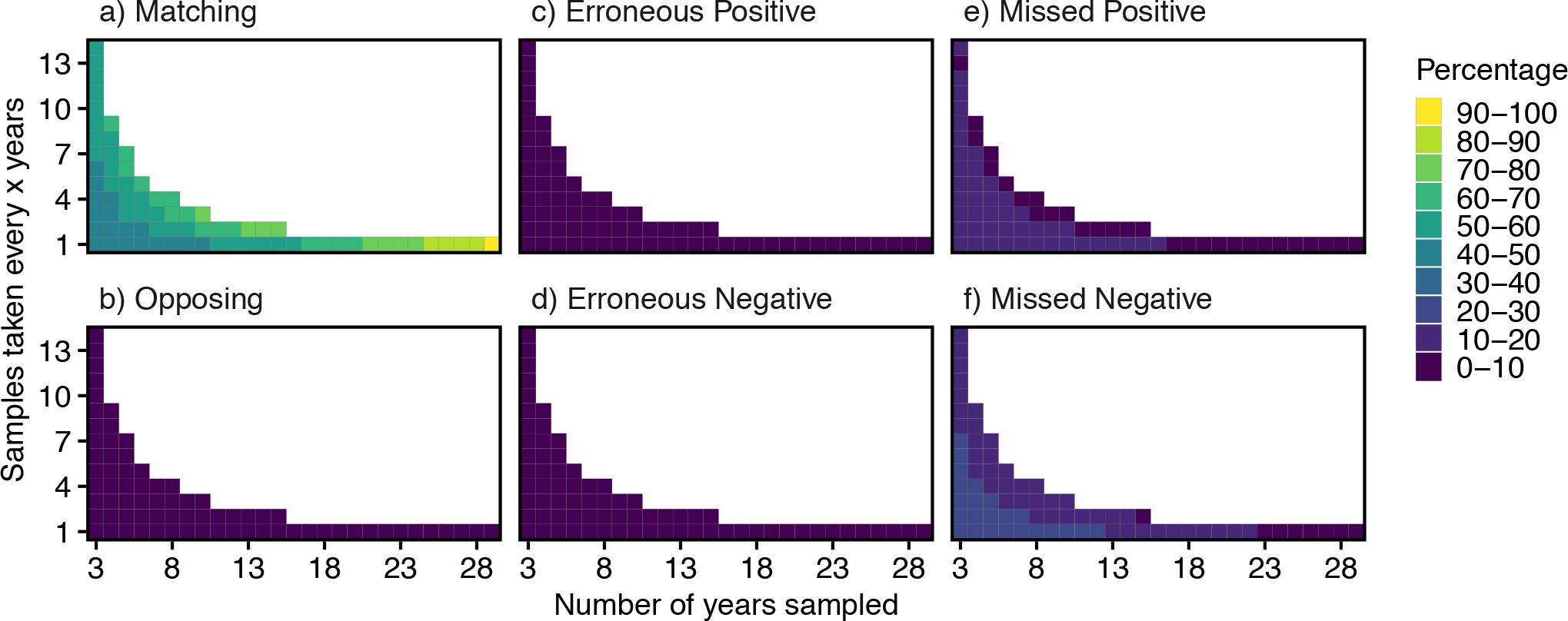
Direction Comparison of sample trends using Interval Sampling. Colour shows percentage of sample trends that, relative to the complete trend, were matching, opposing, an erroneous positive/negative or a missed positive/negative (see Table 1). Shown for all combinations of Interval Sampling, with number of years sampled (x-axis) and interval length (y-axis). Thus 8 on the x-axis and 4 on the y-axis would mean 8 samples were taken, one every 4 years. The bottom row of each plot is equal to the right most column of the equivalent Figure 3 plot, but is included here to ease comparison. Complete trend length is always 30 years.

As with Consecutive Sampling, the percentage opposing for Interval Sampling was very low (Fig. 4b), the chance of making an erroneous positive or negative was also very low for all sampling combinations (Fig. 4c, d) and, though missed positives and negatives were slightly more likely, the likelihood of missed trends never exceeded 40% (Fig. 4e, f). As before, missed negatives were more likely than positives.

### Magnitude Comparison of Consecutive Sampling

When comparing the growth rate (*r*) of sample trends to complete trends, sample trends were regularly correct only at very high tolerances. For ease of interpretation these results are displayed at four complete trend lengths: 5, 10, 20 and 30 years. In order for the sample trend *r* to be within ±0.1 (i.e. 10% population change per year) of the complete trend *r*, the sample time-series needed to be at least 9 years long when compared to complete time-series of 10 years (Fig. 5b); at least 16 years when compared to a 20-year time-series (Fig. 5c); at least 19 years when compared to a 30-year time-series (Fig. 5d); and could not be attained when the complete time-series was 5 years long (Fig. 5a). A sample of 29 years only estimated a trend within 1% (±0.01) of the 30-year trend in 80% of situations (Fig. 5d).

**Figure 5.**
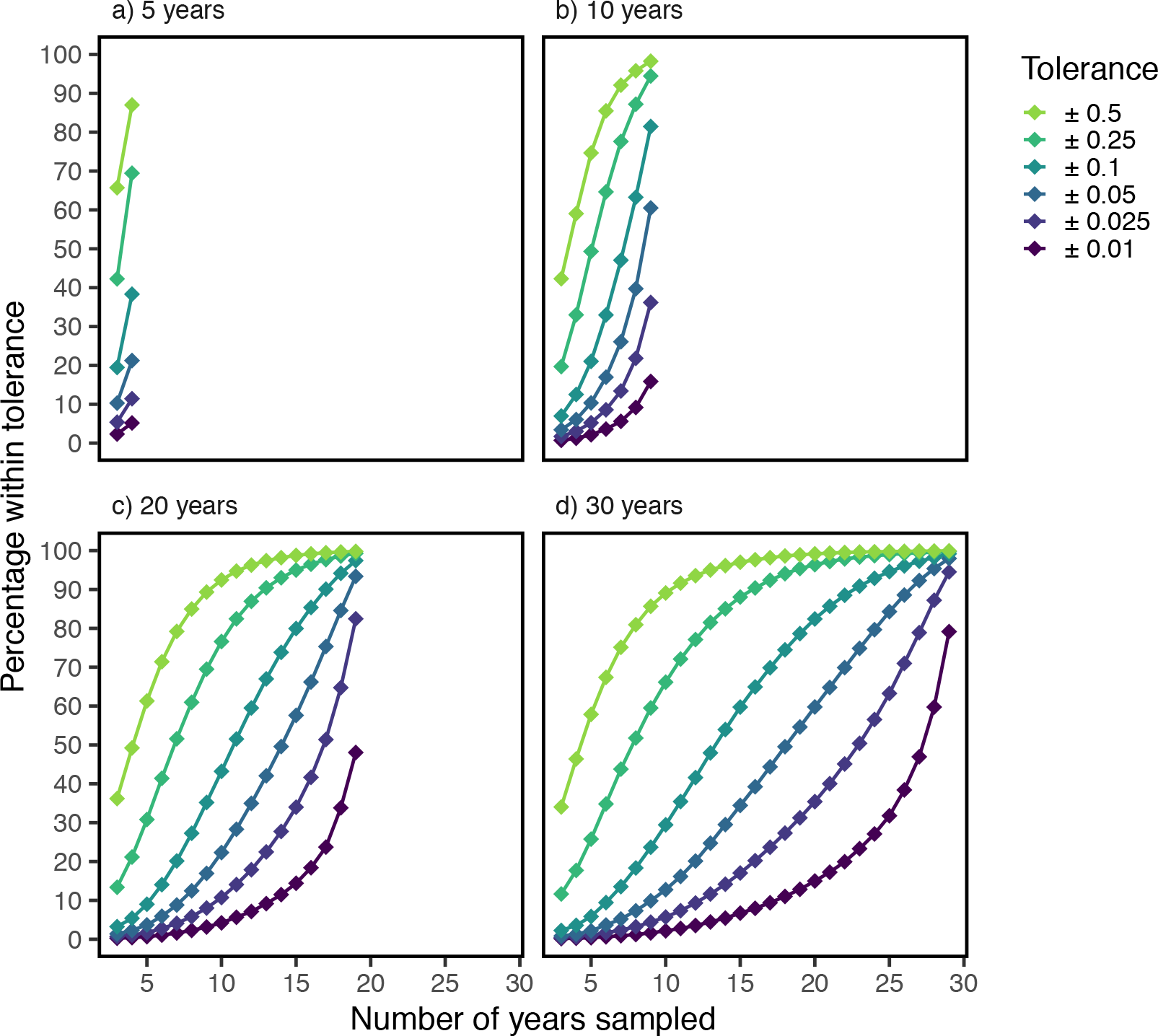
Percentage of sampled trends that correctly estimate the complete trend, measured by whether the sample r matched the complete r within the tolerance (colours). Samples taken using Consecutive Sampling. Shown for four complete trend lengths, 5, 10, 20, and 30 years, and for all sample lengths (x-axis).

### Magnitude Comparison of Interval Sampling

Interval Sampling gave better results when comparing trend direction (above), however did not perform as well for estimating magnitude of change. It was not possible for even 50% of sample trends to be correct at low thresholds (±0.01 – 0.025; Fig 6). The direction of curves indicates that high percentages could be attained with enough years of sampling at large intervals, but this would mean sampling over very long time-scales. Better reliability was achieved at higher tolerances, but only at ±0.5 and ±0.25, i.e. 25 – 50% population change per year.

**Figure 6.**
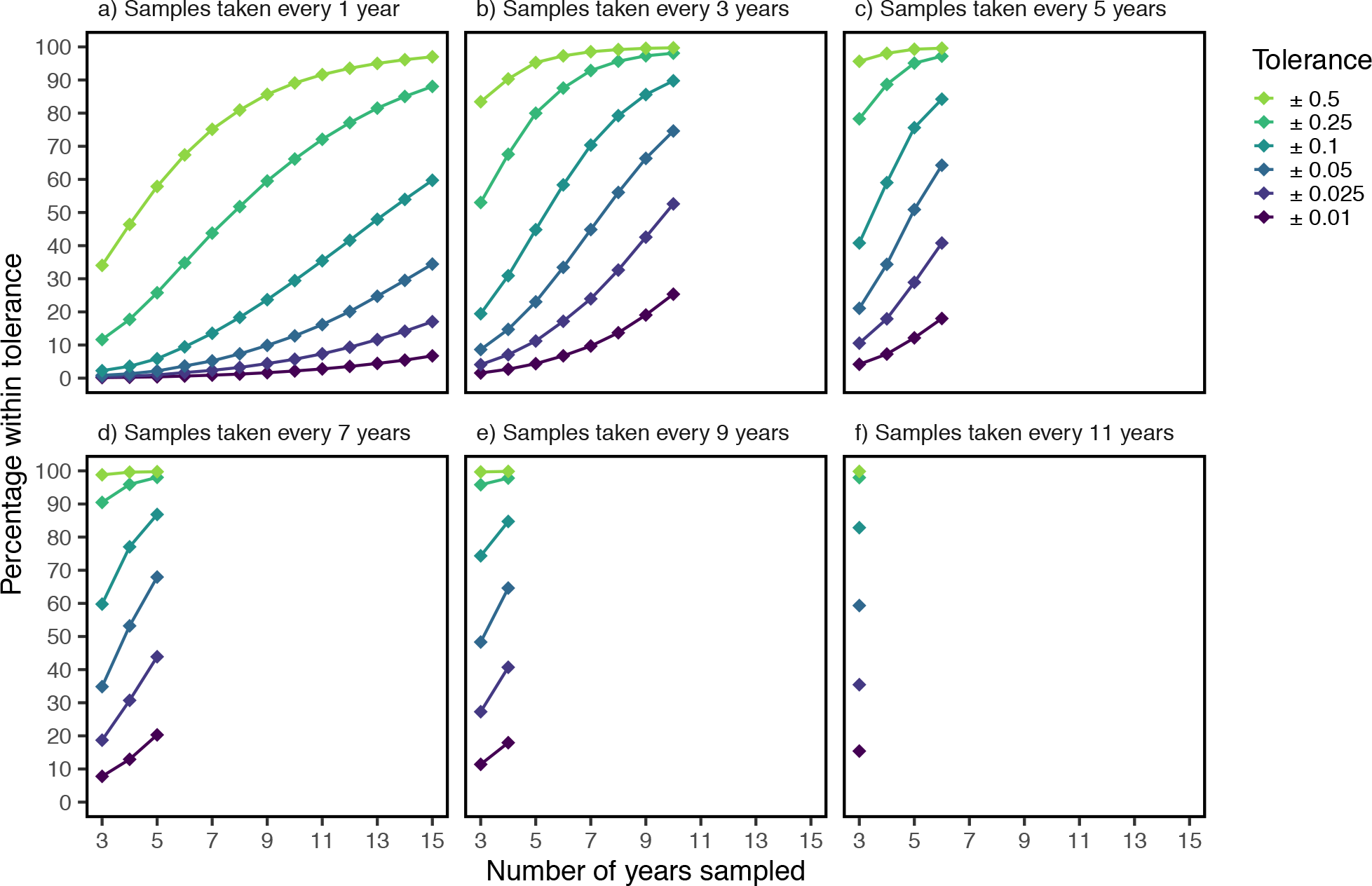
Percentage of samples (y-axis) that correctly estimate the complete trend, measured by whether the sample r matched the complete r within the tolerance (colours); samples taken using Interval Sampling. Shown for six stages of Interval Sampling: samples taken either every 1, 3, 5, 7, 9, or 11 years (facets), for all possible numbers of years sampled (x-axis). Complete trend length is always 30 years. Note that panel a is equal to Figure 5d (with a truncated x axis)

### Generation Length and Trend Shape

For populations with long generation lengths, short samples of long complete time-series were less likely to match the complete trend (Fig. 7, Supporting Figure 3). Erroneous trends and opposing trends were roughly equal among populations of different generation lengths (Fig. 7, Supporting Figure 3). Populations of different generation lengths performed similarly according to the Magnitude Comparison method (Supporting Figures 4 & 5).

**Figure 7.**
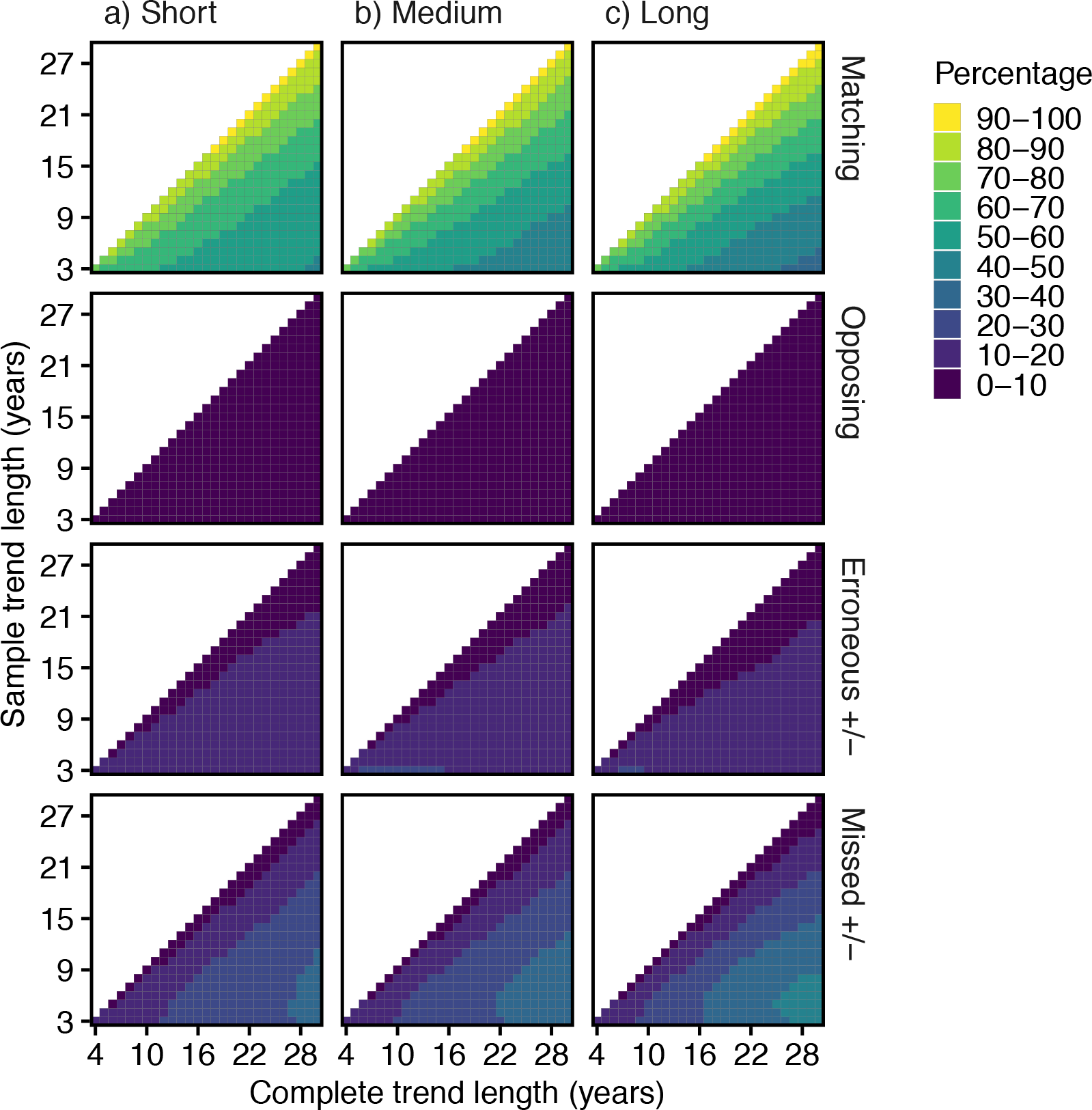
Direction Comparison of sample population trends using Consecutive Sampling, for populations of different generation lengths. Heat shows percentage of sample trends that, relative to the complete trend, were matching, opposing, an erroneous positive/negative or a missed positive/negative (see Table 1). Shown for all combinations of sample lengths (y-axis), and complete lengths (x-axis). Divided by populations with either a) short (1-5 years), b) medium (6-10 years) or c) long (11-15 years) generation lengths.

Where complete trends were highly variable across time (high estimated degrees of freedom), there were more erroneous positives and negatives with short samples of long complete time-series. However the percentage of sample trends that matched complete trends remained reasonably constant regardless of trend shape (Supporting Figures 6 & 7). Populations of different trend shapes performed similarly according to the Magnitude Comparison method (Supporting Figures 8 & 9).

## Discussion

In this paper, we provide and test a method that can estimate reliability of population trends of different lengths and sampling types, based on the total time over which a trend estimate is desired. Our results are derived from an entirely empirical dataset with no simulations and they show a high amount of convergence (e.g. Fig 5), indicating that our sample sizes are large enough to give a reliable estimate of likelihood for each category. Our results were robust to standardisation of the number of populations per species, sub-setting of species into three separate groups based on generation length (see also White, 2018) and sub-setting of populations into different trend shapes. We discuss the meaning of our results in the context of trusting population trends, how our methods can be adapted to other taxa, and how results from our methods can be used to quantify reliability in large-scale studies of population trends and to design future monitoring schemes.

### Waterbird case study

Our results show that if a *significant* trend is obtained from a population time series, even when the data are from a few years, it is likely to reflect the longer-term trend direction of the population, though not necessarily the magnitude. Further, few samples at widely spaced intervals can produce similar levels of reliability in trend direction compared to double the number of samples taken in consecutive years. However, we show that to be confident in a *lack* of trend, i.e. an insignificant result, one must obtain a very high number of samples. Further, if an insignificant trend is obtained, it is more likely to be missing a decline than an increase in a population, and we suggest caution with conclusions and decisions from insignificant trends. Keith *et al.* (2015) studying birds similarly found that past population trajectory was a good predictor of future population trajectory. Møller *et al.* (2008) and Sanderson *et al.* (2006) also found a similar, though weak, correlation between trends of migratory birds between 1970-1990 and 1990-2000. However, Keith *et al.* (2015) found that past trajectories were not a good predictor of future trajectories in mammals, salmon and other fish, meaning this method should be tested on other taxa using relevant data.

Sampling in intervals provided better, relative to effort, and surprisingly accurate results, compared to sampling in consecutive years. When sampling a fixed number of times, accuracy increased with the distance between each sample. Presumably a limit exists at some point, but according to these results it is greater than 13 years. Other studies have found similar results (Urquhart *et al.*, 1998; Starcevich *et al.*, 2018), for example Reynolds *et al.* (2011) found that surveying brown bears every 10 years gave similar model performance to surveying in 3 out of every 5 years. Interval sampling could allow, say, a greater number of sites to be surveyed over a given area (Buckland & Johnston, 2017). However it is not always practical, especially for high risk species where declines may need to be detected and acted on quickly. In addition, whilst Interval Sampling could be good for cheaply obtaining trend estimates, it is not a replacement for long term monitoring that takes yearly samples, which can provide data for analyses considering drivers of population change.

In cases where analyses or management decisions depend not only on the direction of a population trajectory, but the actual rate of change, we find that sample trends are much less likely to be representative of complete trends. To be 80% reliable at a rate of population change of 1-2.5% per year over 30 years, one must sample for at least 19 consecutive years. These results are roughly in agreement with other studies on diverse taxa that find samples should be between 10 (Rueda-Cediel *et al.*, 2015; White, 2018) and 21 years (Reynolds *et al.*, 2011). Sampling in intervals also struggled to produce accurate results: 80% reliability was never achieved if samples had to be correct at anything less than a rate of change of 10% per year (i.e. drastic population change).

This analysis was kept simple by restricting it to single location time series, and by restricting sampling to a minimum of once every year. Increasing the number of samples in each time period can improve confidence and accuracy in derived trends (Atkinson *et al.*, 2006) and it is likely that percentage reliability in derived trends would increase if this was considered. Some studies have found that sampling at more locations improved trend detection better than longer time series (Sims *et al.*, 2006, though see Schumann *et al.*, 2013) and so modelling trends from multiple locations is likely to improve our reliability estimates (Rhodes & Jonzén, 2011).

### Applications

For those working with large datasets who cannot conduct power analysis or quantify measurement error in their populations, reliability can be assigned to trends using the data from this study (see Supporting Information), or data produced using these methods with other taxa (using the provided code, see Data Accessibility). The user should define a target ‘complete’ trend length for the study, and extract the reliability estimates for this complete trend length. After this two options are possible: 1) assign reliability estimates to each time-series based on how long it is, and weight analysis according to these estimates or 2) select a threshold (e.g. time-series must be at least 80% likely to represent the complete trend) and remove any time-series that do not meet this criteria. We readily agree that our values are not infallible: they could vary with location, time period or taxa. However, this is an improvement on making an arbitrary cut-off point, or having no way of weighting population trends.

These results could also be used to help plan future monitoring schemes, but we would advise this be done with a cautious eye. The goals of monitoring have been subject to much discussion (Hauser *et al.*, 2006; Legg & Nagy, 2006; Nichols & Williams, 2006; Lindenmayer & Likens, 2009; McDonald-Madden *et al.*, 2010), but in cases where programs are carried out with the goal of detecting trends (Marsh & Trenham, 2008) we can provide some very general guidance: monitoring for many years is essential to accurately capturing population trends, and resources can be conserved and possibly allocated to more locations or taxa if sampling is conducted in intervals rather than every year.

Finally, our results could help with make cost effective management decisions, for example defining sensible allocations of effort to different species based on confidence in derived trends.

## Conclusions

In this age of increasing large-scale analyses, we believe the scientific community can do better at making informed decisions around uncertainty and reliability. Our methods and results provide a clear and quantitative way to add rigour to large scale population analyses. We advocate an end to arbitrary cut-offs, and recommend that, where possible, users instead consider methods such as ours to quantify reliability and make decisions about their data accordingly. Our methods are fully transferable to other taxa, and the concepts can also be transferred to areas outside of population ecology.

## Supporting information

SupportingInformation

SupportingData

## Acknowledgements

CBC Data is provided by National Audubon Society and through the generous efforts of Bird Studies Canada and countless volunteers across the western hemisphere. HW was supported by a Cambridge Trust Cambridge-Australia Scholarship and the Cambridge Department of Zoology JS Gardiner Fellowship. WJS is funded by Arcadia. We thank Richard Fuller, Benno Simmons and Martin Wauchope for helpful comments and discussion.

## Author Contributions

HW, TA, WJS and AJ conceived the ideas and designed methodology. TA collected the data, HW analysed the data and HW led the writing of the manuscript. All authors contributed critically to the drafts and gave final approval for publication.

## Data Accessibility

All data used in our analysis are available from the Christmas Bird Count (www.christmasbirdcount.org). Code, which is written to be fully adaptable to other data, is available both in the supporting materials and on GitHub (https://github.com/hannahchoppie/TrustingTrends), we recommend using the GitHub code as this will remain up to date.

## Supporting Information

Supporting information contains two files, a PDF containing supporting figures and explanations, and a zip file containing modelled data from our methods, and code the reproduce methods.

