## SupportingInformation for "When can we trust population trends? A method for quantifying the effects of sampling interval and duration"

### 1. Dataset summary statistics

53% of the complete datasets (i.e. the full 30 years for each of the 29,226 populations) showed significant trends at $p<0.05$. Population growth rate ($r$) showed a roughly normal distribution centered on a mean of 0.026 (Supporting Figure 1a), max growth rate was 27.8 and min growth rate was -26.8, but 99.9% of populations had growth rates between -1 and 1. Curve shape was varied, with a skew towards lower Estimated Degrees of Freedom (i.e. less complex curve shapes; Supporting Figure 1); 17% of trends were linear, i.e. EDF=1. Also shown in this section is a map of all sites (Supporting Figure 2) and a table of all species (Supporting Table 1)

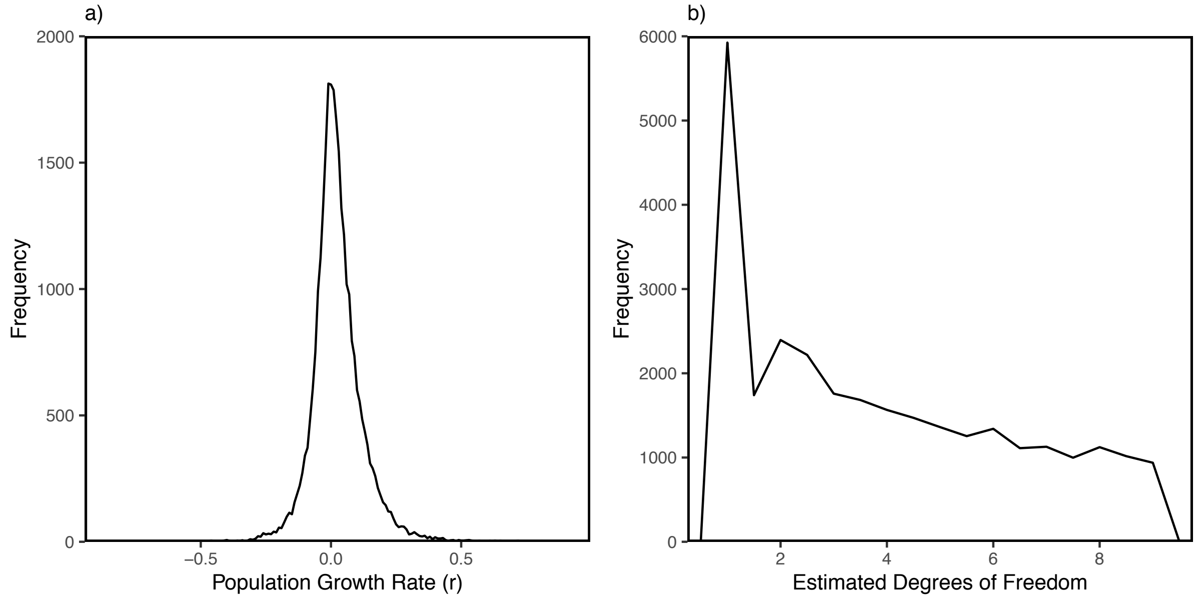

Supporting Figure 1. Summary statistics of population trends derived from 29,226 waterbird populations. Shows a) frequency distribution of population growth rate ($r$), from the 99.9% of trends that had a growth rate of greater than -1 and less than 1 and b) frequency distribution of estimated degrees of freedom.

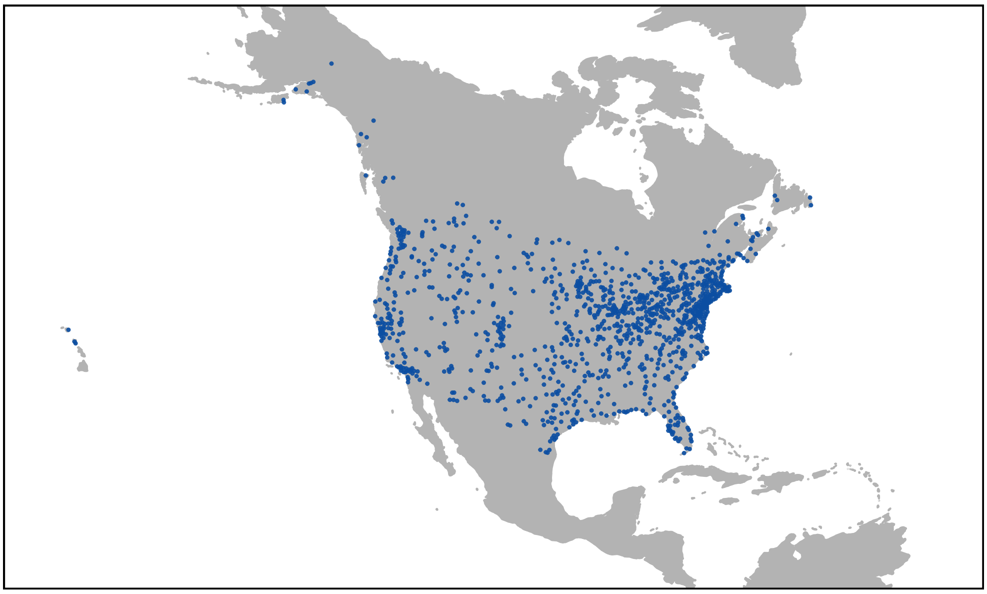

Supporting Figure 2. Map of where 1,110 sites are located across North America.

Supporting Table 1. List of species used in analysis

| **Order** | **Family** | **Species** | |
|---|---|---|---|
| Anseriformes | Anatidae | Aix sponsa | Cairina moschata |
| | | Anas acuta | Cygnus buccinator |
| | | Anas crecca | Cygnus columbianus |
| | | Anas fulvigula | Cygnus olor |
| | | Anas platyrhynchos | Dendrocygna autumnalis |
| | | Anas rubripes | Lophodytes cucullatus |
| | | Anser albifrons | Mareca americana |
| | | Anser caerulescens | Mareca penelope |
| | | Aythya affinis | Mareca strepera |
| | | Aythya americana | Melanitta americana |
| | | Aythya collaris | Melanitta fusca |
| | | Aythya marila | Mergus merganser |
| | | Aythya valisineria | Mergus serrator |
| | | Branta bernicla | Oxyura jamaicensis |
| | | Branta hutchinsii | Spatula clypeata |
| | | Bucephala albeola | Spatula cyanoptera |
| | | Bucephala clangula | Spatula discors |
| | | Bucephala islandica | |
| Charadriiformes | Alcidae | Cepphus columba | Cerorhinca monocerata |
| | Charadriidae | Charadrius alexandrinus | Charadrius vociferus |
| | | Charadrius melodus | Pluvialis squatarola |
| | | Charadrius semipalmatus | |
| | Laridae | Gelochelidon nilotica | Larus livens |
| | | Hydrocoloeus minutus | Larus marinus |
| | | Hydroprogne caspia | Larus occidentalis |
| | | Larus argentatus | Larus philadelphia |
| | | Larus atricilla | Larus thayeri |
| | | Larus californicus | Rynchops niger |
| | | Larus canus | Sterna forsteri |
| | | Larus delawarensis | Sterna hirundo |
| | | Larus fuscus | Thalasseus maximus |
| | | Larus glaucescens | Thalasseus sandvicensis |
| | | Larus heermanni | |
| | Recurvirostridae | Himantopus himantopus | Recurvirostra americana |
| | Scolopacidae | Actitis macularius | Limnodromus griseus |
| | | Arenaria melanocephala | Limnodromus scolopaceus |
| | | Calidris alba | Limosa fedoa |
| | | Calidris alpina | Numenius americanus |
| | | Calidris canutus | Numenius phaeopus |
| | | Calidris himantopus | Scolopax minor |
| | | Calidris mauri | Tringa flavipes |
| | | Calidris minutilla | Tringa melanoleuca |
| | | Calidris pusilla | Tringa semipalmata |
| | | Gallinago gallinago | Tringa solitaria |
| | Stercorariidae | Stercorarius parasiticus | Stercorarius pomarinus |
| Ciconiiformes | Ciconiidae | Mycteria americana | |
| Gaviformes | Gaviidae | Gavia immer | Gavia stellata |
| Gruiformes | Aramidae | Aramus guarauna | |

| | | | |
|---|---|---|---|
| | Rallidae | Coturnicops noveboracensis<br>Fulica americana<br>Gallinula chloropus<br>Laterallus jamaicensis<br>Porzana carolina | Rallus elegans<br>Rallus limicola<br>Rallus longirostris<br>Zapornia akool |
| Pelicaniformes | Ardeidae | Ardea alba<br>Ardea herodias<br>Botaurus lentiginosus<br>Bubulcus ibis<br>Egretta caerulea | Egretta thula<br>Egretta tricolor<br>Ixobrychus exilis<br>Nyctanassa violacea<br>Nycticorax nycticorax |
| | Pelecanidae | Pelecanus erythrorhynchos | Pelecanus occidentalis |
| | Threskiornithidae | Platalea ajaja<br>Plegadis chihi | Plegadis falcinellus |
| Podicipediformes | Podicipedidae | Aechmophorus clarkii<br>Aechmophorus occidentalis<br>Podiceps auritus<br>Podiceps grisegena | Podiceps nigricollis<br>Podilymbus podiceps<br>Tachybaptus dominicus |
| Suliformes | Anhingidae | Anhinga anhinga | |
| | Phalacrocoracidae | Phalacrocorax auritus<br>Phalacrocorax brasilianus | Phalacrocorax pelagicus<br>Phalacrocorax penicillatus |
| | Sulidae | Morus bassanus | |

### 2. Generation length

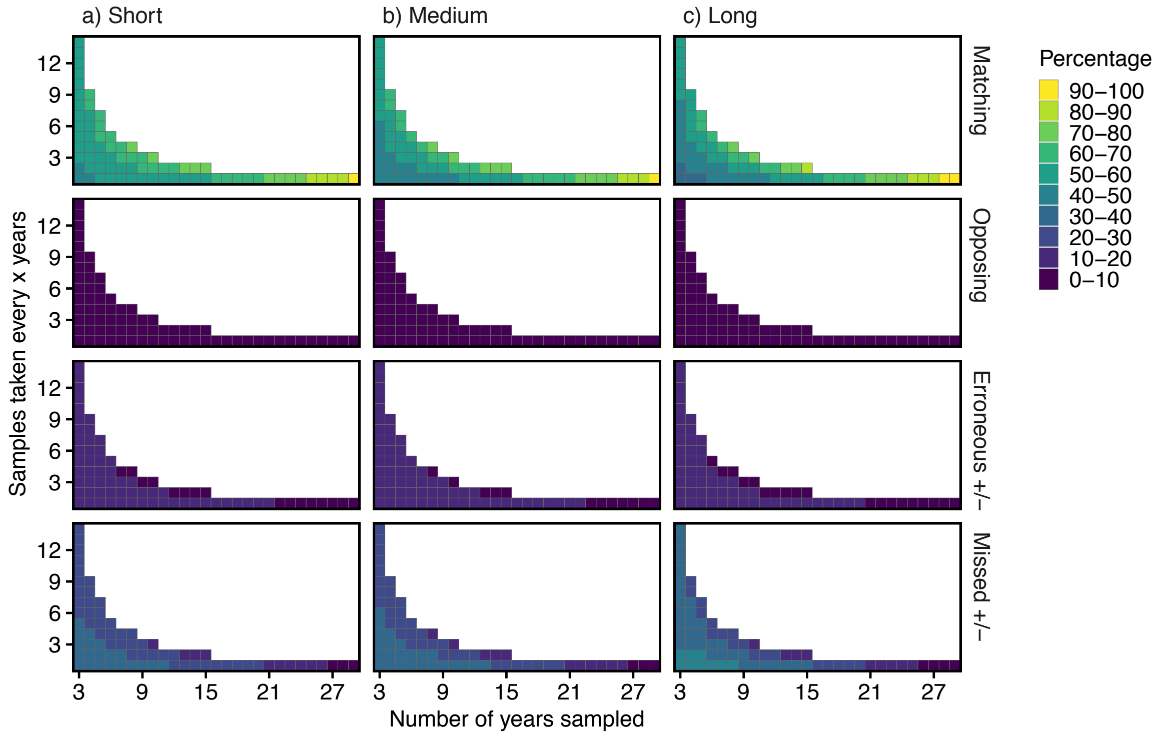

Supporting Figure 3. Direction comparison of sample trends using interval sampling. Heat shows percentage of sample trends that, relative to the complete trend, were correct, opposing, a false positive/negative or a missed positive/negative (see Table 1). Shown for all combinations of interval sampling, with number of years sampled (x-axis) and interval length (y-axis). Thus 8 on the x-axis and 4 on the y-axis would mean 8 samples were taken, one every 4 years. The bottom row of each plot is equal to the right most column of the equivalent Figure 7 plot, but is included here to ease comparison. Complete trend length is always 30 years. Divided by populations with either a) short (1-5 years), b) medium (6-10 years) or c) long (11-15 years) generation lengths.

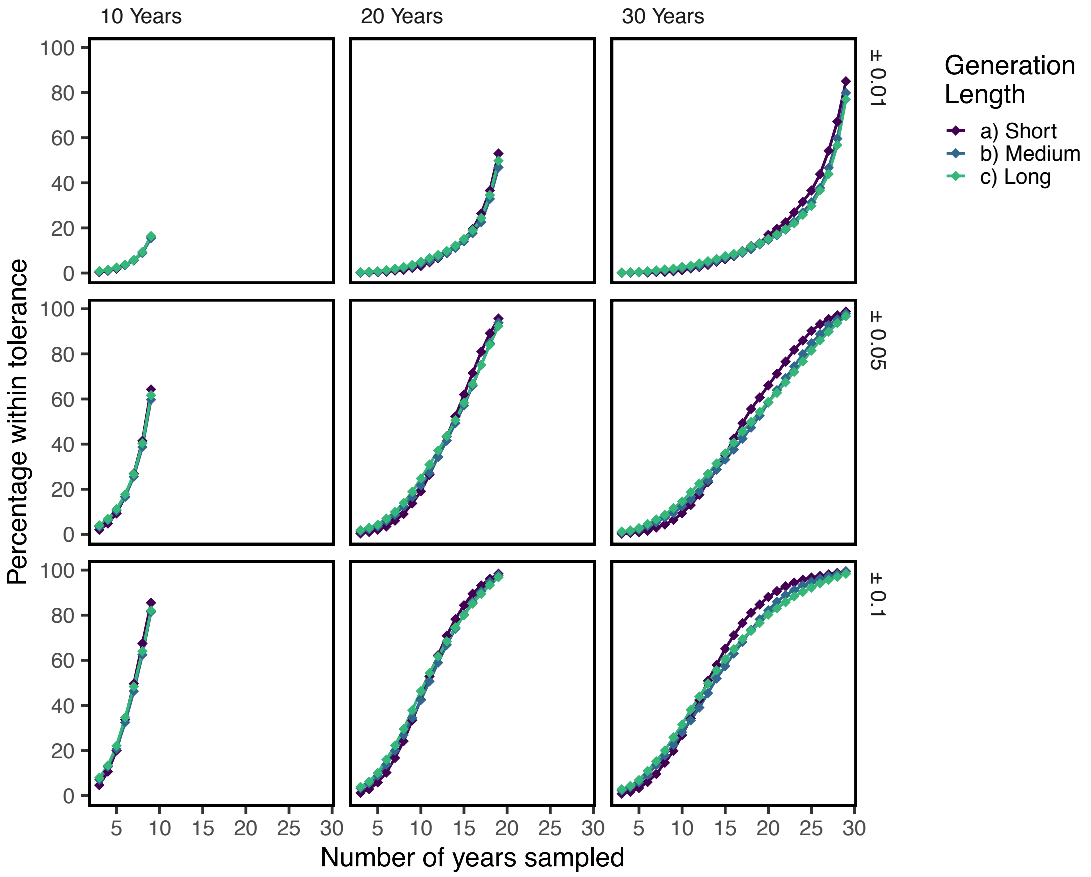

Supporting Figure 4. Percentage of sampled trends that correctly match the complete trend (x axis), measured by whether the sample *r* matches the complete *r* within the threshold (rows); samples taken using Consecutive sampling. Shown for three complete trend lengths, 10, 20, and 30 years (columns), and for all sample lengths (y-axis). Divided by populations with either a) short (1-5 years), b) medium (6-10 years) or c) long (11-15 years) generation lengths (colours).

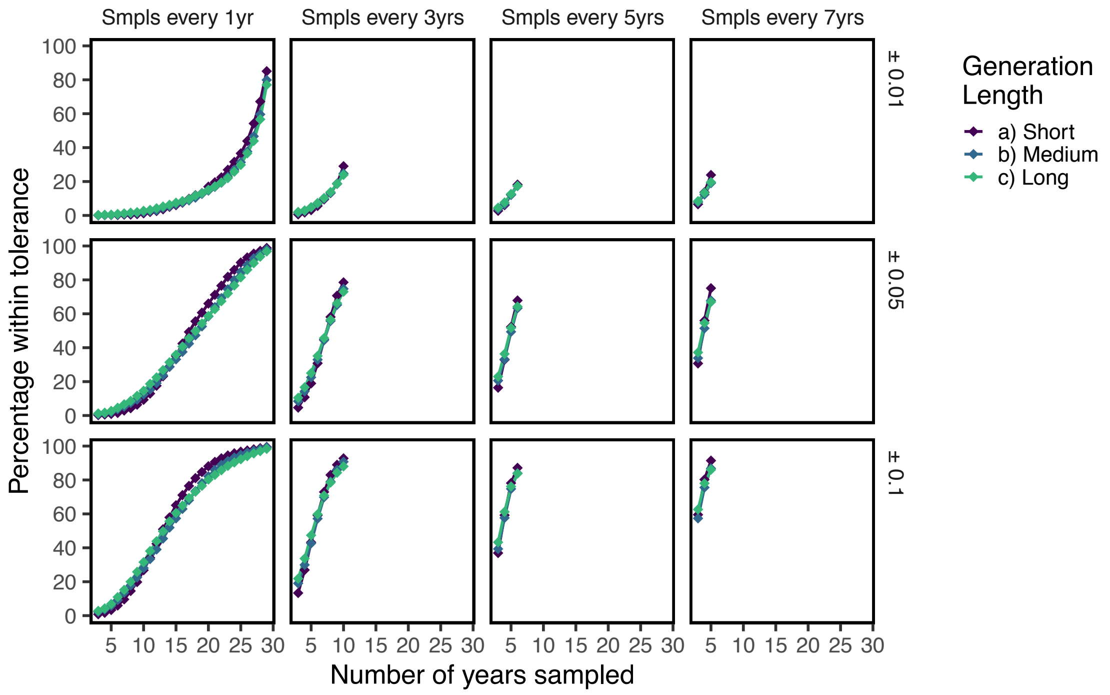

Supporting Figure 5. Percentage of samples (y-axis) that correctly match the complete trend, measured by whether the sample *r* matches the complete *r* within the threshold (rows); samples taken using Interval sampling. Shown for four stages of interval sampling: samples taken either every 1, 3, 5 or 7 years (columns), for all possible number of years sampled (x-axis). Complete trend length is always 30 years. Divided by populations with either a) short (1-5 years), b) medium (6-10 years) or c) long (11-15 years) generation lengths (colours).

### 3. Trend shape

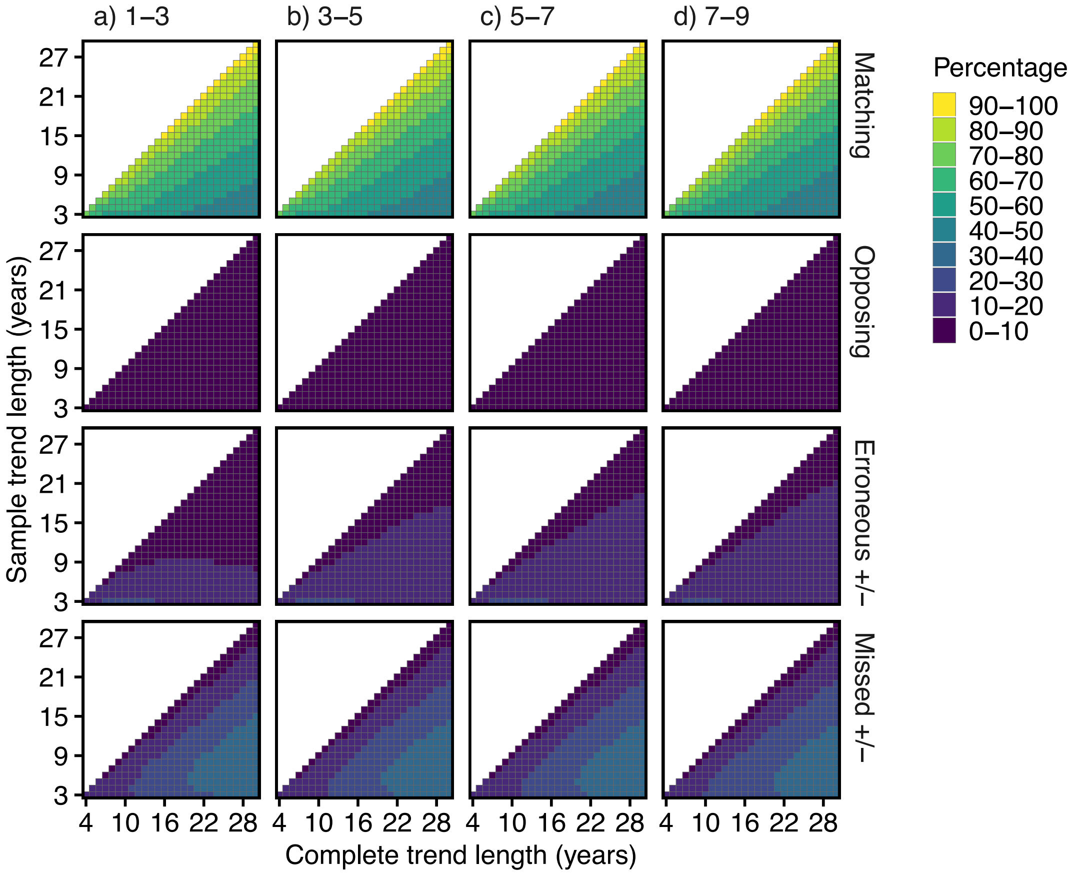

Supporting Figure 6. Direction comparison of sample population trends using consecutive sampling. Heat shows percentage of sample trends that, relative to the complete trend, were correct, opposing, a false positive/negative or a missed positive/negative (see Table 1). Shown for all combinations of sample lengths (y-axis), and complete lengths (x-axis). Divided by complete trends with estimated degrees of freedom (EDF) of 1-3, 3-5, 5-7 and 7-9.

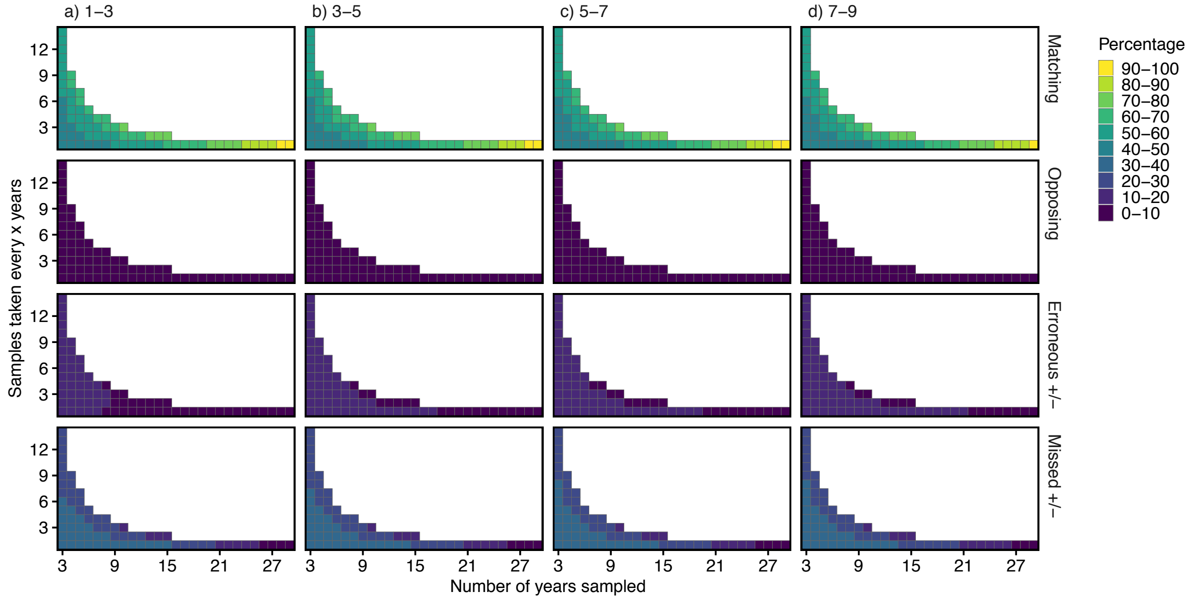

Supporting Figure 7. Direction comparison of sample trends using interval sampling. Heat shows percentage of sample trends that, relative to the complete trend, were correct, opposing, a false positive/negative or a missed positive/negative (see Table 1). Shown for all combinations of interval sampling, with number of years sampled (x-axis) and interval length (y-axis). Thus 8 on the x-axis and 4 on the y-axis would mean 8 samples were taken, one every 4 years. The bottom row of each plot is equal to the right most column of the equivalent Supporting Figure 5 plot, but is included here to ease comparison. Complete trend length is always 30 years. Divided by complete trends with estimated degrees of freedom (EDF) of 1-3, 3-5, 5-7 and 7-9.

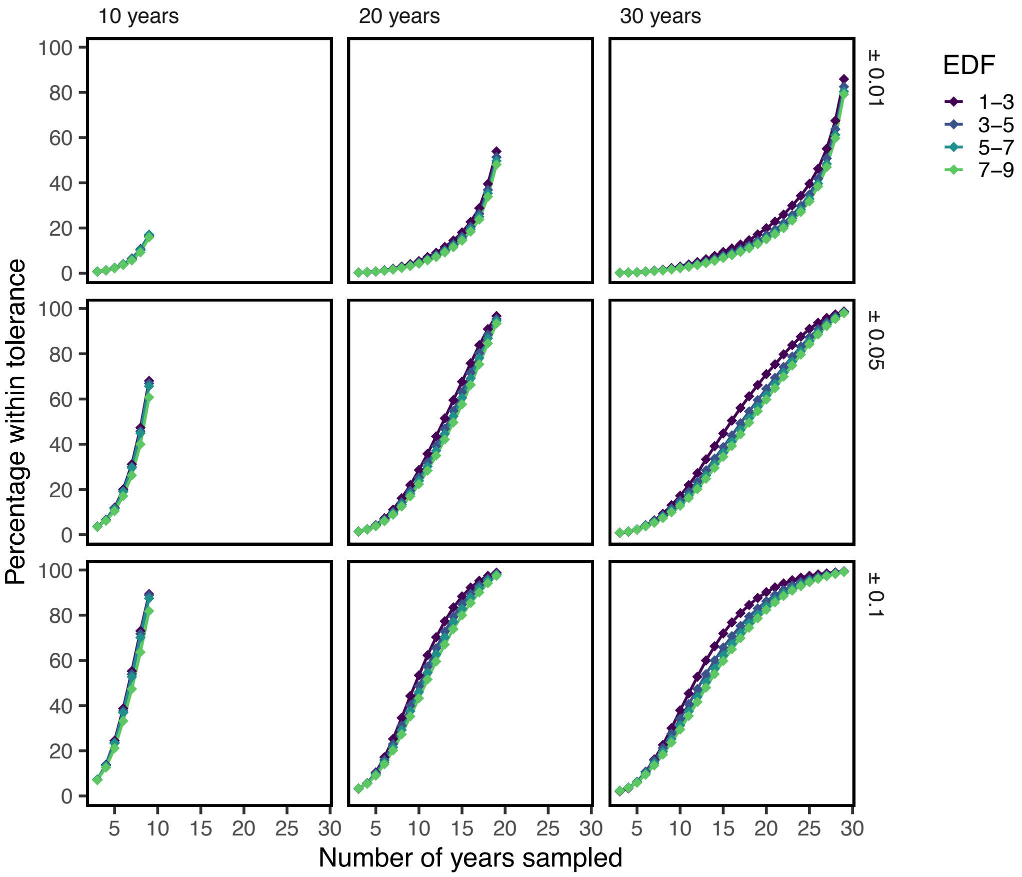

Supporting Figure 8. Percentage of sampled trends that correctly match the complete trend (x axis), measured by whether the sample *r* matches the complete *r* within the threshold (rows); samples taken using Consecutive sampling. Shown for three complete trend lengths, 10, 20, and 30 years (columns), and for all sample lengths (y-axis). Divided by complete trends with estimated degrees of freedom (EDF) of 1-3, 3-5, 5-7 and 7-9 (colours).

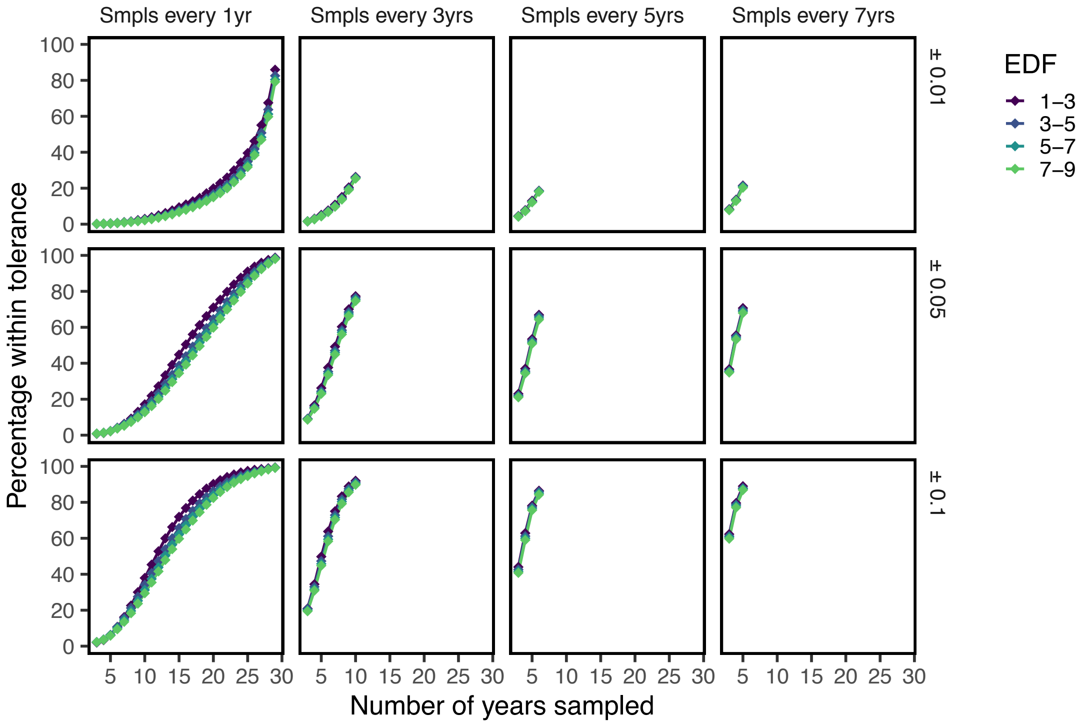

Supporting Figure 9. Percentage of samples (y-axis) that correctly match the complete trend, measured by whether the sample *r* matches the complete *r* within the threshold (rows); samples taken using Interval sampling. Shown for four stages of interval sampling: samples taken either every 1, 3, 5 or 7 years (columns), for all possible number of years sampled (x-axis). Complete trend length is always 30 years. Divided by complete trends with estimated degrees of freedom (EDF) of 1-3, 3-5, 5-7 and 7-9 (colours).

### 4. Supplementary investigation: Rate of matching slopes for insignificant sample trends compared to significant complete trends.

A final investigation we felt would be important to conduct was the frequency with which the slope of *insignificant* sample trends matched the complete slope of *significant* trends. In the paper, we recorded these cases as missed positives/negatives, and did not consider the slope of the insignificant samples. However, in some applications it may be useful to know how likely it is that the direction of the insignificant slope matches the direction of the complete trend.

For this reason we conducted a final investigation, looking only at cases where the complete time-series trend was significant, and the sample trend was insignificant (i.e. the missed positives/negatives of the Direction Comparison method). We calculated how often the direction of the slope of the insignificant sample trend matched the direction of the slope of the complete trend. For Consecutive Sampling, we found that rates of agreement were high, with sample trends being 90-100% likely to match the complete trends when the sample data was approximately 3/4s of the length of the complete data (Supporting Figure 10). When the sample data was a small proportion of the complete data, chances of it matching were at lowest 50-60%. Similar rates of agreement occurred when sampling in intervals, not consecutive years (Supporting Figure 11).

We also investigated how the magnitude of slope compared between insignificant sample trends and significant complete trends. Results were similar to comparing significant sample and complete trends (as in the main paper), except that in this case larger samples were needed for trend magnitudes to be within tolerance thresholds (Supporting Figures 12 & 13).

Code and data for these analysis are provided in the supporting code and results zip file, see below.

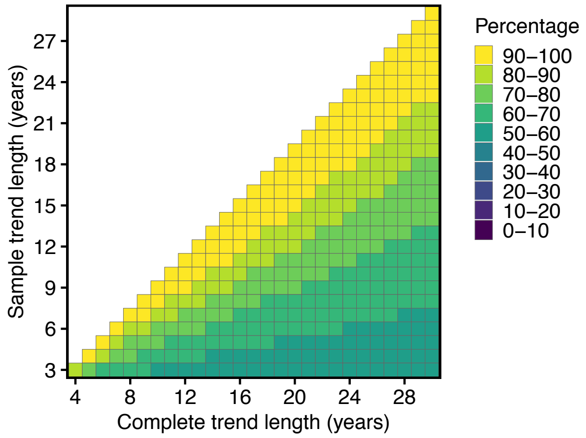

Supporting Figure 10. Direction comparison of insignificant sample population trends to significant complete population trends using Consecutive Sampling. Colour shows percentage of sample trends that, relative to the complete trend, were matching. Shown for all combinations of sample lengths (y-axis), and complete lengths (x-axis).

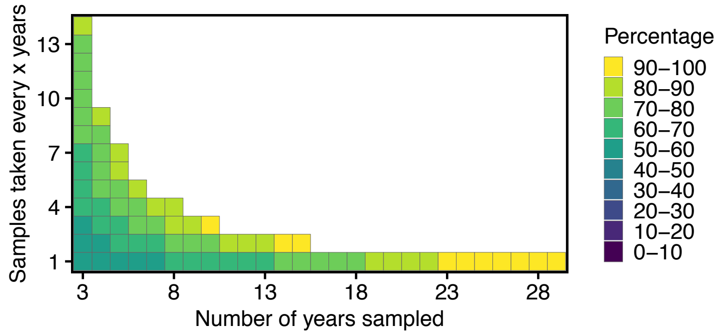

Supporting Figure 11. Direction comparison of insignificant sample trends to significant complete trends using Interval Sampling. Colour shows percentage of sample trends that, relative to the complete trend, were matching. Shown for all combinations of Interval Sampling, with number of years sampled (x-axis) and interval length (y-axis). Thus 8 on the x-axis and 4 on the y-axis would mean 8 samples were taken, one every 4 years. Complete trend length is always 30 years.

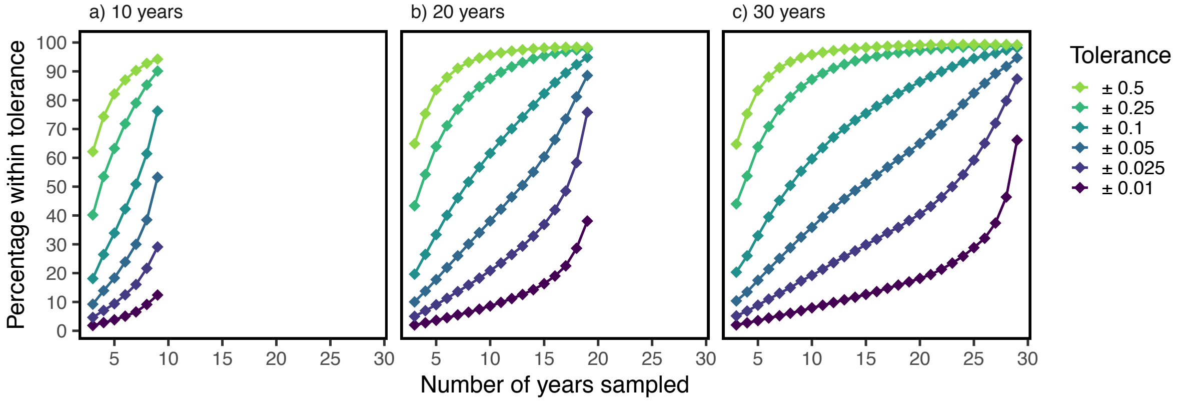

Supporting Figure 12. Percentage of insignificant sampled trends that correctly estimate the significant complete trend, measured by whether the sample $r$ matches the complete $r$ within the tolerance (colours). Samples taken using Consecutive Sampling. Shown for three complete trend lengths, 10, 20, and 30 years, and for all sample lengths (x-axis).

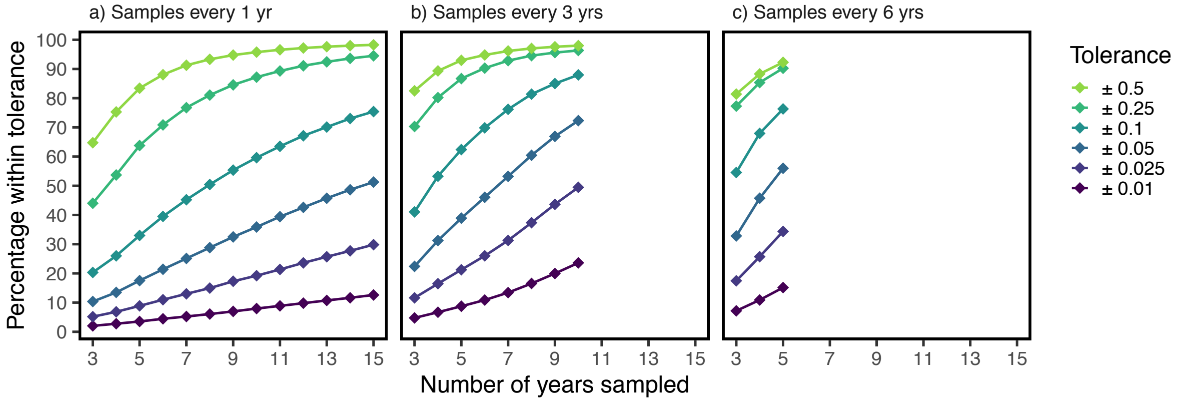

Supporting Figure 13. Percentage of insignificant samples (y-axis) that correctly estimate the significant complete trend, measured by whether the sample $r$ matches the complete $r$ within the tolerance (colours); samples taken using Interval sampling. Shown for three stages of Interval Sampling: samples taken either every 1, 3 or 6 years (facets), for all possible number of years sampled (x-axis). Complete trend length is always 30 years.

### 5. Description of Supporting code and results zip file

The Supporting code is fully annotated code detailing all analysis, with instructions on transferring to other data. The results file contains the full data, detailing propotions of confidence for samples of various lengths (and intervals) when compared to complete trends also of varying lengths. The columns in each file are as follows:

- DirectionDataConsecutive.csv:
  - SampleLength: The length of the sample
  - Prop: The proportion of samples that were the category (e.g. if Category is Correct and Prop is 0.71, then 71% of samples were correct for that sample and complete length)
  - Category: See main paper Table 1 for direction categories, either Correct, Erroneous Positive, Erroneous Negative, Missed Positive, Missed Negative or Opposing.
  - CompleteLength: The length of the complete time-series that the sample was compared to
  - EDF: The estimated degrees of freedom of complete trends in that group of data (if NA, full dataset used, otherwise if 3: EDF was 1-3, 5 = 3-5, 7 = 5-7 and 9 = 7-9)
  - GenLength: The generation length of species in that group of data (if NA, full dataset used, otherwise if 5: generation length was 1-5, 10 = 6-10 and 15 = 11-15)
- DirectionDataInterval.csv
  - IntervalLength: Samples taken every x years (e.g. if 1, samples taken every year, if 2 every two years etc)
  - Category: See main paper Table 1 for direction categories, either Correct, Erroneous Positive, Erroneous Negative, Missed Positive, Missed Negative or Opposing.
  - NumYearsInSample: The actual number of samples (e.g. if IntervalLength is 3 and NumYearsInSample is 4, then 4 samples were taken, one every 3 years)
  - Prop: The proportion of samples that were the category (e.g. if Category is Correct and Prop is 0.71, the 71% of samples were correct for that sample and complete length)
  - CompleteLength: The length of the complete trend that the sample was compared to (nb. always 30 for interval data)
  - EDF: The estimated degrees of freedom of complete trends in that group of data (if NA, full dataset used, otherwise if 3: EDF was 1-3, 5 = 3-5, 7 = 5-7 and 9 = 7-9)
  - GenLength: The generation length of species in that group of data (if NA, full dataset used, otherwise if 5: generation length was 1-5, 10 = 6-10 and 15 = 11-15)
- MagnitudeDataConsecutive.csv:
  - SampleLength: The length of the sample
  - PropWithinTolerance: The proportion of samples for which absolute difference between the sample growth rate (r) and compelte growth rate was not more than the tolerance.
  - Tolerance: Amount by which sample and complete growth rate can differ
  - CompleteLength: The length of the complete time-series that the sample was compared to (nb. always 30 for interval data).

  - EDF: The estimated degrees of freedom of complete trends in that group of data (if NA, full dataset used, otherwise if 3: EDF was 1-3, 5 = 3-5, 7 = 5-7 and 9 = 7-9)
  - GenLength: The generation length of species in that group of data (if NA, full dataset used, otherwise if 5: generation length was 1-5, 10 = 6-10 and 15 = 11-15)
- MagnitudeDataInterval.csv:
  - IntervalLength: Samples taken every x years (e.g. if 1, samples taken every year, if 2 every two years etc)
  - NumYearsInSample: The actual number of samples (e.g. if IntervalLength is 3 and NumYearsInSample is 4, then 4 samples were taken, one every 3 years)
  - PropWithinTolerance: The proportion of samples for which absolute difference between the sample growth rate (r) and compelte growth rate was not more than the tolerance.
  - Tolerance: Amount by which sample and complete growth rate can differ
  - CompleteLength: The length of the complete trend that the sample was compared to
  - EDF: The estimated degrees of freedom of complete trends in that group of data (if NA, full dataset used, otherwise if 3: EDF was 1-3, 5 = 3-5, 7 = 5-7 and 9 = 7-9)
  - GenLength: The generation length of species in that group of data (if NA, full dataset used, otherwise if 5: generation length was 1-5, 10 = 6-10 and 15 = 11-15)
  -

The same column names apply to the files ending in “SupportingDataSection4.csv”, just without the category section (see Section 4 above for an explanation of how this data was created).
