## SupportingData for "When can we trust population trends? A method for quantifying the effects of sampling interval and duration": WauchopeQuantifyingTrends.R

```
### WELCOME ###
### This code is to calcuate confidence in population trends by artifically degrading a count dataset of long term time-series, as used in the paper "When can we trust population trends? A method for quantifying the effects of sampling interval and duration" by Hannah Wauchope, Alison Johnston, Tatsuya Amano and William Sutherland (2019).
### Currently available on Biorxve https://www.biorxiv.org/content/early/2018/12/20/498170
### Produced by Hannah Wauchope, please contact at with any questions/comments/errors you've found. 
### Refer to the paper for explanations of methods, terms used etc.

### THIS CODE CONTAINS THE FOLLOWING SECTIONS ###
#Load packages and read data. This loads all require packages, and sets up filepaths for importing and exporting data, this should be adjusted if code is being used for your own data
#Prepare Count Data and Prepare Generation Length Data. This shows how I organised the data for the paper. These are relevant for exact replicas of my results using CBC downloaded data, but otherwise are only useful as reference.
#Functions. Shows the functions used to compare trends between subsets and complete datasets (by Direction and Magnitude)
#Models 1. Consecutive. Divided into two sections, Run Models and Summarise. In 'Run Models' GLMs are run on all iterations of subsets of the data (i.e. all possible numbers of years, and all possible start years)
  #This data is saved, with columns indicating the SiteSpec (i.e. population), the slope from the model (i.e. population growth rate), the significance of the slope, the estimated degrees of freedom from the GAM run on the data (for the trend shape section of analysis), 
    #the number of years of the sample, the start year of the sample, the sampling type (i.e. consecutive) and model family type (in my case negative binomial)
  #Next, there is the summarise section, which takes all the results from the model runs and compares samples to complete trends according to both direction and magnitude methods, as well as comparing different generation lengths, trend shapes etc. 
#Models 2. Intervals. This is organised the same as Models 1, but for interval sampling instead (see paper). 
#Plot. This section plots all results. This has not been fully optimised to adapt to any results (as it very much depends on what dataset is being used) so will need some tweaking if you wish to apply it to your data, e.g. for axis names etc
#Paper stats. This is a small section just showing how I calculated any statistics reported in the paper/supporting material. 
#Build Supporting data. This organises and saves all model outputs in the format given in supporting materials. 
#Supplementary analysis for supporting information section 4

### TO APPLY ANALYSES TO YOUR OWN DATA ###
#Set all your file paths in the "Load packages and read data" section, make sure you have all packages installed
#Skip to "Models" Sections. 
#Obtain a number of populations with the same number of years surveyed (in my case, it was populations with 30 years surveyed). Set the MCL ("Max Complete Length") within "Load packages and read data" to be whatever that number of years is.
#Set the model family function according to which is best for your data. The options provided are Poisson ("pois"), Quasi-poisson ("quasi") or negative binomial ("nb") (though obviously with more intensive code tweaking you could adjust the actual GLMs to whatever you like). Quasi-poisson and NB are better for over dispersed data
  #Input your choice under 'modelfam' in "Load packages and read data" 

#All you need for data is a list named "TYB_list", saved as an RData file within ResultsFP. This should be a list of dataframes, each being the counts of one population at one site (with MCL number of years). There should be 4 columns, named as follows:
  ##SiteSpec - The name of the Site and Species (or your population name, doesn't really matter BUT must be unique for each dataframe)
  ##Year - The year of data, should be sorted smallest to largest
  ##Count - The number of counts in that year
  ##Hours - An effort term, in my case the number of hours of effort that went into collecting the count. This can be any kind of effort value, as long as it's consistent between all populations. Note that across all populations log(effort) should correlate with count (see main manuscript, and second section of "Prepare Count Data" in this file)
    #If effort is consistent (i.e. counts were taken in a standardised way) this column should just be 1 for all values (NOT 'NA'. I know this is slightly unconventional, but it's the most efficient way to exclude this term without lots of extra code)
#If you like, you can include a list of population names that represents a standardised subset (see final section of Prepare Count Data). This should be named "TYB_St.RData", saved within ResultsFP, and just be a vector of SiteSpec names that match with the names in the main dataset.
  #If you do not want to include any standardised records simply do not run "Direction_St" or "Magnitude_St" at the end of each summarise section in Models 1 and Models 2
#If you like, you can also include data on the generation length of each species. You require a dataframe, saved as "GenLengthSpec.csv" within ResultsFP, with two columns:
  ##SiteSpec - the name of each population (that should correspond to the SiteSpec in TYB_list)
  ##GenLengthMax - the 'bin' which each generation length falls into. In my case I specified 3 bins, 5 = Gen Lengths of 1-5, 10 = 6-10 and 15 = 11-15. 
    ## You can set this to whatever you like, but make sure that, for whatever bins you decide on, you adjust the lapply for Direction_All, Direction_St, Magnitude_All and Magnitude_St at the end of each Summarise section to be: 0 (which indicates include data of all genlengths) and the bin values you have chosen (e.g. 0,5,10,15)
  #If you do not want to consider generation length, in the lapply for Direction_All, Direction_St, Magnitude_All and Magnitude_St at the end of each Summarise section, remove the 5,10 and 15 and just run on 0.
#If you do not want to consider curve shape, simply do not run the Direction_EDF or Magnitude_EDF
#If you are interested in different p value cut offs for significance, adjust at the end of the summarise section also.

#### Load packages and read data ####
library(plyr)
library(dplyr)
library(ggplot2)
library(data.table)
library(reshape2)
library(tidyr)
library(foreign)
library(pbapply)
library(parallel)
library(pbmcapply)
library(gridExtra)
library(scales)
library(stringr)
library(mgcv)
library(e1071)
library(MASS)
library(viridis)
library(gridExtra)

DataFP <- "<insert your data filepath>" #This folder should contain the count data for your populations
ResultsFP <- "<insert filepath to where you will store results>" #This file path is where all data is output (And read in for summaries) it should first contain the formatted data you are using, at minimum TYB_list.RData, but possibly also "TYB_St.Rdata" and "GenLengthSpec.csv". Next, it should also contain a folder named "GLM", and within this folder should be three folders, "Consecutive", "Intervals" and "Summaries" (+ "SummariesInsig if doing the final extra bit of supplementary analysis)
FiguresFP <- "<insert filepath to where you will export figures>" #This is where figures are saved. It should contain four folders, "Direction", "Magnitude", "Gen" and "EDF" (+ another folder named "Insig" containing "Direction" and "Magnitude" folders, if doing the extra supplementary analysis)
ncores <- 4 #for parallelising code. nb - the scale of data I ran these analyses at, i.e. ~29,000 populations with 30 years of counts each, took approx 960 hours to run on one core (i.e. 30 hours on 32 cores). Increasing the number of years or number of populations would increase these, so be aware!

load(file=paste0(DataFP, "birdcounts.RData")) #My original unsorted data (Containing site name, species name, year, count, and 'hours'(for effort), used in "prepare count data")

load(file=paste0(ResultsFP, "TYB_list.RData")) #Created in "Prepare count data", OR supply your own (See above)
load(file=paste0(ResultsFP, "TYB_St.RData")) #Created in "Prepare count data", OR supply your own (See above)
GenLengthSpec <- read.csv(paste0(ResultsFP, "GenLengthSpec.csv")) #Created in "Prepare generation length data", OR supply your own (See above)
MCL <- 30 #MCL = Max Complete Length
modelfam <- "nb" #this can be adjusted to "pois" or "quasi" if you want to test other model families

#### Prepare count data ####

##Reduce data set to sites with at least MCL years of surveying
SitesAndYears <- unique(birdcounts[c("SiteCode", "Species", "Year")]) #Get the years recorded at each survey site
SiteByYear <- dcast(SitesAndYears, SiteCode + Species ~., length, value.var = "Year") #Cast for number of years at each site
names(SiteByYear) <- c("SiteCode", "Species", "NumYears") #Add names
ThirYears <- subset(SiteByYear, NumYears>29) #Cut to sites that have at least MCL years of sampling

ThirYearBirds <- merge(birdcounts, ThirYears, by=c("SiteCode", "Species")) #Bring back in the bird counts
TYB <- ThirYearBirds[c("SiteCode", "Species", "PopulationCode", "Year", "Count", "Hours", "NumYears")] #Simplify dataset

##Because data comes from CBC, need to standardise for survey hours. 
##First we need to see which species show a linear trend between log(survey effort) and counts, and then reduce to only those
##We then include hours as an offset term in the models
TYB$SpecPop <- paste(TYB$Species, TYB$Population, sep="_") #Combine species name and population into one variable
birdcounts$SpecPop <- paste(birdcounts$Species, birdcounts$Population, sep="_") #Now do the same for bird counts, cos we want as many counts as possible for our hours check (not just those with MCL years)

CountvsHoursLogHours <- rbindlist(pbmclapply(unique(TYB$SpecPop), function(i){ #Cycle through each SpecPop
  print(i)
  birdie <- subset(birdcounts, SpecPop==i) #Subset dataframe to specpop
  glmmy <- tryCatch(glm.nb(Count~log(Hours), link=log, data=birdie), error=function(e){
    print(e)
    return(NULL)
  })
  if(is.null(glmmy)){return(NULL)}
  significant <- as.data.frame(ifelse(summary(glmmy)$coeff[-1,4]< 0.05, coef(glmmy, silent=TRUE)[[2]], NA)) #Return the coefficient IF the p value is less than 0.5, otherwise NA
  significant$SpecPop <- i #Add names
  names(significant) <- c("Slope", "SpecPop") #Add names
  return(significant)
}, mc.cores=ncores))

CountvsHoursLogHours <- CountvsHoursLogHours[!is.na(CountvsHoursLogHours$Slope),] #Remove the NAs (i.e. insignificant relations)
CountvsHoursLogHours <- subset(CountvsHoursLogHours, Slope>0) #Remove the negative slopes
TYB <- TYB[TYB$SpecPop %in% CountvsHoursLogHours$SpecPop,] #Reduce main dataset to just cleaned species

##Ok and now continue to clean up the data

TYB$SiteSpec <- paste(TYB$Site, TYB$SpecPop, sep="_") #Combine site and specpop into one field (So we now have a unique identifier for each 'population' i.e. each species at each site)
TYB <- TYB[,c("SiteSpec", "Year", "Count", "Hours")] #Reduce dataset to just what we need
TYB_list <- split(TYB, f = TYB$SiteSpec) #Divide into list based on sitespec

TYB_list <- lapply(TYB_list, function(x){ 
  x <- x[order(x$Year),]
  x$Count <- round(x$Count, 0)
  return(x[(nrow(x)-(29)):nrow(x),])
})#This loop reduces each list element to just MCL years of counts, using the most recent. And also rounds counts because there's a few decimal ones

TYB_list <- lapply(TYB_list, function(x) ifelse(sum(x$Count)<MCL, return(NULL), return(x))) #Remove any list elements with less than MCL counts over the MCL years
TYB_list <- TYB_list[!sapply(TYB_list, is.null)] #Remove null elements

save(TYB_list, file=paste0(ResultsFP, "TYB_list.RData")) #Save

##And now create a standardised list of species
TYB_Names <- as.data.frame(names(TYB_list))
names(TYB_Names) <- "SiteSpec"
TYB_Names$Spec <- as.data.frame(str_split_fixed(TYB_Names$SiteSpec, "[_]", 3))[,2] #Get species names
TYB_Names_Cast <- dcast(TYB_Names, Spec~., length, value.var="SiteSpec") #Find out how many sites of each species
TYB_Names_Cast_50 <- subset(TYB_Names_Cast, .>49) #Subset to species with at least 50 sites

#For each of those species, randomly extract 50 sites
TYB_Standardising <- TYB_Names[TYB_Names$Spec %in% TYB_Names_Cast_50$Spec,] #Get the sitespec values for each of the species
TYB_Standardising$Spec <- factor(TYB_Standardising$Spec, levels=unique(TYB_Standardising$Spec)[order(unique(TYB_Standardising$Spec))]) #Make the species factors (to allow split, next line)
TYB_Standardising <- split(TYB_Standardising, f=TYB_Standardising$Spec)

RandomNumSpec <- lapply(TYB_Names_Cast_50$., function(x) sample(1:x, 50, replace=FALSE)) #Get 50 random numbers sampled from the total number of sites for each species
TYB_St <- rbindlist(lapply(1:length(TYB_Standardising), function(x) TYB_Standardising[[x]][RandomNumSpec[[x]],]))$SiteSpec #Sample from all possible sitespec for each species, using the random numbers
save(TYB_St, file=paste0(ResultsFP, "TYB_St.RData")) #Save

#### Prepare generation length data ####
SiteSpec <- as.data.frame(names(TYB_list)) #Get all the sitespec
SiteSpec$Species <- str_split_fixed(SiteSpec$'names(TYB_list)', "[_]", 3)[,2] #Extract just species names
names(SiteSpec) <- c("SiteSpec", "Species") #Rename

Genl <- read.csv(paste0(ResultsFP, "GenLengths.csv")) #Pull in generation length data
Genl <- subset(Genl, GenerationLength!="NotGiven") #Remove any species where generation length wasn't given
Genl$GenerationLength <- as.numeric(as.character(Genl$GenerationLength)) #Class gen lengths as numeric
GenlAll <- merge(SiteSpec, Genl, by="Species") #Add generation length data to each sitespec

Genl1_5 <- subset(GenlAll, GenerationLength<6)
Genl1_5$GenLengthMax <- 5
Genl6_10 <- subset(GenlAll, GenerationLength>6 & GenerationLength<11)
Genl6_10$GenLengthMax <- 10
Genl11_15 <- subset(GenlAll, GenerationLength>11 & GenerationLength<16)
Genl11_15$GenLengthMax <- 15

SampleSize <- min(nrow(Genl1_5), nrow(Genl6_10), nrow(Genl11_15)) #Find the category with the smallest number of sitespec, and randomly reduce the others to this length (to make sure we're not unfairly comparing)

Genl1_5 <- Genl1_5[sample(1:nrow(Genl1_5), SampleSize),] #Reduce each of the categories to the smallest
Genl6_10 <- Genl6_10[sample(1:nrow(Genl6_10), SampleSize),] #Reduce each of the categories to the smallest
Genl11_15 <- Genl11_15[sample(1:nrow(Genl11_15), SampleSize),] #Reduce each of the categories to the smallest

GenLengthSpec <- rbind(Genl1_5, Genl6_10, Genl11_15)

write.csv(GenLengthSpec, paste0(ResultsFP, "GenLengthSpec.csv"))

#### Functions. Write functions ####
#These functions take the input data from the run models, with a slope and significance level of each population, with each combination of consecutive and interval samplings COMPARED TO ONE value of complete trend lengths (run multiple times for each complete trend length)
#e.g. slope and significance of population x when sampled for 10 years starting at year 4 compared to slope and significance of population x when sampled for 20 years starting at year 3
#Direction function returns the number of samples that are matching, an erroneous positive/negative, missed positive/negative or opposing when compared to the complete trend
#Magnitude function returns the number of samples that are matching according to different tolerances when comparing the slope of the sample to the complete trend
#If Standardise is TRUE, it will subset the input data to the standardised dataset (98 species with 50 randomly selected sites each)
#If GenLength == 0, use all data. If it is 5 the generation length = 5 or less, if 10, = 6-10, if 15, = 11-15
#If EDFNum == 0, use all data. Otherwise 1 == EDF 1,2; 3 == EDF 3,4; 5 == EDF 5,6; 7== EDF 7,8.
#Significance sets the p value cutoff (0.05 for all published results)

Direction <- function(DATA, Standardise, GenLength, EDFNum, Significance){
  Input <- DATA
  SamplingType <- unique(Input$SamplingType)
  #Reduce the data according to the various subsets
  if(GenLength==0){
    if(Standardise==TRUE){Input <- Input[Input$SiteSpec %in% TYB_St,]}
  } else {
    GenLengthSub <- subset(GenLengthSpec, GenLengthMax==GenLength)
    Input <- Input[Input$SiteSpec %in% GenLengthSub$SiteSpec,]
  } 
  
  if(EDFNum!=0){
    Input <- DATA
    Input[is.na(Input$EDFCompleteLength),]$EDFCompleteLength <- 0
    Input <- subset(Input, EDFCompleteLength<EDFNum)
  }
  
  if(nrow(Input)==0){
    return(NULL)
  }
  
  #Change the slope to just positive, negative or insignificant
  Input$Slope <- sapply(Input$Slope, function (x) if(is.na(x)){"Insignificant"} else if(x > 0){ "Positive" } else{ "Negative" })
  Input$CompleteLengthSlope <- sapply(Input$CompleteLengthSlope, function (x) if(is.na(x)){"Insignificant"} else if(x > 0){ "Positive" } else{ "Negative" })
  
  #Create a binary value for significance based on p value
  Input$Significant <- sapply(Input$Significant, function (x) if(is.na(x)){0} else if(x < Significance){1} else{0})
  Input$SignificantCompleteLength <- sapply(Input$SignificantCompleteLength, function (x) if(is.na(x)){0} else if(x < Significance){1} else{0})
  
  #Change the slope to insignificant where relevant
  Input[Input$Significant==0]$Slope <- "Insignificant"
  Input[Input$SignificantCompleteLength==0]$CompleteLengthSlope <- "Insignificant"
  Input <- Input[,c("SiteSpec", "NumYears", "Slope", "CompleteLengthSlope")]
  
  #Calculate the erroneous +/-, missed +/-, matching and opposing
  MPos <- subset(Input, CompleteLengthSlope=="Positive")
  MPos$CompleteLengthSlope <- NULL
  MPos$Slope <- recode_factor(MPos$Slope, "Positive"="Matching", "Negative" = "Opposing", "Insignificant" = "Missed Positive")
  
  MNeg <- subset(Input, CompleteLengthSlope=="Negative")
  MNeg$CompleteLengthSlope <- NULL
  MNeg$Slope <- recode_factor(MNeg$Slope, "Positive"="Opposing", "Negative" = "Matching", "Insignificant" = "Missed Negative")

  Err <- subset(Input, CompleteLengthSlope=="Insignificant")
  Err$CompleteLengthSlope <- NULL
  Err$Slope <- recode_factor(Err$Slope, "Positive" = "Erroneous Positive", "Negative" = "Erroneous Negative", "Insignificant" = "Matching")
  
  summarise <- rbind(Err, MPos, MNeg) 

  rm(Input)
  
  #Cast to find number of populations of each category
  summarise <- dcast(summarise, NumYears~Slope, length, value.var="Slope")
  summarise <- melt(summarise, id.vars="NumYears")
  
  #Split numyears for interval
  if(SamplingType=="Interval"){
    summarise[,c("IntervalLength", "NumYears")] <- str_split_fixed(summarise$NumYears, "[.]", 2)
  }
  
  #Order by number of years, add metadata
  summarise$NumYears <- as.numeric(as.character(summarise$NumYears))
  summarise <- summarise[order(-summarise$NumYears),]
  summarise$SamplingType <- SamplingType
  summarise$Model <- modelfam
  if(SamplingType=="Consecutive"){
    summarise$CompleteLength <- unique(DATA$CompleteLengthNum)
  }
  summarise$GenLength <- GenLength
  if(!Standardise){
    summarise$CastType <- "All"
  } else{
    summarise$CastType <- "St"
  }
  summarise$EDF <- EDFNum
  return(summarise)
}
Magnitude <- function(DATA, Standardise, GenLength, EDFNum, Significance){
  Input <- DATA
  SamplingType <- unique(Input$SamplingType)
  
  #Reduce the data according to the various subsets
  
  if(GenLength==0){
    if(Standardise==TRUE){Input <- Input[Input$SiteSpec %in% TYB_St,]}
  } else {
    GenLengthSub <- subset(GenLengthSpec, GenLengthMax==GenLength)
    Input <- Input[Input$SiteSpec %in% GenLengthSub$SiteSpec,]
  } 
  
  if(EDFNum!=0){
    Input <- DATA
    Input[is.na(Input$EDFCompleteLength),]$EDFCompleteLength <- 0
    Input <- subset(Input, EDFCompleteLength<EDFNum)
  }
  

  Input <- subset(Input, Significant<Significance & SignificantCompleteLength<Significance)
  
  if(nrow(Input)==0){
    return(NULL)
  }
  
  if(SamplingType=="Consecutive"){
    Input[,c("SamplingType", "Model", "CompleteLengthNum", "EDF", "EDFCompleteLength")] <- NULL
  } else{
    Input <- Input[,c("SiteSpec","NumYears", "Slope", "CompleteLengthSlope")]
    Input <- Input[!is.na(Input$Slope),]
  }
  
  Input <- Input[complete.cases(Input)] #Remove NAs
  
  #Find correct and incorrect for the various tolerances
  summarise <- rbindlist(lapply(c(0.01, 0.025, 0.05, 0.1, 0.25, 0.5), function(x){
    ok <- Input
    ok$Trend <- ifelse(Input$Slope<(Input$CompleteLengthSlope+x) & Input$Slope>(Input$CompleteLengthSlope-x), "Correct", "Incorrect")
    ok$Tolerance <- x
    summarise <- dcast(ok, NumYears+Tolerance~Trend, length, value.var="Trend")
    return(summarise)
  }))
  rm(Input)
  
  #Split numyears for interval
  if(SamplingType=="Interval"){
    summarise[,c("IntervalLength")] <- str_split_fixed(summarise$NumYears, "[.]", 2)[,1]
    summarise[,c("NumYears")] <- str_split_fixed(summarise$NumYears, "[.]", 2)[,2]
  }
  
  #Add metadata
  summarise$NumYears <- as.numeric(as.character(summarise$NumYears))
  summarise$SamplingType<- SamplingType
  summarise$Model <- modelfam
  summarise$CompleteLength <- unique(DATA$CompleteLengthNum)
  summarise$GenLength <- GenLength
  if(!Standardise){
    summarise$CastType <- "All"
  } else {
    summarise$CastType <- "St"
  } 
  summarise$EDF <- EDFNum
  return(summarise)
}

#### Models 1. Consecutive ####
#Run models
Consecutive <- rbindlist(lapply(3:MCL, function(CompleteLength){ #This apply runs through the number of years of each model run (3 years to MCL years)
  StartYear <- pbmclapply(1:((MCL+1)-CompleteLength), function(StartYear){ #And this one runs through the possible start years (so e.g. CompleteLength=5, StartYear=10 would mean a model running on 5 years of data, taken from years 10-14 for every SiteSpec)
    if(file.exists(paste0(ResultsFP,"GLM/Consecutive/Consecutive_GLM_", CompleteLength, "Years_",StartYear, "StartYear_", modelfam, ".csv"))==TRUE){ #Because these models take a long time to run code could run on multiple clusters. This means anything that's already done is not done again
      return(NULL)
    } else {
      write.csv(NULL, paste0(ResultsFP,"GLM/Consecutive/Consecutive_GLM_", CompleteLength, "Years_",StartYear, "StartYear_", modelfam, ".csv"), row.names=FALSE)
    }
    Degraded <- lapply(TYB_list, function(x) x[StartYear:(StartYear+CompleteLength-1),]) #Cut the counts down to the correct number of years (CompleteLength), starting at the right year (StartYear)
    Degraded2 <- lapply(Degraded, function(x) ifelse(sum(x$Count)==0, return(NULL), return(x))) #Turn all entries with 0 at all times to NULL (no point running a model on them)
    Degraded2 <- Degraded2[!sapply(Degraded2, is.null)] #Remove null elements
    
    if(modelfam=="quasi"){ #Run the glm, using the model family defined (for the paper, model family is negative binomial - "nb")
      theglmlist <- lapply(Degraded2, function (x) glm(Count~Year, offset=log(x$Hours), quasipoisson(link="log"), data=x))
    } else if(modelfam=="pois"){
      theglmlist <- lapply(Degraded2, function (x) glm(Count~Year, offset=log(x$Hours), poisson(link="log"), data=x))
    } else {
      theglmlist <- lapply(Degraded2, function (x) tryCatch(glm.nb(Count~Year + offset(log(x$Hours)), link=log, data=x), error=function(e){NULL}))
    }

    thegamlist <- lapply(Degraded2, function (x) tryCatch(gam(Count~s(Year) + offset(log(Hours)), family = quasipoisson(link='log'), data=x), error=function(e){NULL})) #Also run a GAM on the model, to determine the curve shape (using estimate degrees of freedom, EDF). 
    #Nb due to issues with running negative binomial GAMs with sparse data, this uses a quasipoisson distribution. Checks indicated high convergence between the two model families for getting EDF. 
    
    theglmlist <- theglmlist[!sapply(theglmlist, is.null)] #Remove nulls 
    thegamlist <- thegamlist[!sapply(thegamlist, is.null)] #Remove nulls
    
    slopes <- sapply(theglmlist, function (x) tryCatch(coef(x, silent=TRUE)[[2]], error=function(e){NULL})) #Extract the coefficient of slope (i.e. r)
    slopes <- slopes[!sapply(slopes, is.null)] #Remove nulls, organise data
    slopesnames <- names(slopes)
    slopes <- as.data.frame(unlist(slopes))
    slopes$Data <- slopesnames
    
    significantglm <- sapply(theglmlist, function (x) tryCatch(summary(x)$coeff[-1,4], error=function(e){NULL})) #Extract the significance of the slope coefficient (i.e. p)
    significantglm <- significantglm[!sapply(significantglm, is.null)] #Remove nulls, organise data
    significantglmnames <- names(significantglm)
    significantglm <- as.data.frame(unlist(significantglm))
    significantglm$Data <- significantglmnames
    
    edfgam <- sapply(thegamlist, function (x) tryCatch(summary(x)$s.table[[1]], error=function(e){NA})) #Extract the estimated degrees of freedom from the GAM (EDF)
    edfgam <- edfgam[!sapply(edfgam, is.null)] #Remove nulls, organise data
    edfgamnames <- names(edfgam)
    edfgam <- as.data.frame(unlist(edfgam))
    edfgam$Data <- edfgamnames
    
    if(nrow(significantglm)!=0 & nrow(slopes)!=0){ #Providing not all models have returned null, create final dataset
      finaldata <- merge(slopes, significantglm, by="Data")
      if(nrow(edfgam)>0){ #Sometimes this is zero, because no GAMs were able to run on the data (often because data only 3 years in length)
        finaldata <- merge(finaldata, edfgam, by="Data", all=TRUE)
      } else {
        finaldata$edfgam <- NA
      }
      names(finaldata) <- c("SiteSpec", "Slope", "Significant", "EDF")
    } else {
      finaldata <- data.frame("SiteSpec"=NULL, "Slope"=NULL, "Significant"=NULL, "EDF"=NULL)
    }
    
    #And now add in the sitespec where models didn't run and/or there were zeros in all years (and therefore conclusion is insignificant)
    if(nrow(finaldata) != length(Degraded)){ #I.e. if there are some records without data
      InsigSiteSpec <- as.data.frame(names(Degraded)[!names(Degraded) %in% finaldata$SiteSpec]) #Extract these records
      names(InsigSiteSpec) <- "SiteSpec" #Define records as NA on all counts
      InsigSiteSpec$Slope <- NA
      InsigSiteSpec$Significant <- NA
      InsigSiteSpec$EDF <- NA
      finaldata <- rbind(finaldata, InsigSiteSpec) #Add them to final dataset
    }
    
    finaldata$NumYears <- as.numeric(as.character(CompleteLength)) #Add metadata to final dataset 
    finaldata$StartYear <- as.numeric(as.character(StartYear))
    finaldata$SamplingType<- "Consecutive"
    finaldata$Model <- modelfam
    write.csv(finaldata, paste0(ResultsFP,"GLM/Consecutive/Consecutive_GLM_", CompleteLength, "Years_",StartYear, "StartYear_", modelfam, ".csv"), row.names=FALSE) #Write out results
    return(finaldata)
  }, mc.cores=ncores)
  return(rbindlist(StartYear))
}))

#Summarise
Consecutive <- rbindlist(pblapply(list.files(path=paste0(ResultsFP, "GLM/Consecutive/"), pattern=paste0("*", modelfam, ".csv"), full.names=TRUE), fread), fill=TRUE) #Pull the data from the above back together (assuming it has taken multiple runs to complete, otherwise the result will just be taken from the function above)
#The following function organises the data so that samples can be compared to multiple complete trend lengths. E.g. A complete trend 10 years long taken from years 10-19 could have 3 year samples taken from it starting from years 10,11,12...17 etc)
#It then compares sample trend to complete trend according to the two comparison techniques (Direction and Magnitude)
CastConsecutive <- pblapply(c(4:MCL), function(CompleteLength){
  if((file.exists(paste0(ResultsFP, "GLM/Summaries/Direction_Consecutive_", modelfam, "_", CompleteLength,".csv"))&
      file.exists(paste0(ResultsFP, "GLM/Summaries/Magnitude_Consecutive_", modelfam, "_", CompleteLength,".csv")))==TRUE){
    return(NULL)
  } else {
    write.csv(NULL, paste0(ResultsFP, "GLM/Summaries/Direction_Consecutive_", modelfam, "_", CompleteLength,".csv"))
    write.csv(NULL, paste0(ResultsFP, "GLM/Summaries/Magnitude_Consecutive_", modelfam, "_", CompleteLength,".csv"))

    #Work down through the years, removing the start years that don't fit (e.g. start yr at yr21 when we're only going up to 20now)
    CompleteLengthSub <- Consecutive
    CompleteLengthSub[,c("SamplingType", "Model")] <- NULL
    CompleteLengthSlopes <- subset(CompleteLengthSub, NumYears==CompleteLength)
    names(CompleteLengthSlopes) <- c("SiteSpec","CompleteLengthSlope", "SignificantCompleteLength", "EDFCompleteLength", "CompleteLengthNum", "CompleteLengthStartYear")
    
    SamplesSub <- subset(CompleteLengthSub, NumYears<CompleteLength)
    rm(CompleteLengthSub)
    AllTheStartYears <- rbindlist(pblapply(unique(CompleteLengthSlopes$CompleteLengthStartYear), function(x){
      CompleteLengthSubStartYear <- subset(CompleteLengthSlopes, CompleteLengthStartYear==x)
      StartYearSub <- SamplesSub
      StartYearSub <- subset(StartYearSub, StartYear > (x-1))
      StartYearSub$Keep <- ifelse((StartYearSub$StartYear+(StartYearSub$NumYears)-1) < (x+CompleteLength), 1, 0)
      StartYearSub <- subset(StartYearSub, Keep==1)
      StartYearMerge <- merge(StartYearSub, CompleteLengthSubStartYear, All=T)
      return(StartYearMerge)
    }))
    rm(CompleteLengthSlopes, SamplesSub)
    AllTheStartYears$Keep <- NULL
    AllTheStartYears$Model <- modelfam
    AllTheStartYears$SamplingType <- "Consecutive"
    
    Direction_All <- rbindlist(pblapply(c(0,5,10,15), function(x) Direction(AllTheStartYears, Standardise=FALSE, GenLength=x, EDFNum=0, Significance = 0.05)))
    Direction_St <- rbindlist(lapply(c(0,5,10,15), function(x) Direction(AllTheStartYears, Standardise=TRUE, GenLength=x, EDFNum=0, Significance = 0.05)))
    Direction_EDF <- rbindlist(pblapply(c(3,5,7,9), function(x) Direction(AllTheStartYears, Standardise=FALSE, GenLength=0, EDFNum=x, Significance = 0.05)))
    write.csv(rbind(Direction_All, Direction_St, Direction_EDF), file=paste0(ResultsFP, "GLM/Summaries/Direction_Consecutive_", modelfam, "_", CompleteLength,".csv"), row.names=FALSE)
    
    Magnitude_All <- rbindlist(pblapply(c(0,5,10,15), function(x) Magnitude(AllTheStartYears, Standardise=FALSE, GenLength=x, EDFNum=0, Significance = 0.05)))
    Magnitude_St <- rbindlist(pblapply(c(0,5,10,15), function(x) Magnitude(AllTheStartYears, Standardise=TRUE, GenLength=x, EDFNum=0, Significance = 0.05)))
    Magnitude_EDF <- rbindlist(pblapply(c(3,5,7,9), function(x) Magnitude(AllTheStartYears, Standardise=FALSE, GenLength=0, EDFNum=x, Significance = 0.05)))
    write.csv(rbind(Magnitude_All, Magnitude_St, Magnitude_EDF), file=paste0(ResultsFP, "GLM/Summaries/Magnitude_Consecutive_", modelfam, "_", CompleteLength,".csv"), row.names=FALSE)
  }
})

#### Models 2. Intervals ####
#Run Models
#Create a list of all the ways MCL years can be sampled in intervals (all possible number of years and space between years)
Intervals <- unlist(unlist(lapply(1:(MCL-2), function(x){
  lapply(2:((MCL/2) + 1), function(y){
    ok <- seq(from = x, to = MCL, by = y)
    if(length(ok)>2){
      ok2 <- lapply(3:(length(ok)), function(z){
        ok[1:z]
      })
      return(ok2)
    } else{return(NULL)}
  })
}), recursive = FALSE), recursive = FALSE)

if(!file.exists(file=paste0(ResultsFP, "GLM/Intervals/Intervals_", modelfam, ".csv"))){
  Interval <- rbindlist(pbmclapply(Intervals, function(IntValues){
    Degraded <- lapply(TYB_list, function(x) x[IntValues,]) #Cut the counts down to the right intervals
    Degraded2 <- lapply(Degraded, function(x) ifelse(sum(x$Count)==0, return(NULL), return(x))) #Remove entries with 0 at all times
    Degraded2 <- Degraded2[!sapply(Degraded2, is.null)]
    
    if(modelfam=="quasi"){ #Run the glm, using the model family defined (for the paper, model family is negative binomial - "nb")
      theglmlist <- lapply(Degraded2, function (x) glm(Count~Year, offset=log(x$Hours), quasipoisson(link="log"), data=x))
    } else if(modelfam=="pois"){
      theglmlist <- lapply(Degraded2, function (x) glm(Count~Year, offset=log(x$Hours), poisson(link="log"), data=x))
    } else {
      theglmlist <- lapply(Degraded2, function (x) tryCatch(glm.nb(Count~Year + offset(log(x$Hours)), link=log, data=x), error=function(e){NULL}))
    }
    
    theglmlist <- theglmlist[!sapply(theglmlist, is.null)] #Remove nulls
    
    slopes <- sapply(theglmlist, function (x) tryCatch(coef(x, silent=TRUE)[[2]], error=function(e){NULL}))  #Extract the coefficient of slope (i.e. r)
    slopes <- slopes[!sapply(slopes, is.null)] #Remove nulls and organise data
    slopes <- as.data.frame(unlist(slopes))
    slopes$Data <- rownames(slopes)
    
    significantglm <- sapply(theglmlist, function (x) tryCatch(summary(x)$coeff[-1,4], error=function(e){NULL})) #Extract the significance of the slope coefficient (i.e. p)
    significantglm <- significantglm[!sapply(significantglm, is.null)] #Remove nulls and organise data
    significantglm <- as.data.frame(unlist(significantglm))
    significantglm$Data <- rownames(significantglm)
    
    if(nrow(significantglm)!=0 & nrow(slopes)!=0){ #Providing not all models have returned null, create final dataset
      finaldata <- merge(slopes, significantglm, by="Data")
      names(finaldata) <- c("SiteSpec", "Slope", "Significant")
    } else {
      finaldata <- data.frame("SiteSpec"=NULL, "Slope"=NULL, "Significant"=NULL)
    }
    
    #And now add in the sitespec that are insignificant
    if(nrow(finaldata) != length(Degraded)){ 
      InsigSiteSpec <- as.data.frame(names(Degraded)[!names(Degraded) %in% finaldata$SiteSpec])
      names(InsigSiteSpec) <- "SiteSpec"
      InsigSiteSpec$Slope <- NA
      InsigSiteSpec$Significant <- NA
      finaldata <- rbind(finaldata, InsigSiteSpec)
    }
    
    #Add metadata
    IntervalLength <- unique(sapply(1:(length(IntValues)-1), function(x) IntValues[x+1]-IntValues[x]))  
    finaldata$IntervalLength <- IntervalLength
    finaldata$StartYear <- IntValues[1]
    finaldata$NumYears <- length(IntValues)
    finaldata$Model <- modelfam
    finaldata$SamplingType <- "Interval"
    return(finaldata)
  }, mc.cores=ncores))
  write.csv(Interval, file=paste0(ResultsFP, "GLM/Intervals/Intervals_", modelfam, ".csv"), row.names=FALSE)
}

#Summarise
if((file.exists(paste0(ResultsFP, "GLM/Summaries/Direction_Interval_", modelfam, ".csv")) &
    file.exists(paste0(ResultsFP, "GLM/Summaries/Magnitude_Interval_", modelfam, ".csv")))==FALSE){
  write.csv(NULL, paste0(ResultsFP, "GLM/Summaries/Direction_Interval_", modelfam, ".csv"))
  write.csv(NULL, paste0(ResultsFP, "GLM/Summaries/Magnitude_Interval_", modelfam, ".csv"))

  Interval <- fread(paste0(ResultsFP, "GLM/Intervals/Intervals_", modelfam, ".csv"))
  CompleteTrend <- fread(paste0(ResultsFP, "GLM/Consecutive/Consecutive_GLM_", MCL, "Years_1StartYear_nb.csv"))[,c("SiteSpec", "Significant", "Slope", "EDF", "NumYears")] #Read in the models run on full 30 years of data (for comparison to complete)
  names(CompleteTrend) <- c("SiteSpec", "SignificantCompleteLength", "CompleteLengthSlope" , "EDFCompleteLength", "CompleteLengthNum")
  Interval <- merge(Interval, CompleteTrend, by="SiteSpec") #Merge complete MCL years with the interval runs
  Interval$NumYears <- paste0(Interval$IntervalLength, ".", Interval$NumYears) #Pull these two together to enable the intervals to run using the functions
  
  Direction_All <- rbindlist(pblapply(c(0,5,10,15), function(x) Direction(Interval, Standardise=FALSE, GenLength=x, EDFNum=0, Significance = 0.05)))
  Direction_St <- rbindlist(pblapply(c(0,5,10,15), function(x) Direction(Interval, Standardise=TRUE, GenLength=x, EDFNum=0, Significance = 0.05)))
  Direction_EDF <- rbindlist(pblapply(c(3,5,7,9), function(x) Direction(Interval, Standardise=FALSE, GenLength=0, EDFNum=x, Significance = 0.05)))

  write.csv(rbind(Direction_All, Direction_St, Direction_EDF), file=paste0(ResultsFP, "GLM/Summaries/Direction_Interval_", modelfam, ".csv"), row.names=FALSE)
  
  Magnitude_All <- rbindlist(pblapply(c(0,5,10,15), function(x) Magnitude(Interval, Standardise=FALSE, GenLength=x, EDFNum=0, Significance = 0.05)))
  Magnitude_St <- rbindlist(pblapply(c(0,5,10,15), function(x) Magnitude(Interval, Standardise=TRUE, GenLength=x, EDFNum=0, Significance = 0.05)))
  Magnitude_EDF <- rbindlist(pblapply(c(3,5,7,9), function(x) Magnitude(Interval, Standardise=FALSE, GenLength=0, EDFNum=x, Significance = 0.05)))

  write.csv(rbind(Magnitude_All, Magnitude_St, Magnitude_EDF), file=paste0(ResultsFP, "GLM/Summaries/Magnitude_Interval_", modelfam, ".csv"), row.names=FALSE)
}

#### Plot #### 
plotfontsize <- 10 #Set the font size for plots

#The general scaffolding of these plots could be used for one's own data, however a lot of specific parameters would need to change so bear that in mind. 

#This function gives us the proportion of samples that fell into each of the Direction categories (for each complete length, subset length and interval period)
#The models sections return the raw number of samples in each category, but not the proportion, so we need to calculate
PropOverall <- function(Direction, Category){
  Direction2 <- if(SamplingType=="Consecutive"){split(Direction, Direction$CompleteLength)}else{split(Direction, Direction$NumYears)} #Split into a list of dataframes, one for each Complete length (consecutive sampling) or number of years sampled (interval sampling)
  DataFrame <- rbindlist(lapply(Direction2, function(y){
    okcast <- if(SamplingType=="Consecutive"){
      dcast(y, NumYears~variable, value.var="value") #Cast so that the Direction categories are put as columns, with the length of samples (Numyears) as the rows, and cell values the number of populations
    } else {
      dcast(y, IntervalLength~variable, value.var="value") #If interval sampling, cast so that interval lengths are the rows
    }
    if(Category=="Matching"){ #The following 'if elses' calculate the proportion of populations of each category, by finding (number of category x)/(number in all categories)
      okcast$Prop <- sapply(1:nrow(okcast), function(x) okcast$Matching[x]/sum(okcast[x,2:ncol(okcast)]))
      okcast$Category <- "a) Matching"
    } else if(Category=="ErrPos"){
      okcast$Prop <- sapply(1:nrow(okcast), function(x) okcast$'Erroneous Positive'[x]/sum(okcast[x,2:ncol(okcast)]))
      okcast$Category <- "d) Erroneous Negative"
    } else if(Category=="ErrNeg"){
      okcast$Prop <- sapply(1:nrow(okcast), function(x) okcast$'Erroneous Negative'[x]/sum(okcast[x,2:ncol(okcast)]))
      okcast$Category <- "c) Erroneous Positive"
    } else if(Category=="MPos"){
      okcast$Prop <- sapply(1:nrow(okcast), function(x) okcast$'Missed Positive'[x]/sum(okcast[x,2:ncol(okcast)]))
      okcast$Category <- "f) Missed Negative"
    } else if(Category=="MNeg"){
      okcast$Prop <- sapply(1:nrow(okcast), function(x) okcast$'Missed Negative'[x]/sum(okcast[x,2:ncol(okcast)]))
      okcast$Category <- "e) Missed Positive"
    } else {
      okcast$Prop <- sapply(1:nrow(okcast), function(x) okcast$Opposing[x]/sum(okcast[x,2:ncol(okcast)]))
      okcast$Category <- "b) Opposing"
    }
    if(SamplingType=="Consecutive"){
      okcast <- okcast[,c("NumYears", "Prop", "Category")] #Cut dataset down to relevant columns
      okcast$CompleteLength <- unique(y$CompleteLength) #Add complete length into the dataset
      okcast$NumYears <- as.numeric(as.character(okcast$NumYears))
    } else {
      okcast <- okcast[,c("IntervalLength", "Prop", "Category")] #Cut dataset down to relevant columns
      okcast$NumYears <- unique(y$NumYears) #Add complete length into the dataset
      okcast$IntervalLength <- as.numeric(as.character(okcast$IntervalLength))
    }
    okcast$Prop <- as.numeric(okcast$Prop)
    okcast[is.na(okcast)] <- 0 #Change NAs to zeros
    return(okcast)
  }))
  return(DataFrame)
}

#This function gives us the proportion of samples that fell into each of the Direction categories (for each complete length, subset length and interval period), combining Missed positive and negative into one term, ditto for Erroneous positive and negative
PropOverallCombine <- function(Direction, Category){
  Direction2 <- if(SamplingType=="Consecutive"){split(Direction, Direction$CompleteLength)}else{split(Direction, Direction$NumYears)} #Split into a list of dataframes, one for each Complete length (consecutive sampling) or number of years sampled (interval sampling)
  DataFrame <- rbindlist(lapply(Direction2, function(y){
    okcast <- if(SamplingType=="Consecutive"){
      dcast(y, NumYears~variable, value.var="value") #Cast so that the Direction categories are put as columns, with the length of samples (Numyears) as the rows, and cell values the number of populations
    } else {
      dcast(y, IntervalLength~variable, value.var="value") #If interval sampling, cast so that interval lengths are the rows
    }
    if(Category=="Matching"){ #The following 'if elses' calculate the proportion of populations of each category, by finding (number of category x)/(number in all categories)
      okcast$Prop <- sapply(1:nrow(okcast), function(x) okcast$Matching[x]/sum(okcast[x,2:ncol(okcast)]))
      okcast$Category <- "a) Matching"
    } else if(Category=="Erroneous"){
      AllErroneous <- if("Erroneous Positive" %in% colnames(okcast)){
        if("Erroneous Negative" %in% colnames(okcast)){
          okcast$`Erroneous Negative` + okcast$`Erroneous Positive`
        } else {
          okcast$`Erroneous Positive`
        }
      } else {
        okcast$`Erroneous Negative`
      }
      okcast$Prop <- sapply(1:length(AllErroneous), function(x) AllErroneous[x]/sum(okcast[x,2:ncol(okcast)]))
      okcast$Category <- "c) Erroneous +/-"
    } else if(Category=="Missed"){
      AllMissed <- if("Missed Positive" %in% colnames(okcast)){
        if("Missed Negative" %in% colnames(okcast)){
          okcast$`Missed Negative` + okcast$`Missed Positive`
        } else {
          okcast$`Missed Positive`
        }
      } else {
        okcast$`Missed Negative`
      }
      okcast$Prop <- sapply(1:length(AllMissed), function(x) AllMissed[x]/sum(okcast[x,2:ncol(okcast)]))
      okcast$Category <- "d) Missed +/-"
    } else {
      okcast$Prop <- sapply(1:nrow(okcast), function(x) okcast$Opposing[x]/sum(okcast[x,2:ncol(okcast)]))
      okcast$Category <- "b) Opposing"
    }
    if(SamplingType=="Consecutive"){
      okcast <- okcast[,c("NumYears", "Prop", "Category")] #Cut dataset down to relevant columns
      okcast$CompleteLength <- unique(y$CompleteLength) #Add complete length into the dataset
      okcast$NumYears <- as.numeric(as.character(okcast$NumYears))
    } else {
      okcast <- okcast[,c("IntervalLength", "Prop", "Category")] #Cut dataset down to relevant columns
      okcast$NumYears <- unique(y$NumYears) #Add complete length into the dataset
      okcast$IntervalLength <- as.numeric(as.character(okcast$IntervalLength))
    }
    okcast$Prop <- as.numeric(okcast$Prop)
    okcast[is.na(okcast)] <- 0 #Change NAs to zeros
    return(okcast)
  }))
  return(DataFrame)
}

#General plot function for direction data
DirectionPlot <- function(Direction){ 
  Direction$Prop <- round_any(Direction$Prop*100, 10, floor) #Convert proportions to percentages and round down to nearest 10
  Direction$Prop <- paste0(Direction$Prop, "-", Direction$Prop+10) #Create label (e.g. "0-10" rather than just "0")
  Direction$Prop <- factor(Direction$Prop, levels=sapply(seq(0,90,10), function(x) paste0(x, "-", x+10))) #Convert to factor
  
  if(SamplingType=="Consecutive"){ #Create labels (adjusts based on consecutive or interval sampling)
    xlabel <- "Complete trend length (years)"
    ylabel <- "Sample trend length (years)"
    xbreaks <- (c(seq(4,MCL,4)))
    ybreaks <- (c(seq(3,MCL,3)))
    ex <- Direction$CompleteLength
    why <- Direction$NumYears
    aspectrat <- 1
  } else {
    xlabel <- "Number of years sampled"
    ylabel <- "Samples taken every x years"
    xbreaks <- (c(seq(3,(MCL-1),5)))
    ybreaks <- (c(seq(1,(MCL/2),3)))
    aspectrat <- 0.52
    ex <- Direction$NumYears
    why <- Direction$IntervalLength
  }
  
  ggplot(Direction, aes(x=ex,y=why))+
    geom_tile(aes(fill=Prop), colour = "grey40",size = 0.01)+ #Tile makes the nice squares
    facet_wrap(~Category, nrow=2)+ #Add a facet for each direction category
    ylab(ylabel)+
    xlab(xlabel)+
    scale_fill_manual(values = viridis(n=10), name=paste0("Percentage"), guide=guide_legend(reverse = TRUE, keywidth=unit(3, "mm"), keyheight = unit(3, "mm")), drop=FALSE)+ #This sets the legend as the same regardless of which colours appear (it always shows all options from 0-100)
    scale_x_continuous(breaks=xbreaks, expand = c(0,0.15))+
    scale_y_continuous(breaks=ybreaks, expand = c(0,0.15))+
    theme(aspect.ratio=aspectrat, 
          panel.background = element_blank(),
          panel.grid = element_blank(), 
          text = element_text(size=plotfontsize),
          panel.border = element_rect(size = 1, fill = NA),
          strip.background = element_blank(),
          strip.text = element_text(hjust = 0, size=plotfontsize),
          legend.title=element_text(size=plotfontsize),
          axis.text = element_text(size=plotfontsize, colour="black"),
          axis.ticks = element_line(colour="black"),
          legend.text = element_text(size=plotfontsize, colour="black"),
          legend.justification = "top")
}
#Plot function for direction data split by generation length
DirectionPlotGen <- function(Direction){
  Direction$Prop <- round_any(Direction$Prop*100, 10, floor) #Convert proportions to percentages and round down to nearest 10
  Direction$Prop <- paste0(Direction$Prop, "-", Direction$Prop+10) #Create label (e.g. "0-10" rather than just "0")
  Direction$Prop <- factor(Direction$Prop, levels=sapply(seq(0,90,10), function(x) paste0(x, "-", x+10))) #Convert to factor
  
  if(SamplingType=="Consecutive"){ #Create labels (adjusts based on consecutive or interval sampling)
    xlabel <- "Complete trend length (years)"
    ylabel <- "Sample trend length (years)"
    xbreaks <- (c(seq(4,MCL,6)))
    ybreaks <- (c(seq(3,MCL,6)))
    aspectrat <- 1
    ex <- Direction$CompleteLength
    why <- Direction$NumYears
  } else {
    xlabel <- "Number of years sampled"
    ylabel <- "Samples taken every x years"
    xbreaks <- (c(seq(3,29,6)))
    ybreaks <- (c(seq(0,14,3)))
    aspectrat <- 0.52
    ex <- Direction$NumYears
    why <- Direction$IntervalLength
  }
  
  Direction$GenLength <- as.character(as.numeric(Direction$GenLength)) #This block adjusts all the names so the plot works
  Direction[Direction$GenLength==5,]$GenLength <- "a) Short"
  Direction[Direction$GenLength==10,]$GenLength <- "b) Medium"
  Direction[Direction$GenLength==15,]$GenLength <- "c) Long"
  Direction[Direction$Category=="a) Matching",]$Category <- "Matching"
  Direction[Direction$Category=="c) Erroneous +/-",]$Category <- "Erroneous +/-"
  Direction[Direction$Category=="d) Missed +/-",]$Category <- "Missed +/-"
  Direction[Direction$Category=="b) Opposing",]$Category <- "Opposing"
  Direction$GenLength <- factor(Direction$GenLength)
  Direction$GenLength <- factor(Direction$GenLength,levels(Direction$GenLength)[c(1,2,3)])
  Direction$Category <- factor(Direction$Category)
  Direction$Category <- factor(Direction$Category,levels(Direction$Category)[c(2,4,1,3)])
  
  ggplot(Direction, aes(x=ex,y=why))+
    geom_tile(aes(fill=Prop), colour = "grey40",size = 0.01)+
    facet_grid(Category~GenLength)+ #Make the facets show category vs. generation length
    ylab(ylabel)+
    xlab(xlabel)+
    scale_fill_manual(values = viridis(n=10), name=paste0("Percentage"), guide=guide_legend(reverse = TRUE, keywidth=unit(3, "mm"), keyheight = unit(3, "mm")), drop=FALSE)+ #This sets the legend as the same regardless of which colours appear (it always shows all options from 0-100)
    scale_x_continuous(breaks=xbreaks, expand = c(0,0.15))+
    scale_y_continuous(breaks=ybreaks, expand = c(0,0.15))+
    theme(aspect.ratio=aspectrat, 
          panel.background = element_blank(),
          panel.grid = element_blank(), 
          text = element_text(size=plotfontsize),
          panel.border = element_rect(size = 1, fill = NA),
          strip.background = element_blank(),
          strip.text = element_text(hjust = 0, size=plotfontsize),
          legend.title=element_text(size=plotfontsize),
          axis.text = element_text(size=plotfontsize, colour="black"),
          axis.ticks = element_line(colour="black"),
          legend.text = element_text(size=plotfontsize, colour="black"),
          legend.justification = "top")
}
#Plot function for direction data split by trend shape
DirectionPlotEDF <- function(Direction){
  Direction$Prop <- round_any(Direction$Prop*100, 10, floor) #Convert proportions to percentages and round down to nearest 10
  Direction$Prop <- paste0(Direction$Prop, "-", Direction$Prop+10) #Create label (e.g. "0-10" rather than just "0")
  Direction$Prop <- factor(Direction$Prop, levels=sapply(seq(0,90,10), function(x) paste0(x, "-", x+10))) #Convert to factor
   
  if(SamplingType=="Consecutive"){  #Create labels (adjusts based on consecutive or interval sampling)
    xlabel <- "Complete trend length (years)"
    ylabel <- "Sample trend length (years)"
    xbreaks <- (c(seq(4,MCL,6)))
    ybreaks <- (c(seq(3,MCL,6)))
    aspectrat <- 1
    ex <- Direction$CompleteLength
    why <- Direction$NumYears
  } else {
    xlabel <- "Number of years sampled"
    ylabel <- "Samples taken every x years"
    xbreaks <- (c(seq(3,29,6)))
    ybreaks <- (c(seq(0,14,3)))
    aspectrat <- 0.52
    ex <- Direction$NumYears
    why <- Direction$IntervalLength
  }
  
  Direction[Direction$Category=="a) Matching",]$Category <- "Matching" #This block adjusts all the names so the plot works
  Direction[Direction$Category=="c) Erroneous +/-",]$Category <- "Erroneous +/-"
  Direction[Direction$Category=="d) Missed +/-",]$Category <- "Missed +/-"
  Direction[Direction$Category=="b) Opposing",]$Category <- "Opposing"
  Direction$EDF <- as.character(as.numeric(Direction$EDF))
  Direction[Direction$EDF==3,]$EDF <- "a) 1-3"
  Direction[Direction$EDF==5,]$EDF <- "b) 3-5"
  Direction[Direction$EDF==7,]$EDF <- "c) 5-7"
  Direction[Direction$EDF==9,]$EDF <- "d) 7-9"
  Direction$EDF <- factor(Direction$EDF)
  Direction$EDF <- factor(Direction$EDF,levels(Direction$EDF)[c(1,2,3,4)])
  Direction$Category <- factor(Direction$Category)
  Direction$Category <- factor(Direction$Category,levels(Direction$Category)[c(2,4,1,3)])
  
  ggplot(Direction, aes(x=ex,y=why))+
    geom_tile(aes(fill=Prop), colour = "grey40",size = 0.01)+
    facet_grid(Category~EDF)+ #Make the facets show category vs. trend shape
    ylab(ylabel)+
    xlab(xlabel)+
    scale_fill_manual(values = viridis(n=10), name=paste0("Percentage"), guide=guide_legend(reverse = TRUE, keywidth=unit(3, "mm"), keyheight = unit(3, "mm")), drop=FALSE)+ #This sets the legend as the same regardless of which colours appear (it always shows all options from 0-100)
    scale_x_continuous(breaks=xbreaks, expand = c(0,0.15))+
    scale_y_continuous(breaks=ybreaks, expand = c(0,0.15))+
    theme(aspect.ratio=aspectrat, 
          panel.background = element_blank(),
          panel.grid = element_blank(), 
          text = element_text(size=plotfontsize),
          panel.border = element_rect(size = 1, fill = NA),
          strip.background = element_blank(),
          strip.text = element_text(hjust = 0, size=plotfontsize),
          legend.title=element_text(size=plotfontsize),
          axis.text = element_text(size=plotfontsize, colour="black"),
          axis.ticks = element_line(colour="black"),
          legend.text = element_text(size=plotfontsize, colour="black"),
          legend.justification = "top")
}

#General plot function for Magnitude data
MagnitudePlot <- function(Direction){
  MagnitudeSub <- c(0.01,0.025, 0.05, 0.1, 0.25, 0.5) #Subset to the tolerances we want to plot (I modelled more just out of interest)
  Direction <- Direction[Direction$Tolerance %in% MagnitudeSub,] #Subset to the tolerances we want to plot (I modelled more just out of interest)
  Direction$Tolerance <- paste0("± ", Direction$Tolerance) #Add a plusminus sign so labels are correct
  if(SamplingType=="Consecutive"){
    Direction$CompleteLength <- as.character(as.numeric(Direction$CompleteLength))
  }
  if(SamplingType=="Interval"){
    Direction$CompleteLength <- Direction$IntervalLength #Cheeky relabel of interval length to be complete length so that this function can be applied to both data (bit janky but it works, and labels ensure it's labelled as interval length in the actual plot)
  }
  Direction$Tolerance <- as.factor(Direction$Tolerance)
  if(SamplingType=="Consecutive"){ #Add labels to the years (consecutive)
    Direction$CompleteLength <- sapply(Direction$CompleteLength, function(x){
      if(x==5){
        paste0("a) ", x, " years")
      } else if(x==10){
        paste0("b) ", x, " years")
      } else if(x==20){
        paste0("c) ", x, " years")
      } else if(x==30){
        paste0("d) ", x, " years")
      }
    })
    Direction$CompleteLength <- factor(Direction$CompleteLength, levels=c("a) 5 years","b) 10 years","c) 20 years","d) 30 years"))
  } else {
    Direction$CompleteLength <- sapply(Direction$CompleteLength, function(x){ #Add labels to the years between samples (intervals)
      if(x==1){
        paste0("a) Samples taken every ", x, " year")
      } else if(x==3){
        paste0("b) Samples taken every ", x, " years")
      } else if(x==5){
        paste0("c) Samples taken every ", x, " years")
      } else if(x==7){
        paste0("d) Samples taken every ", x, " years")
      } else if(x==9){
        paste0("e) Samples taken every ", x, " years")
      } else if(x==11){
        paste0("f) Samples taken every ", x, " years")
      }
    })
    Direction$CompleteLength <- factor(Direction$CompleteLength, levels=c("a) Samples taken every 1 year","b) Samples taken every 3 years","c) Samples taken every 5 years","d) Samples taken every 7 years","e) Samples taken every 9 years","f) Samples taken every 11 years"))
  }
  if(SamplingType=="Consecutive"){ #Set breaks for plot
    breaksx <- seq(0,MCL,5)
    breaksy <- seq(0,100,10)
  } else {
    breaksx <- seq(3,15,2)
    breaksy <- seq(0,100,10) 
  }
  Direction$PropCorrect <- Direction$PropCorrect*100 #Change proportion correct to a percentage
  ggplot(Direction, aes(x=NumYears,y=PropCorrect, colour=Tolerance))+ #make a line plot!
    geom_line()+
    geom_point(shape=18, size=2)+
    facet_wrap(~CompleteLength, nrow=2)+
    ylab("Percentage within tolerance")+
    xlab("Number of years sampled")+
    scale_colour_manual(values = viridis(n=7), name="Tolerance", guide=guide_legend(reverse = TRUE, keywidth=unit(3, "mm"), keyheight = unit(3, "mm")), drop=FALSE)+
    scale_x_continuous(breaks=breaksx)+ #, expand = c(0,0)
    scale_y_continuous(breaks=breaksy)+ #, expand = c(0,0)
    theme(aspect.ratio=1, 
          panel.background = element_blank(),
          panel.grid = element_blank(), 
          text = element_text(size=plotfontsize),
          panel.border = element_rect(size = 1, fill = NA),
          strip.background = element_blank(),
          strip.text = element_text(hjust = 0),
          legend.title=element_text(size=plotfontsize),
          legend.key=element_blank(),
          legend.justification = "top")
}
#Plot function for magnitude data split by generation length
MagnitudePlotGen <- function(Direction){
  MagnitudeSub <- c(0.01,0.05, 0.1) #Subset to the tolerances we want to plot 
  Direction <- Direction[Direction$Tolerance %in% MagnitudeSub,] #Subset to the tolerances we want to plot
  Direction$Tolerance <- paste0("± ", Direction$Tolerance) #Add plus minus so the labels are correct
  Direction$GenLength <- as.character(as.numeric(Direction$GenLength))
  Direction[Direction$GenLength==5,]$GenLength <- "a) Short" #Get the labels correct
  Direction[Direction$GenLength==10,]$GenLength <- "b) Medium"
  Direction[Direction$GenLength==15,]$GenLength <- "c) Long"
  Direction$GenLength <- factor(Direction$GenLength)
  Direction$GenLength <- factor(Direction$GenLength,levels(Direction$GenLength)[c(1,2,3)])
  if(SamplingType=="Consecutive"){
    Direction$CompleteLength <- as.character(as.numeric(Direction$CompleteLength)) 
  }
  if(SamplingType=="Interval"){
    Direction$CompleteLength <- Direction$IntervalLength #Cheeky relabel of interval length to be complete length so that this function can be applied to both data (bit janky but it works, and labels ensure it's labelled as interval length in the actual plot)
  }
  Direction$Tolerance <- as.factor(Direction$Tolerance)
  if(SamplingType=="Consecutive"){ #Add labels
    Direction$CompleteLength <- paste0(Direction$CompleteLength, " Years")
    Direction$CompleteLength <- factor(Direction$CompleteLength, levels=c("10 Years","20 Years","30 Years"))
  } else {
    Direction$CompleteLength <- sapply(Direction$CompleteLength, function(x){
      if(x==1){
        paste0("Smpls every ", x, "yr")
      } else {
        paste0("Smpls every ", x, "yrs")
      }
    })
  }
  breaksy <- seq(0,100,20)
  breaksx <- seq(0,MCL,5)
  Direction$PropCorrect <- Direction$PropCorrect*100 #Change proportion to percentage
  ggplot(Direction, aes(x=NumYears,y=PropCorrect, colour=GenLength))+ #Make a line plot!
    geom_line()+
    geom_point(shape=18, size=1.5)+
    facet_grid(Tolerance~CompleteLength)+
    ylab("Percentage within tolerance")+
    xlab("Number of years sampled")+
    scale_colour_manual(values = viridis(n=4), name=str_wrap("Generation Length", width=10), guide=guide_legend(reverse = FALSE, keywidth=unit(3, "mm"), keyheight = unit(3, "mm")), drop=FALSE)+
    scale_x_continuous(breaks=breaksx)+ #, expand = c(0,0)
    scale_y_continuous(breaks=breaksy)+ #, expand = c(0,0)
    theme(aspect.ratio=1, 
          panel.background = element_blank(),
          panel.grid = element_blank(), 
          text = element_text(size=plotfontsize),
          panel.border = element_rect(size = 1, fill = NA),
          strip.background = element_blank(),
          strip.text = element_text(hjust = 0),
          legend.title=element_text(size=plotfontsize),
          legend.key=element_blank(),
          legend.justification = "top")
}
#Plot function for magnitude data split by trend shape
MagnitudePlotEDF <- function(Direction){
  MagnitudeSub <- c(0.01,0.05, 0.1) #Subset to the tolerances we want to plot (I modelled more just out of interest)
  Direction <- Direction[Direction$Tolerance %in% MagnitudeSub,] #Subset to the tolerances we want to plot (I modelled more just out of interest)
  Direction$Tolerance <- paste0("± ", Direction$Tolerance) #Add a plusminus sign so labels are correct
  Direction$EDF <- as.character(as.numeric(Direction$EDF))
  Direction[Direction$EDF==3,]$EDF <- "1-3"
  Direction[Direction$EDF==5,]$EDF <- "3-5"
  Direction[Direction$EDF==7,]$EDF <- "5-7"
  Direction[Direction$EDF==9,]$EDF <- "7-9"
  Direction$EDF <- factor(Direction$EDF)
  Direction$EDF <- factor(Direction$EDF,levels(Direction$EDF)[c(1,2,3,4)])
  if(SamplingType=="Consecutive"){
    Direction$CompleteLength <- as.character(as.numeric(Direction$CompleteLength))
  }
  if(SamplingType=="Interval"){
    Direction$CompleteLength <- Direction$IntervalLength #Cheeky relabel of interval length to be complete length so that this function can be applied to both data (bit janky but it works, and labels ensure it's labelled as interval length in the actual plot)
  }
  Direction$Tolerance <- as.factor(Direction$Tolerance)
  if(SamplingType=="Consecutive"){ #Add labels
    Direction$CompleteLength <- paste0(Direction$CompleteLength, " years")
    Direction$CompleteLength <- factor(Direction$CompleteLength, levels=c("10 years","20 years","30 years"))
  } else {
    Direction$CompleteLength <- sapply(Direction$CompleteLength, function(x){
      if(x==1){
        paste0("Smpls every ", x, "yr")
      } else {
        paste0("Smpls every ", x, "yrs")
      }
    })
  }
  breaksy <- seq(0,100,20)
  breaksx <- seq(0,MCL,5)
  Direction$PropCorrect <- Direction$PropCorrect*100 #Change proportion to percentage
  ggplot(Direction, aes(x=NumYears,y=PropCorrect, colour=EDF))+ #Make a line plot!
    geom_line()+
    geom_point(shape=18, size=1.5)+
    facet_grid(Tolerance~CompleteLength)+
    ylab("Percentage within tolerance")+
    xlab("Number of years sampled")+
    scale_colour_manual(values = viridis(n=5), name=str_wrap("EDF", width=10), guide=guide_legend(reverse = FALSE, keywidth=unit(3, "mm"), keyheight = unit(3, "mm")), drop=FALSE)+
    scale_x_continuous(breaks=breaksx)+ #, expand = c(0,0)
    scale_y_continuous(breaks=breaksy)+ #, expand = c(0,0)
    theme(aspect.ratio=1, 
          panel.background = element_blank(),
          panel.grid = element_blank(), 
          text = element_text(size=plotfontsize),
          panel.border = element_rect(size = 1, fill = NA),
          strip.background = element_blank(),
          strip.text = element_text(hjust = 0),
          legend.title=element_text(size=plotfontsize),
          legend.key=element_blank(),
          legend.justification = "top")
}

modelfam <- "nb" #Adjust this if you used a different model fam in above runs

DirectionSummary <- rbindlist(lapply(list.files(path=paste0(ResultsFP, "GLM/Summaries/"), pattern=paste0("^Direction_Consecutive_",modelfam, "..*.csv$"), full.names=TRUE), fread), fill=TRUE) #Read in summarised CONSECUTIVE data from model runs
MagnitudeSummary <- rbindlist(lapply(list.files(path=paste0(ResultsFP, "GLM/Summaries/"), pattern=paste0("^Magnitude_Consecutive_",modelfam, "..*.csv$"), full.names=TRUE), fread), fill=TRUE) #Read in summarised CONSECUTIVE data from model runs

DirectionSummaryInterval <- read.csv(file=paste0(ResultsFP, "GLM/Summaries/Direction_Interval_", modelfam, ".csv")) #Read in summarised data from model runs
DirectionSummaryComplete <- subset(DirectionSummary, CompleteLength==MCL) #Extract MCL data from consecutive data (to add to intervals)
DirectionSummaryComplete$IntervalLength <- 1 #Set the interval length at one
DirectionSummaryComplete$CompleteLength <- NULL #Remove complete length (they're all MCL)
DirectionSummaryInterval <- rbind(DirectionSummaryComplete, DirectionSummaryInterval) #Add MCL data to interval data

MagnitudeSummaryInterval <- read.csv(file=paste0(ResultsFP, "GLM/Summaries/Magnitude_Interval_", modelfam, ".csv")) #As above but for magnitude
MagnitudeSummaryComplete <- subset(MagnitudeSummary, CompleteLength==MCL) 
MagnitudeSummaryComplete$IntervalLength <- 1
MagnitudeSummaryInterval <- rbind(MagnitudeSummaryComplete, MagnitudeSummaryInterval)

##Build and save direction plots
for(SamplingType in c("Consecutive", "Interval")){ #Loop through consecutive and interval
  for(Casttype in c("All", "St")){ #Loop through all data ("All") and standardised data ("St", which checks for biases)
    if(SamplingType=="Consecutive"){ #Subset to relevant category and type
      Direction <- subset(DirectionSummary, CastType==Casttype)
    } else {
      Direction <- subset(DirectionSummaryInterval, CastType==Casttype)
    }
    
    #All
    DirectionAll <- subset(Direction, GenLength==0 & EDF==0) #Remove data subsetted by Generation Length or Trend Shape
    DirectionPlots <- rbindlist(pblapply(c("Matching","Opposing", "MPos","MNeg", "ErrPos", "ErrNeg"), function(Category) PropOverall(DirectionAll, Category))) #Apply the PropOverall Function
    DirectionPlots$Category <- factor(DirectionPlots$Category, levels = c("a) Matching", "c) Erroneous Positive", "e) Missed Positive", "b) Opposing", "d) Erroneous Negative","f) Missed Negative")) #Set levels of factor
    if(SamplingType=="Consecutive"){ #Set plot dimensions
      W <- 173
      H <- 116
    } else {
      W <- 189
      H <- 80
    }
    #Apply plot functions and save
    ggsave(paste0(FiguresFP,"Direction/Direction_", SamplingType, "_", Casttype, "_", modelfam, ".pdf"), DirectionPlot(DirectionPlots), device="pdf", width = W, height = H, units = "mm") #Save as a pdf
  }
  
  #For Generation Length and Trend Shape (i.e. EDF), just use "All" data
  if(SamplingType=="Consecutive"){
    Direction <- subset(DirectionSummary, CastType=="All")
  } else {
    Direction <- subset(DirectionSummaryInterval, CastType=="All")
  }
  
  #Generation length
  DirectionGen <- subset(Direction, GenLength!=0 & EDF==0) #Subset to Generation Length categories (but not EDF)
  DirectionPlotsGen <- rbindlist(lapply(c(5,10,15), function(Gen){ #Apply PropOverallCombine within each generation length
    gendata <- subset(DirectionGen, GenLength==Gen)
    PropGen <- rbindlist(lapply(c("Matching","Opposing", "Missed","Erroneous"), function(Category) PropOverallCombine(gendata, Category)))
    PropGen$GenLength <- Gen
    PropGen$Prop <- as.numeric(PropGen$Prop)
    return(PropGen)
  }))
  if(SamplingType=="Consecutive"){ #Set plot dimensions
    W <- 130
    H <- 130
  } else {
    W <- 190
    H <- 120
  }
  #Apply plot functions and save
  ggsave(paste0(FiguresFP,"Gen/DirectionGen_", SamplingType, "_", modelfam, ".pdf"), DirectionPlotGen(DirectionPlotsGen), device="pdf", width = W, height = H, units = "mm") #Save as an image
  
  #EDF
  DirectionEDF <- subset(Direction, GenLength==0 & EDF!=0) #Subset to EDF categories (but not generation length)
  DirectionPlotsEDF <- rbindlist(lapply(unique(DirectionEDF$EDF), function(edf){ #Apply PropOverallCombine within each EDF category
    edfdata <- subset(DirectionEDF, EDF==edf)
    PropEDF <- rbindlist(lapply(c("Matching","Opposing", "Missed","Erroneous"), function(Category) PropOverallCombine(edfdata, Category)))
    PropEDF$EDF <- edf
    PropEDF$Prop <- as.numeric(PropEDF$Prop)
    return(PropEDF)
  }))
  if(SamplingType=="Consecutive"){ #Set plot dimensions
    W <- 150
    H <- 120
  } else {
    W <- 248
    H <- 132
  }
  #Apply plot functions and save
  ggsave(paste0(FiguresFP,"EDF/Direction_EDF_", SamplingType, "_", modelfam, ".pdf"), DirectionPlotEDF(DirectionPlotsEDF), device="pdf", width = W, height = H, units = "mm") #Save as an image
}

##Build and save magnitude plots
for(SamplingType in c("Consecutive", "Interval")){ #Loop through consecutive and interval
  for(Casttype in c("All", "St")){ #Loop through all data ("All") and standardised data ("St", which checks for biases)
    if(SamplingType=="Consecutive"){ #Subset to relevant category and type
      Magnitude <- subset(MagnitudeSummary, CastType==Casttype)
    } else {
      Magnitude <- subset(MagnitudeSummaryInterval, CastType==Casttype)
    }
    MagnitudeAll <- subset(Magnitude, GenLength==0 & EDF==0) #First look at all data within subsetting
    if(Casttype=="All"){MagnitudeGen <- subset(Magnitude, GenLength!=0 & EDF==0)} #Then at generation length
    if(Casttype=="All"){MagnitudeEDF <- subset(Magnitude, GenLength==0 & EDF!=0)} #Then at trend shape

    if(SamplingType=="Consecutive"){
      MagnitudeAll <- MagnitudeAll[MagnitudeAll$CompleteLength %in% c(5,10,20,30),] #For consecutive, reduce to just a few complete lengths (otherwise too much data to plot)
      if(Casttype=="All"){MagnitudeGen <- MagnitudeGen[MagnitudeGen$CompleteLength %in% c(10,20,30),]}
      if(Casttype=="All"){MagnitudeEDF <- MagnitudeEDF[MagnitudeEDF$CompleteLength %in% c(10,20,30),]}
    } else {
      MagnitudeAll <- MagnitudeAll[MagnitudeAll$IntervalLength %in% c(1,3,5,7,9,11),] #For interval, reduce to just a few interval lengths (otherwise too much data to plot)
      MagnitudeAll <- subset(MagnitudeAll, NumYears<16)
      if(Casttype=="All"){MagnitudeGen <- MagnitudeGen[MagnitudeGen$IntervalLength %in% c(1,3,5,7),]}
      if(Casttype=="All"){MagnitudeEDF <- MagnitudeEDF[MagnitudeEDF$IntervalLength %in% c(1,3,5,7),]}
    }
    
    MagnitudeAll$PropCorrect <- MagnitudeAll$Correct/(MagnitudeAll$Correct+MagnitudeAll$Incorrect) #Find proportion correct
    if(Casttype=="All"){MagnitudeGen$PropCorrect <- MagnitudeGen$Correct/(MagnitudeGen$Correct+MagnitudeGen$Incorrect)}
    if(Casttype=="All"){MagnitudeEDF$PropCorrect <- MagnitudeEDF$Correct/(MagnitudeEDF$Correct+MagnitudeEDF$Incorrect)}

    if(SamplingType=="Consecutive"){ #Set plot dimensions
      H <- 120
      W <- 135
      H_GenEDF <- 120
      W_GenEDF <- 150
    } else {
      H <- 119
      W <- 179
      H_GenEDF <- 100
      W_GenEDF <- 150
    }
    
    #Apply plot functions and save
    ggsave(paste0(FiguresFP,"Magnitude/Magnitude_", SamplingType, "_", Casttype, "_", modelfam, ".pdf"), MagnitudePlot(MagnitudeAll), device="pdf", width = W, height = H, units = "mm") #Save as image
    if(Casttype=="All"){ggsave(paste0(FiguresFP,"Gen/MagnitudeGen_", SamplingType, "_", modelfam, ".pdf"), MagnitudePlotGen(MagnitudeGen), device="pdf", width = W_GenEDF, height = H_GenEDF, units = "mm")}
    if(Casttype=="All"){ggsave(paste0(FiguresFP,"EDF/MagnitudeEDF_", SamplingType, "_", modelfam, ".pdf"), MagnitudePlotEDF(MagnitudeEDF), device="pdf", width = W_GenEDF, height = H_GenEDF, units = "mm")}
  }
}

#### Paper stats ####
modelfam <- "nb" #Set all stats
SamplingType<- "Consecutive"
Casttype <- "All"
plotfontsize <- 10 #Set the font size for plots

#Uncleaned datastats
length(unique(ThirYearBirds$SiteCode))
length(unique(ThirYearBirds$Species))
nrow(unique(ThirYearBirds[,c("SiteCode", "Species")]))

#Cleaned final data stats
TYB_listStats <- rbindlist(TYB_list) #Convert the list of populations into one big dataset
TYB_listStats$Site <- str_split_fixed(TYB_listStats$SiteSpec, "[_]",2)[,1] #Extra site names
TYB_listStats$Spec <- str_split_fixed(TYB_listStats$SiteSpec, "[_]",3)[,2] #Extract species names

length(unique(TYB_listStats$Site)) #Find number of sites
length(unique(TYB_listStats$Spec)) #Find number of species
length(TYB_list)

SpeciesNames <- unique(str_split_fixed(names(TYB_list), "[_]", 3)[,2])
SpecStats <- read.csv(file=paste0(DataFP, "/SpeciesStats.csv"))
SpecStats$Order <- as.factor(SpecStats$Order)
SpecStats$Order <- recode_factor(SpecStats$Order, "CHARADRIIFORMES" = "Charadriiformes", "ANSERIFORMES" = "Anseriformes", "GRUIFORMES" = "Gruiformes", "CICONIIFORMES" = "Ciconiiformes",
                                 "SULIFORMES" = "Suliformes", "PELECANIFORMES" = "Pelicaniformes", "PROCELLARIIFORMES" = "Procellariiformes", "GAVIIFORMES" = "Gaviformes", 
                                 "PHOENICOPTERIFORMES" = "Phoenicopteriformes", "PODICIPEDIFORMES" = "Podicipediformes")
SpecStats$Genus <- str_split_fixed(SpecStats$Species, "[ ]", 2)[,1]
SpecStats2 <- SpecStats[SpecStats$Species %in% SpeciesNames,][,c("Order", "Family", "Species")]
write.csv(SpecStats2, paste0(FiguresFP,"SpeciesData.csv"), row.names = FALSE)

#Standardised data stat
length(TYB_St) #Number of standardised populations
length(unique(str_split_fixed(TYB_St, "[_]", 3)[,2])) #Number of standardised species

#Check for overdispersion
OverDis <- sapply(TYB_list, function(x) var(x$Count)/mean(x$Count))
length(OverDis[(OverDis>1)])/length(OverDis) #Proportion of data that is over dispersed
length(OverDis[(OverDis>10)])/length(OverDis) #Proportion of data that is over dispersed by over an order of magnitude

#Modelled trends stats
AllModels <- read.csv(paste0(ResultsFP, "GLM/Consecutive/Consecutive_GLM_",MCL, "Years_1StartYear_nb.csv")) #Read in just MCL year data
AllModels <- AllModels[!is.na(AllModels$Slope),] #Remove NAs

AllModelsSub <- subset(AllModels, Slope<1 & Slope>-1) #Find out how many trends are between -1 and 1
nrow(AllModelsSub)/nrow(AllModels)

nrow(subset(AllModels, EDF<1.0001))/nrow(AllModels) #Find out the proportion of trends that have an EDF of 1 (i.e. are linear)
nrow(subset(AllModels, Significant<=0.05))/nrow(AllModels) #Find out the proportion of trends that are significant
mean(AllModels$Slope) #Find the mean of all trends
max(AllModels$Slope) #Find the max trend
min(AllModels$Slope) #Find the min trend

GrowthRateHist <- ggplot(data=AllModelsSub)+ #Make a histogram of all trend growth rate values
  geom_freqpoly(aes(x=Slope), binwidth=0.01)+
  scale_x_continuous(breaks=c(seq(-1,1,0.5)), expand = c(0,0))+
  scale_y_continuous(limits=c(0, 2000), expand = c(0,0))+
  ggtitle("a)")+
  xlab("Population Growth Rate (r)")+
  ylab("Frequency")+
  theme(aspect.ratio=1, 
        panel.background = element_blank(),
        panel.grid = element_blank(), 
        text = element_text(size=plotfontsize),
        panel.border = element_rect(size = 1, fill = NA),
        plot.title = element_text(size=plotfontsize))

EDFHist <- ggplot(data=AllModels)+ #Make a histogram of all EDFs
  geom_freqpoly(aes(x=EDF), binwidth=0.5)+
  scale_x_continuous(breaks=c(0,2,4,6,8,10),expand = c(0,0))+
  scale_y_continuous(limits=c(0, 6000), expand = c(0,0))+
  xlab("Estimated Degrees of Freedom")+
  ylab("Frequency")+
  ggtitle("b)")+
  theme(aspect.ratio=1, 
        panel.background = element_blank(),
        panel.grid = element_blank(), 
        text = element_text(size=plotfontsize),
        panel.border = element_rect(size = 1, fill = NA),
        plot.title = element_text(size=plotfontsize))

W <- 200 #Set plot dimensions
H <- 110

ggsave(paste0(FiguresFP,"SummaryStats.pdf"), grid.arrange(GrowthRateHist,EDFHist, ncol=2), device="pdf", width = W, height = H, units = "mm") #Save as an image

#### Build supporting data ####
modelfam <- "nb"

#This function gives us the proportion of corrects for each CompleteLength (as in the plots section)
#(The models sections return the raw number of samples in each category, but not the proportion, so we need to calculate)

PropOverallSuppData <- function(Direction, Category){
  Direction2 <- if(SamplingType=="Consecutive"){split(Direction, Direction$CompleteLength)}else{split(Direction, Direction$NumYears)}
  DataFrame <- rbindlist(lapply(Direction2, function(y){
    okcast <- if(SamplingType=="Consecutive"){
      dcast(y, NumYears~variable, value.var="value")
    } else {
      dcast(y, IntervalLength~variable, value.var="value")
    }
    if(Category=="Matching"){
      okcast$Prop <- sapply(1:nrow(okcast), function(x) okcast$Matching[x]/sum(okcast[x,2:ncol(okcast)]))
      okcast$Category <- "Matching"
    } else if(Category=="EPos"){
      okcast$Prop <- sapply(1:nrow(okcast), function(x) okcast$'Erroneous Positive'[x]/sum(okcast[x,2:ncol(okcast)]))
      okcast$Category <- "Erroneous Negative"
    } else if(Category=="ENeg"){
      okcast$Prop <- sapply(1:nrow(okcast), function(x) okcast$'Erroneous Negative'[x]/sum(okcast[x,2:ncol(okcast)]))
      okcast$Category <- "Erroneous Positive"
    } else if(Category=="MPos"){
      okcast$Prop <- sapply(1:nrow(okcast), function(x) okcast$'Missed Positive'[x]/sum(okcast[x,2:ncol(okcast)]))
      okcast$Category <- "Missed Negative"
    } else if(Category=="MNeg"){
      okcast$Prop <- sapply(1:nrow(okcast), function(x) okcast$'Missed Negative'[x]/sum(okcast[x,2:ncol(okcast)]))
      okcast$Category <- "Missed Positive"
    } else {
      okcast$Prop <- sapply(1:nrow(okcast), function(x) okcast$Opposing[x]/sum(okcast[x,2:ncol(okcast)]))
      okcast$Category <- "Opposing"
    }
    if(SamplingType=="Consecutive"){
      okcast <- okcast[,c("NumYears", "Prop", "Category")]
      okcast$CompleteLength <- unique(y$CompleteLength)
      okcast$NumYears <- as.numeric(as.character(okcast$NumYears))
    } else {
      okcast <- okcast[,c("IntervalLength", "Prop", "Category")]
      okcast$NumYears <- unique(y$NumYears)
      okcast$IntervalLength <- as.numeric(as.character(okcast$IntervalLength))
    }
    okcast$Prop <- as.numeric(okcast$Prop)
    okcast[is.na(okcast)] <- 0
    return(okcast)
  }))
  return(DataFrame)
} 

DirectionSummary <- rbindlist(lapply(list.files(path=paste0(ResultsFP, "GLM/Summaries/"), pattern=paste0("^Direction_Consecutive_",modelfam, "..*.csv$"), full.names=TRUE), fread), fill=TRUE) #Read in data
MagnitudeSummary <- rbindlist(lapply(list.files(path=paste0(ResultsFP, "GLM/Summaries/"), pattern=paste0("^Magnitude_Consecutive_",modelfam, "..*.csv$"), full.names=TRUE), fread), fill=TRUE) #Read in data

DirectionSummaryInterval <- read.csv(file=paste0(ResultsFP, "GLM/Summaries/Direction_Interval_", modelfam, ".csv")) #As in plots section, add MCL data to intervals
DirectionSummaryComplete <- subset(DirectionSummary, CompleteLength==MCL)
DirectionSummaryComplete$IntervalLength <- 1
DirectionSummaryComplete$CompleteLength <- NULL
DirectionSummaryInterval <- rbind(DirectionSummaryComplete, DirectionSummaryInterval)

MagnitudeSummaryInterval <- read.csv(file=paste0(ResultsFP, "GLM/Summaries/Magnitude_Interval_", modelfam, ".csv")) #As in plots section, add MCL data to intervals
MagnitudeSummaryComplete <- subset(MagnitudeSummary, CompleteLength==MCL)
MagnitudeSummaryComplete$IntervalLength <- 1
MagnitudeSummaryInterval <- rbind(MagnitudeSummaryComplete, MagnitudeSummaryInterval)

for (SamplingType in c("Consecutive", "Interval")){ # Loop through consecutive and interval
  if(SamplingType=="Consecutive"){
    Direction <- subset(DirectionSummary, CastType=="All") #Get relevant data
  } else {
    Direction <- subset(DirectionSummaryInterval, CastType=="All")
  }
  
  DirectionAll <- subset(Direction, GenLength==0 & EDF==0) #First get proportions from all data
  DirectionPlots <- rbindlist(pblapply(c("Matching","Opposing", "MPos","MNeg", "EPos", "ENeg"), function(Category) PropOverallSuppData(DirectionAll, Category))) #Apply PropOverallSuppData function
  
  DirectionGen <- subset(Direction, GenLength!=0 & EDF==0) #Repeat for Generation length data
  DirectionPlotsGen <- rbindlist(lapply(c(5,10,15), function(Gen){
    gendata <- subset(DirectionGen, GenLength==Gen)
    PropGen <- rbindlist(lapply(c("Matching","Opposing", "MPos","MNeg", "EPos", "ENeg"), function(Category) PropOverallSuppData(gendata, Category)))
    PropGen$GenLength <- Gen
    PropGen$Prop <- as.numeric(PropGen$Prop)
    return(PropGen)
  }))
  
  DirectionEDF <- subset(Direction, GenLength==0 & EDF!=0) #Repeat for trend shape data
  DirectionPlotsEDF <- rbindlist(lapply(unique(DirectionEDF$EDF), function(edf){
    edfdata <- subset(DirectionEDF, EDF==edf)
    PropEDF <- rbindlist(lapply(c("Matching","Opposing", "MPos","MNeg", "EPos", "ENeg"), function(Category) PropOverallSuppData(edfdata, Category)))
    PropEDF$EDF <- edf
    PropEDF$Prop <- as.numeric(PropEDF$Prop)
    return(PropEDF)
  }))
  
  DirectionPlotsGen$EDF <- NA
  DirectionPlotsEDF$GenLength <- NA
  DirectionPlots[,c("EDF", "GenLength")] <- NA
  
  DirectionData <- rbind(DirectionPlots, DirectionPlotsEDF, DirectionPlotsGen) #bring data together
  if(SamplingType=="Consecutive"){ #Rename columns for clarity
    names(DirectionData)[1] <- "SampleLength"
  } else {
    names(MagnitudeData)[1] <- "SampleLength"
    names(DirectionData)[4] <- "NumYearsInSample"
    DirectionData$CompleteLength <- MCL
  }
  
  write.csv(DirectionData, paste0(FiguresFP, "Supporting/DirectionData",SamplingType, ".csv"), row.names=FALSE) #Save!
  
  if(SamplingType=="Consecutive"){
    Magnitude <- subset(MagnitudeSummary, CastType=="All")
  } else {
    Magnitude <- subset(MagnitudeSummaryInterval, CastType=="All")
  }
  MagnitudeAll <- subset(Magnitude, GenLength==0 & EDF==0) #Subset to various data categories
  MagnitudeGen <- subset(Magnitude, GenLength!=0 & EDF==0)
  MagnitudeEDF <- subset(Magnitude, GenLength==0 & EDF!=0)
  
  MagnitudeAll$PropCorrect <- MagnitudeAll$Correct/(MagnitudeAll$Correct+MagnitudeAll$Incorrect) #Find proportion correct
  MagnitudeGen$PropCorrect <- MagnitudeGen$Correct/(MagnitudeGen$Correct+MagnitudeGen$Incorrect)
  MagnitudeEDF$PropCorrect <- MagnitudeEDF$Correct/(MagnitudeEDF$Correct+MagnitudeEDF$Incorrect)
  
  MagnitudeAll[,c("Correct", "Incorrect","SamplingType", "Model", "CastType")] <- NULL #Remove irrelevant columns
  MagnitudeAll[,c("GenLength", "EDF")] <- NA #Make these NA for "All" data
  MagnitudeGen[,c("Correct", "Incorrect", "SamplingType","Model", "CastType")] <- NULL
  MagnitudeGen[,c("EDF")] <- NA #make EDF NA for GenLength data
  MagnitudeEDF[,c("Correct", "Incorrect", "SamplingType","Model", "CastType")] <- NULL
  MagnitudeEDF[,c("GenLength")] <- NA #make genlength NA for EDF data
  
  MagnitudeData <- rbind(MagnitudeAll, MagnitudeGen, MagnitudeEDF) #Bring the three together
  if(SamplingType=="Consecutive"){ #Rename columns for clarity
    names(MagnitudeData)[1] <- "SampleLength"
  } else {
    names(MagnitudeData)[1] <- "SampleLength"
    names(MagnitudeData)[7] <- "NumYearsInSample"
  }
  names(MagnitudeData)[ncol(MagnitudeData)] <- "PropWithinTolerance"
  write.csv(MagnitudeData, paste0(FiguresFP, "Supporting/MagnitudeData",SamplingType, ".csv"), row.names=FALSE) #Save!
  
}

#### Supplementary analysis for supporting information section 4 ####
#This is a little bit of extra supporting code to calculate how often insignificant samples are still approximating the same trend direction/magnitude as complete samples. (In the paper, we treat any insigicant slopes as just 'insignificant' and don't consider the actual slope)
#So, in the direction section, we are looking for cases where the complete trend was significant, and the sample insignificant, and comparing slope direction. This has been termed "missed sig" in the code below. 
#In the magnitude section we are looking for cases where the complete trend was significant and the sample insignificant, and comparing their slope magnitude to see whether the sample is still within the tolerance range
#This set of code requires that the initial models have been run, but not the summary sections of code

### Write summary functions
DirectionInsig <- function(DATA, Standardise, GenLength, EDFNum, Significance){
  Input <- DATA
  SamplingType <- unique(Input$SamplingType)
  #Reduce the data according to the various subsets
  if(GenLength==0){
    if(Standardise==TRUE){Input <- Input[Input$SiteSpec %in% TYB_St,]}
  } else {
    GenLengthSub <- subset(GenLengthSpec, GenLengthMax==GenLength)
    Input <- Input[Input$SiteSpec %in% GenLengthSub$SiteSpec,]
  } 
  
  if(EDFNum!=0){
    Input <- DATA
    Input[is.na(Input$EDFCompleteLength),]$EDFCompleteLength <- 0
    Input <- subset(Input, EDFCompleteLength<EDFNum)
  }
  
  if(nrow(Input)==0){
    return(NULL)
  }
  
  #Change the slope to just positive, negative or insignificant
  Input$Slope <- sapply(Input$Slope, function (x) if(is.na(x)){"Insignificant"} else if(x > 0){ "Positive" } else{ "Negative" })
  Input$CompleteLengthSlope <- sapply(Input$CompleteLengthSlope, function (x) if(is.na(x)){"Insignificant"} else if(x > 0){ "Positive" } else{ "Negative" })
  
  Input <- subset(Input, Slope != "Insignificant" & CompleteLengthSlope != "Insignificant")
  
  #Create a binary value for significance based on p value
  Input$Significant <- sapply(Input$Significant, function (x) if(is.na(x)){0} else if(x < Significance){1} else{0})
  Input$SignificantCompleteLength <- sapply(Input$SignificantCompleteLength, function (x) if(is.na(x)){0} else if(x < Significance){1} else{0})
  
  Input <- Input[,c("SiteSpec", "NumYears", "Slope", "Significant", "CompleteLengthSlope", "SignificantCompleteLength")]
  
  #Investigate the missed +/-
  MissedSig <- subset(Input, Significant==0 & SignificantCompleteLength==1)
  MissedSig$Slope <- ifelse(MissedSig$Slope==MissedSig$CompleteLengthSlope, "Matching", "Opposing")
  MissedSig <- MissedSig[,c("SiteSpec", "NumYears", "Slope")]

  summarise <- MissedSig
  
  rm(Input)
  
  #Cast to find number of populations of each category
  summarise <- dcast(summarise, NumYears~Slope, length, value.var="Slope")
  summarise$PropCorrect <- summarise$Matching/(summarise$Matching + summarise$Opposing)
  summarise[,c("Matching", "Opposing")] <- NULL
  
  #Split numyears for interval
  if(SamplingType=="Interval"){
    summarise[,c("IntervalLength", "NumYears")] <- str_split_fixed(summarise$NumYears, "[.]", 2)
  }
  
  #Order by number of years, add metadata
  summarise$NumYears <- as.numeric(as.character(summarise$NumYears))
  summarise <- summarise[order(-summarise$NumYears),]
  summarise$SamplingType <- SamplingType
  summarise$Model <- modelfam
  if(SamplingType=="Consecutive"){
    summarise$CompleteLength <- unique(DATA$CompleteLengthNum)
  }
  summarise$GenLength <- GenLength
  if(!Standardise){
    summarise$CastType <- "All"
  } else{
    summarise$CastType <- "St"
  }
  summarise$EDF <- EDFNum
  return(summarise)
} #This is a modified version of the same function in the "Functions" Section
MagnitudeInsig <- function(DATA, Standardise, GenLength, EDFNum, Significance){
  Input <- DATA
  SamplingType <- unique(Input$SamplingType)
  
  #Reduce the data according to the various subsets
  
  if(GenLength==0){
    if(Standardise==TRUE){Input <- Input[Input$SiteSpec %in% TYB_St,]}
  } else {
    GenLengthSub <- subset(GenLengthSpec, GenLengthMax==GenLength)
    Input <- Input[Input$SiteSpec %in% GenLengthSub$SiteSpec,]
  } 
  
  if(EDFNum!=0){
    Input <- DATA
    Input[is.na(Input$EDFCompleteLength),]$EDFCompleteLength <- 0
    Input <- subset(Input, EDFCompleteLength<EDFNum)
  }
  
  
  Input <- subset(Input, Significant>Significance & SignificantCompleteLength<Significance)
  
  if(nrow(Input)==0){
    return(NULL)
  }
  
  if(SamplingType=="Consecutive"){
    Input[,c("SamplingType", "Model", "CompleteLengthNum", "EDF", "EDFCompleteLength")] <- NULL
  } else{
    Input <- Input[,c("SiteSpec","NumYears", "Slope", "CompleteLengthSlope")]
    Input <- Input[!is.na(Input$Slope),]
  }
  
  Input <- Input[complete.cases(Input)] #Remove NAs
  
  #Find correct and incorrect for the various tolerances
  summarisePos <- rbindlist(lapply(c(0.01, 0.025, 0.05, 0.1, 0.25, 0.5), function(Tol){
    ok <- subset(Input, Slope>0 & CompleteLengthSlope>0)
    ok$Trend <- ifelse(ok$Slope<(ok$CompleteLengthSlope+Tol) & ok$Slope>(ok$CompleteLengthSlope-Tol), "Correct", "Incorrect")
    ok$Tolerance <- Tol
    summarise <- dcast(ok, NumYears+Tolerance~Trend, length, value.var="Trend")
    summarise$Investigation <- "Pos"
    return(summarise)
  }), fill=TRUE)
  
  summariseNeg <- rbindlist(lapply(c(0.01, 0.025, 0.05, 0.1, 0.25, 0.5), function(Tol){
    ok <- subset(Input, Slope<0 & CompleteLengthSlope<0)
    ok$Trend <- ifelse(ok$Slope<(ok$CompleteLengthSlope+Tol) & ok$Slope>(ok$CompleteLengthSlope-Tol), "Correct", "Incorrect")
    ok$Tolerance <- Tol
    summarise <- dcast(ok, NumYears+Tolerance~Trend, length, value.var="Trend")
    summarise$Investigation <- "Neg"
    return(summarise)
  }), fill=TRUE)
  
  summarisePosNeg <- rbindlist(lapply(c(0.01, 0.025, 0.05, 0.1, 0.25, 0.5), function(Tol){
    ok <- subset(Input, Slope>0 & CompleteLengthSlope<0)
    ok <- rbind(ok, subset(Input, Slope<0 & CompleteLengthSlope>0))
    ok$Trend <- ifelse(ok$Slope<(ok$CompleteLengthSlope+Tol) & ok$Slope>(ok$CompleteLengthSlope-Tol), "Correct", "Incorrect")
    ok$Tolerance <- Tol
    summarise <- dcast(ok, NumYears+Tolerance~Trend, length, value.var="Trend")
    summarise$Investigation <- "PosNeg"
    return(summarise)
  }), fill=TRUE)
  
  rm(Input)
  
  summarise <- rbind(summarisePos, summariseNeg, summarisePosNeg)
  summarise[is.na(summarise$Correct),]$Correct <- 0
  #Split numyears for interval
  if(SamplingType=="Interval"){
    summarise[,c("IntervalLength")] <- str_split_fixed(summarise$NumYears, "[.]", 2)[,1]
    summarise[,c("NumYears")] <- str_split_fixed(summarise$NumYears, "[.]", 2)[,2]
  }
  
  #Add metadata
  summarise$NumYears <- as.numeric(as.character(summarise$NumYears))
  summarise$SamplingType<- SamplingType
  summarise$Model <- modelfam
  summarise$CompleteLength <- unique(DATA$CompleteLengthNum)
  summarise$GenLength <- GenLength
  if(!Standardise){
    summarise$CastType <- "All"
  } else {
    summarise$CastType <- "St"
  } 
  summarise$EDF <- EDFNum
  return(summarise)
} #This is a modified version of the same function in the "Functions" Section

### Summarise Consecutive
Consecutive <- rbindlist(pblapply(list.files(path=paste0(ResultsFP, "GLM/Consecutive/"), pattern=paste0("*", modelfam, ".csv"), full.names=TRUE), fread), fill=TRUE) 
#The following function organises the data so that samples can be compared to multiple complete trend lengths. E.g. A complete trend 10 years long taken from years 10-19 could have 3 year samples taken from it starting from years 10,11,12...17 etc)
#It then compares sample trend to complete trend according to the two comparison techniques (Direction and Magnitude), but modified for this supplementary section
CastConsecutive <- pblapply(c(4:MCL), function(CompleteLength){
  if((file.exists(paste0(ResultsFP, "GLM/SummariesInsig/Direction_Consecutive_", modelfam, "_", CompleteLength,".csv"))&
      file.exists(paste0(ResultsFP, "GLM/SummariesInsig/Magnitude_Consecutive_", modelfam, "_", CompleteLength,".csv")))==TRUE){
    return(NULL)
  } else {
    write.csv(NULL, paste0(ResultsFP, "GLM/SummariesInsig/Direction_Consecutive_", modelfam, "_", CompleteLength,".csv"))
    write.csv(NULL, paste0(ResultsFP, "GLM/SummariesInsig/Magnitude_Consecutive_", modelfam, "_", CompleteLength,".csv"))
    
    #Work down through the years, removing the start years that don't fit (e.g. start yr at yr21 when we're only going up to 20now)
    CompleteLengthSub <- Consecutive
    CompleteLengthSub[,c("SamplingType", "Model")] <- NULL
    CompleteLengthSlopes <- subset(CompleteLengthSub, NumYears==CompleteLength)
    names(CompleteLengthSlopes) <- c("SiteSpec","CompleteLengthSlope", "SignificantCompleteLength", "EDFCompleteLength", "CompleteLengthNum", "CompleteLengthStartYear")
    
    SamplesSub <- subset(CompleteLengthSub, NumYears<CompleteLength)
    rm(CompleteLengthSub)
    AllTheStartYears <- rbindlist(pblapply(unique(CompleteLengthSlopes$CompleteLengthStartYear), function(x){
      CompleteLengthSubStartYear <- subset(CompleteLengthSlopes, CompleteLengthStartYear==x)
      StartYearSub <- SamplesSub
      StartYearSub <- subset(StartYearSub, StartYear > (x-1))
      StartYearSub$Keep <- ifelse((StartYearSub$StartYear+(StartYearSub$NumYears)-1) < (x+CompleteLength), 1, 0)
      StartYearSub <- subset(StartYearSub, Keep==1)
      StartYearMerge <- merge(StartYearSub, CompleteLengthSubStartYear, All=T)
      return(StartYearMerge)
    }))
    rm(CompleteLengthSlopes, SamplesSub)
    AllTheStartYears$Keep <- NULL
    AllTheStartYears$Model <- modelfam
    AllTheStartYears$SamplingType <- "Consecutive"
    
    Direction_All <- rbindlist(pblapply(c(0,5,10,15), function(x) DirectionInsig(AllTheStartYears, Standardise=FALSE, GenLength=x, EDFNum=0, Significance = 0.05)))
    Direction_St <- rbindlist(lapply(c(0,5,10,15), function(x) DirectionInsig(AllTheStartYears, Standardise=TRUE, GenLength=x, EDFNum=0, Significance = 0.05)))
    Direction_EDF <- rbindlist(pblapply(c(3,5,7,9), function(x) DirectionInsig(AllTheStartYears, Standardise=FALSE, GenLength=0, EDFNum=x, Significance = 0.05)))
    write.csv(rbind(Direction_All, Direction_St, Direction_EDF), file=paste0(ResultsFP, "GLM/SummariesInsig/Direction_Consecutive_", modelfam, "_", CompleteLength,".csv"), row.names=FALSE)
    
    Magnitude_All <- rbindlist(pblapply(c(0,5,10,15), function(x) MagnitudeInsig(AllTheStartYears, Standardise=FALSE, GenLength=x, EDFNum=0, Significance = 0.05)))
    Magnitude_St <- rbindlist(pblapply(c(0,5,10,15), function(x) MagnitudeInsig(AllTheStartYears, Standardise=TRUE, GenLength=x, EDFNum=0, Significance = 0.05)))
    Magnitude_EDF <- rbindlist(pblapply(c(3,5,7,9), function(x) MagnitudeInsig(AllTheStartYears, Standardise=FALSE, GenLength=0, EDFNum=x, Significance = 0.05)))
    write.csv(rbind(Magnitude_All, Magnitude_St, Magnitude_EDF), file=paste0(ResultsFP, "GLM/SummariesInsig/Magnitude_Consecutive_", modelfam, "_", CompleteLength,".csv"), row.names=FALSE)
  }
})

### Summarise Interval
if((file.exists(paste0(ResultsFP, "GLM/SummariesInsig/Direction_Interval_", modelfam, ".csv")) &
    file.exists(paste0(ResultsFP, "GLM/SummariesInsig/Magnitude_Interval_", modelfam, ".csv")))==FALSE){
  write.csv(NULL, paste0(ResultsFP, "GLM/SummariesInsig/Direction_Interval_", modelfam, ".csv"))
  write.csv(NULL, paste0(ResultsFP, "GLM/SummariesInsig/Magnitude_Interval_", modelfam, ".csv"))
  
  Interval <- fread(paste0(ResultsFP, "GLM/Intervals/Intervals_", modelfam, ".csv"))
  CompleteTrend <- fread(paste0(ResultsFP, "GLM/Consecutive/Consecutive_GLM_", MCL, "Years_1StartYear_nb.csv"))[,c("SiteSpec", "Significant", "Slope", "EDF", "NumYears")] #Read in the models run on full 30 years of data (for comparison to complete)
  names(CompleteTrend) <- c("SiteSpec", "SignificantCompleteLength", "CompleteLengthSlope" , "EDFCompleteLength", "CompleteLengthNum")
  Interval <- merge(Interval, CompleteTrend, by="SiteSpec") #Merge complete MCL years with the interval runs
  Interval$NumYears <- paste0(Interval$IntervalLength, ".", Interval$NumYears) #Pull these two together to enable the intervals to run using the functions
  
  Direction_All <- rbindlist(pblapply(c(0,5,10,15), function(x) DirectionInsig(Interval, Standardise=FALSE, GenLength=x, EDFNum=0, Significance = 0.05)))
  Direction_St <- rbindlist(pblapply(c(0,5,10,15), function(x) DirectionInsig(Interval, Standardise=TRUE, GenLength=x, EDFNum=0, Significance = 0.05)))
  Direction_EDF <- rbindlist(pblapply(c(3,5,7,9), function(x) DirectionInsig(Interval, Standardise=FALSE, GenLength=0, EDFNum=x, Significance = 0.05)))
  
  write.csv(rbind(Direction_All, Direction_St, Direction_EDF), file=paste0(ResultsFP, "GLM/SummariesInsig/Direction_Interval_", modelfam, ".csv"), row.names=FALSE)
  
  Magnitude_All <- rbindlist(pblapply(c(0,5,10,15), function(x) MagnitudeInsig(Interval, Standardise=FALSE, GenLength=x, EDFNum=0, Significance = 0.05)))
  Magnitude_St <- rbindlist(pblapply(c(0,5,10,15), function(x) MagnitudeInsig(Interval, Standardise=TRUE, GenLength=x, EDFNum=0, Significance = 0.05)))
  Magnitude_EDF <- rbindlist(pblapply(c(3,5,7,9), function(x) MagnitudeInsig(Interval, Standardise=FALSE, GenLength=0, EDFNum=x, Significance = 0.05)))
  
  write.csv(rbind(Magnitude_All, Magnitude_St, Magnitude_EDF), file=paste0(ResultsFP, "GLM/SummariesInsig/Magnitude_Interval_", modelfam, ".csv"), row.names=FALSE)
}

### Extract and export summary data
modelfam <- "nb" #Adjust this if you used a different model fam in above runs
plotfontsize <- 10 #Set the font size for plots

#Read in summaries
DirectionSummary <- rbindlist(lapply(list.files(path=paste0(ResultsFP, "GLM/SummariesInsig/"), pattern=paste0("^Direction_Consecutive_",modelfam, "..*.csv$"), full.names=TRUE), fread), fill=TRUE) #Read in summarised CONSECUTIVE data from model runs
MagnitudeSummary1 <- rbindlist(lapply(list.files(path=paste0(ResultsFP, "GLM/SummariesInsig/"), pattern=paste0("^Magnitude_Consecutive_",modelfam, "..*.csv$"), full.names=TRUE), fread), fill=TRUE) #Read in summarised CONSECUTIVE data from model runs

DirectionSummaryData <- subset(DirectionSummary, CastType=="All")
DirectionSummaryData[,c("SamplingType", "Model", "CastType")] <- NULL
names(DirectionSummaryData) <- c("SampleLength", "PropCorrect", "CompleteLength", "GenLength", "EDF")
write.csv(DirectionSummaryData, paste0(FiguresFP, "Supporting/DirectionDataConsecutive_SupportingDataSection4.csv"), row.names=FALSE)

#This block organises the direction interval data
DirectionSummaryInterval <- read.csv(file=paste0(ResultsFP, "GLM/SummariesInsig/Direction_Interval_", modelfam, ".csv")) #Read in summarised data from model runs

DirectionSummaryComplete <- subset(DirectionSummary, CompleteLength==MCL) #Extract MCL data from consecutive data (to add to intervals)
DirectionSummaryComplete$IntervalLength <- 1 #Set the interval length at one
DirectionSummaryComplete$CompleteLength <- NULL #Remove complete length (they're all MCL)
DirectionSummaryInterval <- rbind(DirectionSummaryComplete, DirectionSummaryInterval) #Add MCL data to interval data

DirectionSummaryIntervalData <- subset(DirectionSummaryInterval, CastType=="All")
DirectionSummaryIntervalData[,c("SamplingType", "Model", "CastType")] <- NULL
names(DirectionSummaryIntervalData) <- c("SampleLength", "PropCorrect", "IntervalLength", "GenLength", "EDF")
write.csv(DirectionSummaryIntervalData, paste0(FiguresFP, "Supporting/DirectionDataInterval_SupportingDataSection4.csv"), row.names=FALSE) #Write out the data

#This block organises the magnitude data, by calculating the number of corrects and incorrects (within each tolerance level) and getting a proportion
MagnitudeSummary2 <- dcast(MagnitudeSummary1, NumYears + Tolerance + SamplingType + Model + CompleteLength + GenLength + CastType + EDF ~ Investigation, value.var="Correct")
MagnitudeSummary2$Correct <- MagnitudeSummary2$Neg + MagnitudeSummary2$Pos + MagnitudeSummary2$PosNeg
MagnitudeSummary2[,c("Pos", "Neg", "PosNeg")] <- NULL
MagnitudeSummary3 <- dcast(MagnitudeSummary1, NumYears + Tolerance + SamplingType + Model + CompleteLength + GenLength + CastType + EDF ~ Investigation, value.var="Incorrect")
MagnitudeSummary3$Incorrect <- MagnitudeSummary3$Neg + MagnitudeSummary3$Pos + MagnitudeSummary3$PosNeg
MagnitudeSummary3[,c("Pos", "Neg", "PosNeg")] <- NULL
MagnitudeSummary <- merge(MagnitudeSummary2, MagnitudeSummary3)

#This does the same but for the interval magnitude data
MagnitudeSummaryInterval <- read.csv(file=paste0(ResultsFP, "GLM/SummariesInsig/Magnitude_Interval_", modelfam, ".csv")) #As above but for magnitude
MagnitudeSummaryComplete <- subset(MagnitudeSummary1, CompleteLength==MCL) 
MagnitudeSummaryComplete$IntervalLength <- 1
MagnitudeSummaryInterval <- rbind(MagnitudeSummaryComplete, MagnitudeSummaryInterval)

MagnitudeSummaryInterval2 <- dcast(MagnitudeSummaryInterval, NumYears + Tolerance + SamplingType + Model + CompleteLength + GenLength + CastType + EDF + IntervalLength ~ Investigation, value.var="Correct")
MagnitudeSummaryInterval2$Correct <- MagnitudeSummaryInterval2$Neg + MagnitudeSummaryInterval2$Pos + MagnitudeSummaryInterval2$PosNeg
MagnitudeSummaryInterval2[,c("Pos", "Neg", "PosNeg")] <- NULL
MagnitudeSummaryInterval3 <- dcast(MagnitudeSummaryInterval, NumYears + Tolerance + SamplingType + Model + CompleteLength + GenLength + CastType + EDF + IntervalLength ~ Investigation, value.var="Incorrect")
MagnitudeSummaryInterval3$Incorrect <- MagnitudeSummaryInterval3$Neg + MagnitudeSummaryInterval3$Pos + MagnitudeSummaryInterval3$PosNeg
MagnitudeSummaryInterval3[,c("Pos", "Neg", "PosNeg")] <- NULL
MagnitudeSummaryInterval <- merge(MagnitudeSummaryInterval2, MagnitudeSummaryInterval3) #"Tolerance", "SamplingType", "Model", "CompleteLength", "GenLength", "CastType", "EDF", "IntervalLength"

MagnitudeSummary$PropCorrect <- MagnitudeSummary$Correct/(MagnitudeSummary$Correct+MagnitudeSummary$Incorrect) #Find proportion correct
MagnitudeSummaryData <- subset(MagnitudeSummary, CastType=="All")
MagnitudeSummaryData[,c("Correct", "Incorrect", "SamplingType", "Model", "CastType")] <- NULL
names(MagnitudeSummaryData) <- c("SampleLength", "Tolerance", "CompleteLength", "GenLength", "EDF", "PropWithinTolerance")
write.csv(MagnitudeSummaryData, paste0(FiguresFP, "Supporting/MagntiudeDataConsecutive_SupportingDataSection4.csv"), row.names=FALSE) #Write out the data

MagnitudeSummaryInterval$PropCorrect <- MagnitudeSummaryInterval$Correct/(MagnitudeSummaryInterval$Correct+MagnitudeSummaryInterval$Incorrect) #Find proportion correct
MagnitudeSummaryIntervalData <- subset(MagnitudeSummaryInterval, CastType=="All")
MagnitudeSummaryIntervalData[,c("Correct", "Incorrect", "SamplingType", "Model", "CastType")] <- NULL
names(MagnitudeSummaryIntervalData) <- c("SampleLength", "Tolerance", "CompleteLength", "GenLength", "EDF", "IntervalLength", "PropWithinTolerance")
write.csv(MagnitudeSummaryIntervalData, paste0(FiguresFP, "Supporting/MagntiudeDataInterval_SupportingDataSection4.csv"), row.names=FALSE) #Write out the data

### Write functions for plotting
DirectionPlot <- function(Direction){ 
  Direction$Prop <- Direction$PropCorrect
  Direction$Prop <- round_any(Direction$Prop*100, 10, floor) #Convert proportions to percentages and round down to nearest 10
  Direction$Prop <- paste0(Direction$Prop, "-", Direction$Prop+10) #Create label (e.g. "0-10" rather than just "0")
  Direction$Prop <- factor(Direction$Prop, levels=sapply(seq(0,90,10), function(x) paste0(x, "-", x+10))) #Convert to factor
  
  if(SamplingType=="Consecutive"){ #Create labels (adjusts based on consecutive or interval sampling)
    xlabel <- "Complete trend length (years)"
    ylabel <- "Sample trend length (years)"
    xbreaks <- (c(seq(4,MCL,4)))
    ybreaks <- (c(seq(3,MCL,3)))
    ex <- Direction$CompleteLength
    why <- Direction$NumYears
    aspectrat <- 1
  } else {
    xlabel <- "Number of years sampled"
    ylabel <- "Samples taken every x years"
    xbreaks <- (c(seq(3,(MCL-1),5)))
    ybreaks <- (c(seq(1,(MCL/2),3)))
    aspectrat <- 0.52
    ex <- Direction$NumYears
    why <- Direction$IntervalLength
  }
  
  ggplot(Direction, aes(x=ex,y=why))+
    geom_tile(aes(fill=Prop), colour = "grey40",size = 0.01)+ #Tile makes the nice squares
    ylab(ylabel)+
    xlab(xlabel)+
    scale_fill_manual(values = viridis(n=10), name=paste0("Percentage"), guide=guide_legend(reverse = TRUE, keywidth=unit(3, "mm"), keyheight = unit(3, "mm")), drop=FALSE)+ #This sets the legend as the same regardless of which colours appear (it always shows all options from 0-100)
    scale_x_continuous(breaks=xbreaks, expand = c(0,0.15))+
    scale_y_continuous(breaks=ybreaks, expand = c(0,0.15))+
    theme(aspect.ratio=aspectrat, 
          panel.background = element_blank(),
          panel.grid = element_blank(), 
          text = element_text(size=plotfontsize),
          panel.border = element_rect(size = 1, fill = NA),
          strip.background = element_blank(),
          strip.text = element_text(hjust = 0, size=plotfontsize),
          legend.title=element_text(size=plotfontsize),
          axis.text = element_text(size=plotfontsize, colour="black"),
          axis.ticks = element_line(colour="black"),
          legend.text = element_text(size=plotfontsize, colour="black"),
          legend.justification = "top")
}
MagnitudePlot <- function(Direction){
  MagnitudeSub <- c(0.01,0.025, 0.05, 0.1, 0.25, 0.5) #Subset to the tolerances we want to plot (I modelled more just out of interest)
  Direction <- Direction[Direction$Tolerance %in% MagnitudeSub,] #Subset to the tolerances we want to plot (I modelled more just out of interest)
  Direction$Tolerance <- paste0("± ", Direction$Tolerance) #Add a plusminus sign so labels are correct
  if(SamplingType=="Consecutive"){
    Direction$CompleteLength <- as.character(as.numeric(Direction$CompleteLength))
  }
  if(SamplingType=="Interval"){
    Direction$CompleteLength <- Direction$IntervalLength #Cheeky relabel of interval length to be complete length so that this function can be applied to both data (bit janky but it works, and labels ensure it's labelled as interval length in the actual plot)
  }
  Direction$Tolerance <- as.factor(Direction$Tolerance)
  if(SamplingType=="Consecutive"){ #Add labels to the years (consecutive)
    Direction$CompleteLength <- sapply(Direction$CompleteLength, function(x){
      if(x=="10"){
        paste0("a) ", x, " years")
      } else if(x=="20"){
        paste0("b) ", x, " years")
      } else if(x=="30"){
        paste0("c) ", x, " years")
      }
    })
    Direction$CompleteLength <- factor(Direction$CompleteLength, levels=c("a) 10 years","b) 20 years","c) 30 years"))
  } else {
    Direction$CompleteLength <- sapply(Direction$CompleteLength, function(x){ #Add labels to the years between samples (intervals)
      if(x==1){
        paste0("a) Samples every ", x, " yr")
      } else if(x==3){
        paste0("b) Samples every ", x, " yrs")
      } else if(x==6){
        paste0("c) Samples every ", x, " yrs")
      }
    })
    Direction$CompleteLength <- factor(Direction$CompleteLength, levels=c("a) Samples every 1 yr","b) Samples every 3 yrs","c) Samples every 6 yrs"))
  }
  if(SamplingType=="Consecutive"){ #Set breaks for plot
    breaksx <- seq(0,MCL,5)
    breaksy <- seq(0,100,10)
  } else {
    breaksx <- seq(3,15,2)
    breaksy <- seq(0,100,10) 
  }
  Direction$PropCorrect <- Direction$PropCorrect*100 #Change proportion correct to a percentage
  ggplot(Direction, aes(x=NumYears,y=PropCorrect, colour=Tolerance))+ #make a line plot!
    geom_line()+
    geom_point(shape=18, size=2)+
    facet_wrap(~CompleteLength)+
    ylab("Percentage within tolerance")+
    xlab("Number of years sampled")+
    scale_colour_manual(values = viridis(n=7), name="Tolerance", guide=guide_legend(reverse = TRUE, keywidth=unit(3, "mm"), keyheight = unit(3, "mm")), drop=FALSE)+
    scale_x_continuous(breaks=breaksx)+ #, expand = c(0,0)
    scale_y_continuous(breaks=breaksy)+ #, expand = c(0,0)
    theme(aspect.ratio=1, 
          panel.background = element_blank(),
          panel.grid = element_blank(), 
          text = element_text(size=plotfontsize),
          panel.border = element_rect(size = 1, fill = NA),
          strip.background = element_blank(),
          strip.text = element_text(hjust = 0),
          legend.title=element_text(size=plotfontsize),
          legend.key=element_blank(),
          legend.justification = "top")
}

modelfam <- "nb" #Adjust this if you used a different model fam in above runs

### Build and save direction plots (supplementary insig code)
for(SamplingType in c("Consecutive", "Interval")){ #Loop through consecutive and interval
  for(Casttype in c("All", "St")){ #Loop through all data ("All") and standardised data ("St", which checks for biases)
    if(SamplingType=="Consecutive"){ #Subset to relevant type
      Direction <- subset(DirectionSummary, CastType==Casttype)
    } else {
      Direction <- subset(DirectionSummaryInterval, CastType==Casttype)
    }
    
    DirectionAll <- subset(Direction, GenLength==0 & EDF==0) #Remove data subsetted by Generation Length or Trend Shape
    if(SamplingType=="Consecutive"){ #Set plot dimensions
      W <- 113
      H <- 77
    } else {
      W <- 124
      H <- 55
    }
    #Apply plot functions and save
    ggsave(paste0(FiguresFP,"Insig/Direction/Direction_", SamplingType, "_", Casttype, "_", modelfam, ".pdf"), DirectionPlot(DirectionAll), device="pdf", width = W, height = H, units = "mm") #Save as a pdf
  }
}

### Build and save magnitude plots (supplementary insig code)
for(SamplingType in c("Consecutive", "Interval")){ #Loop through consecutive and interval
  for(Casttype in c("All", "St")){ #Loop through all data ("All") and standardised data ("St", which checks for biases)
    if(SamplingType=="Consecutive"){ #Subset to relevant type
      Magnitude <- subset(MagnitudeSummary, CastType==Casttype)
    } else {
      Magnitude <- subset(MagnitudeSummaryInterval, CastType==Casttype)
    }
    MagnitudeAll <- subset(Magnitude, GenLength==0 & EDF==0) #First look at all data within subsetting
    
    if(SamplingType=="Consecutive"){
      MagnitudeAll <- MagnitudeAll[MagnitudeAll$CompleteLength %in% c(10,20,30),] #For consecutive, reduce to just a few complete lengths (otherwise too much data to plot)
    } else {
      MagnitudeAll <- MagnitudeAll[MagnitudeAll$IntervalLength %in% c(1,3,6),] #For interval, reduce to just a few interval lengths (otherwise too much data to plot)
      MagnitudeAll <- subset(MagnitudeAll, NumYears<16)
    }
    
    H <- 80
    W <- 180
    
    #Apply plot functions and save
    ggsave(paste0(FiguresFP,"Insig/Magnitude/Magnitude_", SamplingType, "_", Casttype, "_", modelfam, ".pdf"), MagnitudePlot(MagnitudeAll), device="pdf", width = W, height = H, units = "mm") #Save as image
  }
}
```
